# Development of photo-lenalidomide for cellular target identification

**DOI:** 10.1101/2021.07.12.452075

**Authors:** Zhi Lin, Yuka Amako, Farah Kabir, Hope A. Flaxman, Bogdan Budnik, Christina M. Woo

**Author notes:** These authors contributed equally to this work (Zhi Lin, Yuka Amako).

## Abstract

The thalidomide analog lenalidomide is a clinical therapeutic that alters the substrate engagement of cereblon (CRBN), a substrate receptor for the CRL4 E3 ubiquitin ligase. Here, we report the development of photo-lenalidomide, a lenalidomide probe with a photo-affinity label and enrichment handle, for target identification by chemical proteomics. After evaluating a series of lenalidomide analogs, we identified a specific amide linkage to lenalidomide that allowed for installation of the desired functionality, while preserving the substrate degradation profile, phenotypic anti-proliferative and immunomodulatory properties of lenalidomide. Photo-lenalidomide maintains these properties by enhancing binding interactions with the thalidomide-binding domain of CRBN, as revealed by binding site mapping and molecular modeling. Using photo-lenalidomide, we captured the known targets IKZF1 and CRBN from multiple myeloma MM.1S cells, and further identified a new target, eukaryotic translation initiation factor 3 subunit i (eIF3i), from HEK293T cells. eIF3i is directly labeled by photo-lenalidomide and forms a complex with CRBN in the presence of lenalidomide, but is itself not ubiquitylated or degraded. These data point to the potentially broader array of substrates induced by ligands to CRBN that may or may not be degraded, which can be revealed by the highly translatable application of photo-lenalidomide and chemical proteomics in additional biological settings.

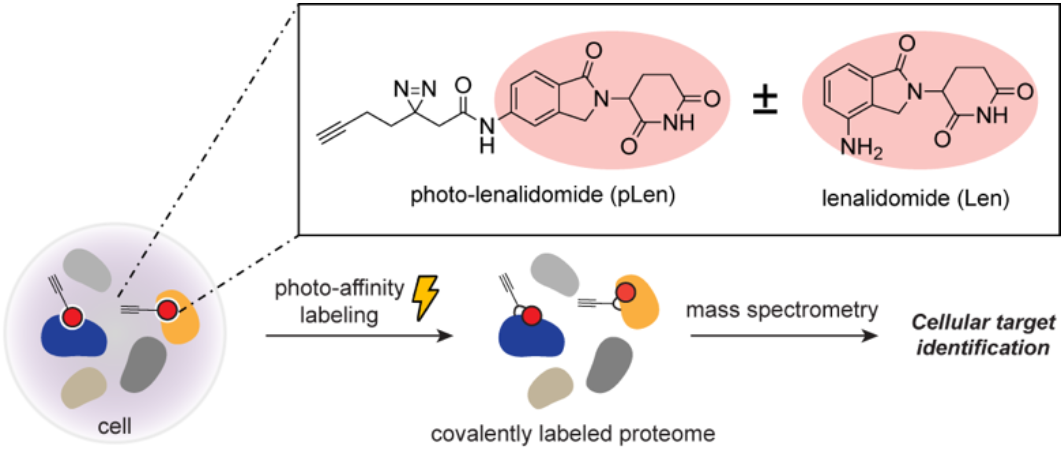

## INTRODUCTION

Lenalidomide is an analog of thalidomide used in treatment of multiple myeloma, del(5q) myelodysplastic syndrome, and additional hematopoietic malignancies. The therapeutic effects of lenalidomide arise in part via the engagement of cereblon (CRBN), a substrate receptor for the CRL4 E3 ubiquitin ligase complex (CRL4^CRBN^),^1^ which promotes ubiquitylation and degradation of protein substrates, including IKZF1, IKZF3, and CK1α.^2–4^ Lenalidomide and thalidomide additionally possess context-dependent anti-angiogenic, anti-inflammatory, immunostimulatory, and teratogenic effects.^5–6^ While the teratogenic effects of thalidomide have been traced to substrates such as SALL4^7–8^ and p63,^9^ the mechanistic targets underlying the immunomodulatory and anti-angiogenic effects remain unknown.^10–11^ Understanding the mechanistic targets engaged towards these effects is essential due to the growing use of thalidomide analogs in clinical practice and targeted protein degradation strategies.^12–13^

The discovery of substrates targeted by lenalidomide and analogs has typically relied on approaches that either report on the ubiquitylation and degradation of the substrate or selective co-immunoprecipitation with CRBN. Methods to discover substrates through their ubiquitylation and degradation include global proteomics, ubiquitin profiling, and plasmid library screening.^2–4, 7^ These mechanism-based approaches may overlook lower abundance substrates or those that are not degraded under the treatment conditions, and often have strict requirements on fractionation, sequencing or sample preparation to enhance coverage in each cell type of interest. Alternatively, in vitro co-immunoprecipitation of CRBN in the presence or absence of lenalidomide and analogs allows for identification of shifts in protein complexes that may include the targeted substrates,^14–16^ but requires additional studies to ensure a direct binding interaction between CRBN and the identified targets and may overlook more transient but functional binding interactions within a cell. A strategy to identify the direct targets of lenalidomide, independent of substrate degradation, would provide a distinct approach for target identification and thus accelerate mechanism of action studies and the development of next-generation therapeutics in the long-term.

The combination of photo-affinity labeling (PAL) with chemical proteomics allows for cellular target identification via the unbiased characterization of small molecule–protein interactions over a dynamic range of affinities.^17–21^ In a typical experiment, cells are treated with a chemical probe embedded with a PAL functional group (e.g., a diazirine) and an enrichment handle. The protein binding partners are captured on photolysis of the probe, followed by enrichment and analysis by mass spectrometry (MS). Proteins that are selectively enriched represent direct binding events with the chemical probe, which can yield new targets of small molecules in combination with competitive displacement. We have been characterizing diazirine chemistry for its ability to yield novel targets and binding site measurements of small molecules,^22^ with the objective of expediting the discovery of novel molecular glues, like rapamycin,^23^ sanglifehrin, or lenalidomide, and the protein complexes that underpin their biological activities. However, to date, the propensity of PAL to report on protein complexes mediated by a molecular glue within cells remains undetermined.

Here we report the development of photo-lenalidomide, a probe that enables identification of the cellular targets of lenalidomide by chemical proteomics (Figure 1a). We demonstrate that photo-lenalidomide preserves the phenotypic profile of lenalidomide and enables binding site mapping in vitro and target identification from protein complexes in cells. Evaluation of photo-lenalidomide in HEK293T cells revealed eukaryotic translation initiation factor 3 subunit i (eIF3i) as a novel target of the lenalidomide– CRBN complex that is not ubiquitylated or degraded. The ready translatability of photo-lenalidomide to different cellular contexts may reveal additional targets of lenalidomide that contribute to the diverse biological effects of lenalidomide and further demonstrate the utility of PAL-based chemical proteomics approaches to discover new targets of molecular glues.

**Figure 1.**
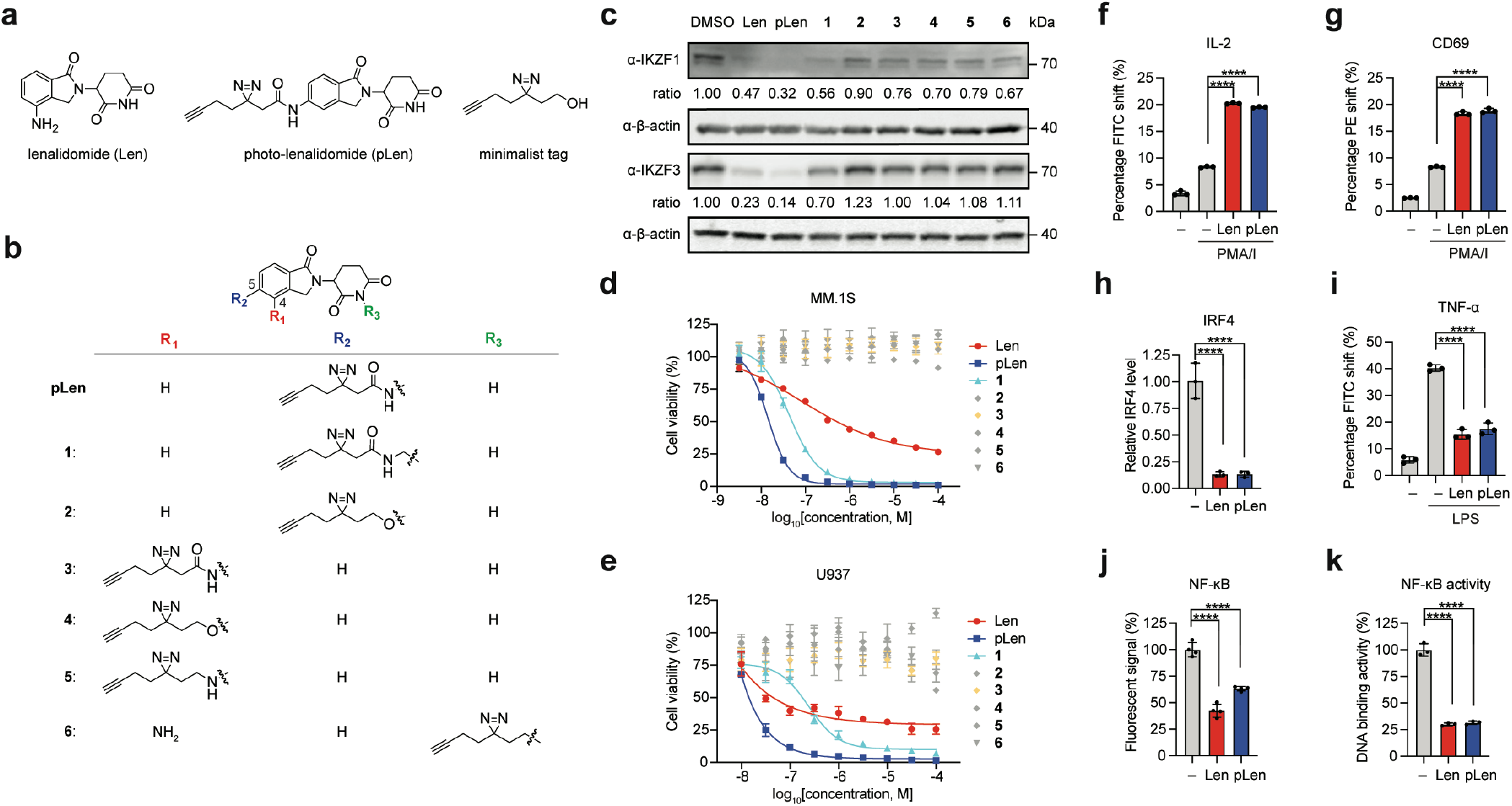
Development of photo-lenalidomide as a chemical proteomics probe for lenalidomide. (**a**) Structures of lenalidomide (Len), photo-lenalidomide (pLen), and the minimalist tag. (**b**) The structure of lenalidomide analogs (pLen, **1–6**) evaluated as probes for lenalidomide. (**c**) Western blot of substrates IKZF1 and IKZF3 after treatment of MM.1S cells with 10 *μ*M Len, pLen, or analogs **1–6** for 8 h. (**d, e**) Cell viability (MTT) assay of MM.1S (**d**) or U937 (**e**) cells after treatment with Len, pLen, or analogs **1– 6** for 5 d. (**f, g**) Percentage of interleukin-2 (IL-2) (**f**) and CD69 (**g**) positive Jurkat cells after treatment with 1 *μ*M Len or pLen, and stimulation with PMA/I for 48 h (total cell count = 100,000). (**h**) Levels of IRF4 mRNA in MM.1S cells after treatment with 10 *μ*M Len or pLen for 48 h. Data was normalized to GAPDH for each condition. (**i**) Levels of TNF-α in RAW264.7 cells after treatment with 150 *μ*M Len or pLen and stimulation with 1 ng/mL lipopolysaccharide (LPS) for 3 h. (**j**) Levels of a NF-κB luciferase reporter in HEK293T cells after treatment with 100 *μ*M Len or pLen and stimulation with TNF-α for 5 h. Luminescence was normalized to that of untreated cells. (**k**) Percent of nuclear NF-κB p65 subunit bound to DNA in MM.1S cells after treatment with 10 *μ*M Len or pLen for 48 h. Data was normalized to the untreated samples. All graphs are shown as mean ± s.d., n = 3 or 4. Statistical significance was analyzed in f–k using a 2-way Student’s t-test. *P < 0.05; **P< 0.005; ***P < 0.0005, ****P < 0.0001; n.s. not significant. Full immunoblot images are shown in Figure S10.

## RESULTS

### Development of photo-lenalidomide as a probe for lenalidomide

We initiated our efforts with the development of a photo-affinity probe of lenalidomide that preserves the native functionality of the parent compound. Lenalidomide is composed of an isoindolinone conjugated to a glutarimide, where the glutarimide forms contacts with the thalidomide-binding pocket of CRBN.^24–25^ We developed seven candidate photo-lenalidomide analogs bearing an alkyl diazirine alkyne, termed the “minimalist tag”,^19^ at positions 4 and 5 of the isoindolinone, or via derivatization of the glutarimide (Figure 1b, Scheme S1).

To identify the probe that best recapitulated the biological activities of lenalidomide, the lenalidomide analogs were initially assessed for their ability to induce degradation of known lenalidomide targets IKZF1 and IKZF3.^2–3^ The multiple myeloma cell line MM.1S was treated with 10 *μ*M lenalidomide (Len) or analogs **1–6** for 8 h, and levels of IKZF1 and IKZF3 were visualized by Western blot (Figure 1c). We found that one of the seven compounds depleted IKZF1 and IKZF3 levels comparably to lenalidomide, which we henceforth termed photo-lenalidomide (pLen). Depletion of IKZF1 and IKZF3 by both lenalidomide and photo-lenalidomide occurred over a similar dose-response range (0.1–10 *μ*M) and was attenuated by the neddylation inhibitor MLN4924 (Figure S1). Photo-lenalidomide is functionalized at the 5-position of the isoindolinone by an amide linkage to the minimalist tag. Insertion of a single methylene unit as in analog **1** yielded moderate degradation efficiency for IKZF1 and IKZF3 (relative ratios = 0.56, 0.70, respectively), while the ether analog **2** displayed no effect on the levels of these target proteins (Figure 1c). Analogs **3–5**, which are substituted with the minimalist tag at the 4-position of the isoindolinone, were similarly inactive in degradation of IKZF1 and IKZF3. Likewise, functionalization of the glutarimide in analog **6** inactivated degradation of these target proteins, presumably by blocking engagement of CRBN. Notably, photo-lenalidomide additionally depleted the lenalidomide substrate CK1α and both photo-lenalidomide and analog **1** depleted GSPT1, a common target of lenalidomide analogs,^14, 26^ at 10 *μ*M in MM.1S cells (Figure S2).

### Phenotypic characterization of photo-lenalidomide

Lenalidomide possesses cell-type specific anti-proliferative, anti-inflammatory, anti-angiogenesis, and immuno-stimulatory properties.^5–6^ We therefore evaluated the ability of photo-lenalidomide to recapitulate these activities of lenalidomide. First, both lenalidomide and photo-lenalidomide displayed dosedependent anti-proliferative effects against MM.1S cells (GI_50_ = 59.2 nM, 27.2 nM, respectively) and U937 leukemia cells (GI_50_ = 15.3 nM, 15.2 nM, respectively) (Figure 1d, 1e). Analog **1** additionally displayed anti-proliferative effects (GI_50_ MM.1S = 42.9 nM, U937 = 116.2 nM), whereas the five additional analogs displayed no anti-proliferative activity up to 100 *μ*M in MM.1S cells, consistent with their limited effects on IKZF1 and IKZF3 levels. The higher potency observed with photo-lenalidomide and analog **1** as compared to lenalidomide may be attributable to their expanded substrate scope (e.g., GPST1, Figure S2). As expected, lenalidomide-insensitive cell lines,^2^ such as MOLM13, Jurkat, K562, and mouse macrophage RAW264.7 cells were likewise not sensitive to photo-lenalidomide treatment (Figure S3). Therefore, the anti-proliferative effects of photo-lenalidomide correlate with those previously observed for lenalidomide.

Next, the co-immunostimulatory properties of the seven probes were evaluated. Lenalidomide promotes T-cell activation on co-stimulation with phorbol 12-myristate 13-acetate and ionomycin (PMA/I), resulting in elevated IL-2 and CD69 expression.^27^ Co-stimulation of Jurkat cells with lenalidomide or photo-lenalidomide elevated expression of IL-2 and CD69 by 3-fold compared to DMSO control (Figure 1f, 1g). Notably, all other lenalidomide analogs also increased cytokine production apart from the analog **6**, which implies that the co-stimulatory effect may be dependent on the engagement of CRBN (Figure S4a, S4b).

Separately, we compared the immunosuppressive and anti-inflammatory properties of lenalidomide and photo-lenalidomide. Reduced IRF4 expression is a downstream effect of lenalidomide treatment in multiple myeloma.^2^ We observed equivalent suppression of IRF4 mRNA in MM. 1S cells upon treatment with lenalidomide or photo-lenalidomide (Figure 1h). Lenalidomide additionally possesses immunosuppressive effects through TNF-α suppression and NF-κB down-regulation.^28–29^ Likewise, photo-lenalidomide significantly decreased TNF-α at the protein and mRNA levels in lipopolysaccharide (LPS)-stimulated mouse macrophage RAW264.7 cells (Figure 1i, S4c–d). In addition, both compounds decreased NF-κB signaling as measured by a luciferase NF-κB reporter gene system with exogenous TNF-α stimulation (Figure 1j). The effect on NF-κB signaling by lenalidomide and photo-lenalidomide was also observed in inhibition of the DNA binding activity of the endogenous NF-κB p65 subunit (Figure 1k).^28^ Collectively, these phenotypic assays indicate that the activity profile of photo-lenalidomide is similar to lenalidomide, suggesting that photo-lenalidomide recapitulates these biological effects of lenalidomide and is a suitable probe to reveal targets of lenalidomide that underlie their mechanisms.

### Binding site mapping of CRBN thalidomide-binding domain with photo-lenalidomide

Intrigued by the selective structure–activity relationship we observed in the evaluation of lenalidomide analogs, we next sought to analyze the binding mode of photo-lenalidomide to the thalidomide-binding domain of CRBN (CULT domain, aa318–442, Figure 2a, S5a),^30^ in comparison to the structurally similar analog **1** which differs from photo-lenalidomide by a single methylene. Lenalidomide binds to the CULT domain through the glutarimide, where the binding pocket is composed of three tryptophan residues (Trp380, Trp386, Trp400) and the isoindolinone is exposed to the surface of the protein with one face interacting with Asn351, Pro352 and His353.^24–25^ To gain structural insight into the interaction between CRBN and photo-lenalidomide and analog **1**, the CULT domain and each compound were incubated for 30 min and photo-crosslinking was induced upon photo-irradiation. The labeled domain was subsequently conjugated with an isotopic code using a cleavable biotin picolyl azide via copper-catalyzed azide alkyne cycloaddition (CuAAC) to enhance confidence in the binding site assignment.^21^ After chymotryptic digestion, the peptides were analyzed by mass spectrometry (MS). The spectra obtained were searched by Sequest HT against a database of semi-chymotryptic peptides of CULT and common contaminant proteins allowing for one modification of the corresponding probe on any amino acid. Tandem mass spectrometry (MS2) spectra assigned to peptides conjugated by a probe were manually validated by evaluating the isotopic coding in the MS1 precursor spectra. Photo-lenalidomide was conjugated with 5 unique peptides across 28 PSMs, which encompassed residues 347–363 of CULT (Table S1, S2). The photo-lenalidomide modification site was primarily localized to His353, which is found in a loop facing one side of the isoindolinone of lenalidomide (Figure 2b, S5b).^25^ The result of binding site mapping suggests that photo-lenalidomide binds to the tri-tryptophan pocket with a similar conformation as lenalidomide with the diazirine projecting up towards the flexible loop. By contrast, analog **1** labeled two regions: residues 350–363 and residues 369–384 (Figure 2b, Fig S5c, Table S3, S4). The first region corresponds to the same loop region labeled by photo-lenalidomide, but the second region represents another loop on the opposing face of CULT (Figure S5b). The multisite labeling with analog **1** may reflect an increased flexibility of the minimalist tag imparted by the methylene between the isoindolinone and the amide linkage, which may contribute to the differential activity observed between photo-lenalidomide and analog **1** in cells.

**Figure 2.**
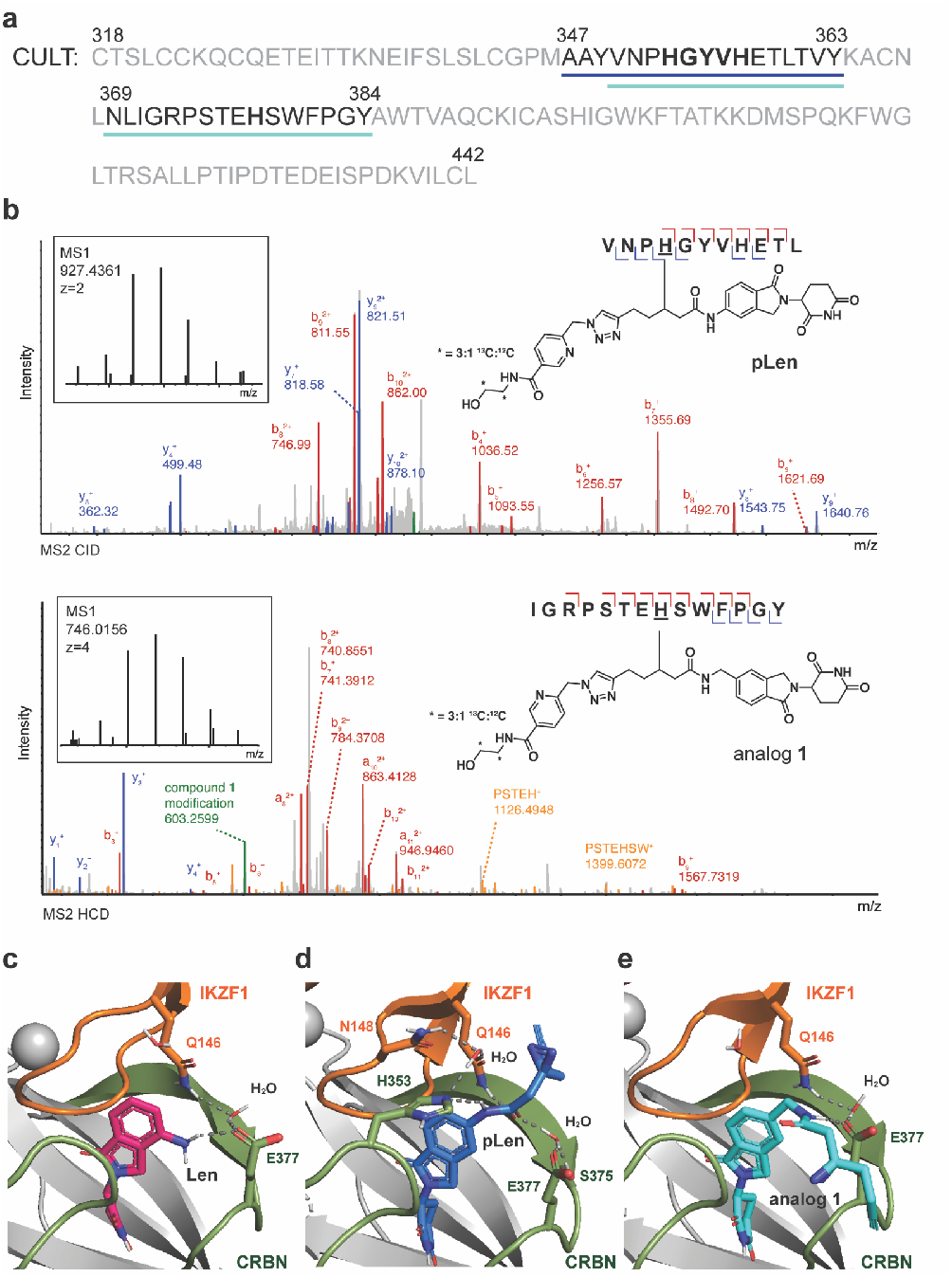
Evaluation of ligand interactions of photo-lenalidomide and analogs **1**. (**a**) Amino acid sequence of the CULT domain. Numbers indicate those of the corresponding amino acid residues in CRBN. Blue underline indicates residues identified as photo-lenalidomide-conjugated peptides and cyan underline indicates those of analog **1**-conjugated peptides. Bold indicates labeled amino acid residues. (**b**) Representative assignment of a conjugated peptide from CULT with pLen (top) and analog **1**(bottom). (**c–e**) Docked structure of CRBN-Len-IKZF1 ZF2 complex (**c**), CRBN-pLen-IKZF1 ZF2 complex (**d**) and CRBN-analog **1**-IKZF1 ZF2 complex (**e**) adapted from PDB 6H0F. The conjugated peptides of CRBN are colored green and the rest of residues is colored gray. IKZF1 ZF2 is colored orange.

With a map of the binding interactions between photo-lenalidomide and analog **1** and the CULT domain in hand, we next analyzed the structural implications of these interactions on the ternary complex with CRBN and IKZF1 by molecular modeling. Lenalidomide, photo-lenalidomide and analog **1** were adapted into its ligand pomalidomide in the crystal structure of DDB1-CRBN-pomalidomide-IKZF1 (ZF2) complex (PDB 6H0F)^31^ and docked to the complex using Molecular Operating Environment (MOE). The docking results were scored with the GBVI/WSA dG scoring function and the ligand interactions of the docked structure with the lowest score were analyzed. The docking analysis of lenalidomide afforded a structure with a similar conformation to pomalidomide, where the amine on the isoindolinone maintains a hydrogen bond network with Glu377 and Gln146 residues and a water molecule (Figure 2c). Docking with photo-lenalidomide revealed that it forms an extensive hydrogen bond network between CRBN and IKZF1. Specifically, the amide linker forms hydrogen bonding interactions directly with Gln146 and His353, and with Ser375 and Glu377 through a water molecule (Figure 2d). This hydrogen bond network may strengthen the binding of photo-lenalidomide to CRBN and its substrates, which may lead to the slight enhancement in degradation of IKZF1, IKZF3, and CK1α, and the recruitment of additional substrates (e.g. GSPT1). By contrast, the docked structure of the analog **1** showed a hydrogen bond network between the amide-NH and Glu377 and Gln146 residues and a water molecule (Figure 2e). Molecular modeling of the structurally related analog **3** was additionally performed to show it interacted solely with Gln146, which may explain its limited degradation profile (Figure S6). In summary, these data and the hydrogen bond patterns in the resulting models correlates with the substrate degradation profiles observed in cells.

### Mapping lenalidomide targets by chemical proteomics in live cells

We next evaluated the ability of photo-lenalidomide to label CRBN and IKZF1 in the ternary complex in live cells. In initial studies, HEK293T cells stably expressing Flag-CRBN (HEK293T-CRBN)^32^ were transfected with IKZF1 bearing a 4-amino acid EPEA tag (IKZF1-EPEA) and were treated with 50*μ*M of photo-lenalidomide with or without competition with 150 *μ*M of lenalidomide for 1 h prior to photoirradiation (120 sec). Treatment with 50 *μ*M of the minimalist tag was performed alongside as a control. The target proteins Flag-CRBN or IKZF1-EPEA were immunoprecipitated and the extent of photolenalidomide labeling was visualized by the conjugation of AlexaFluor^®^ 488 azide (AF488) via CuAAC and in-gel fluorescent imaging. We were pleased to observe selective labeling of both CRBN and IKZF1 by photo-lenalidomide in an UV-dependent manner (Figure 3a, S7).

**Figure 3.**
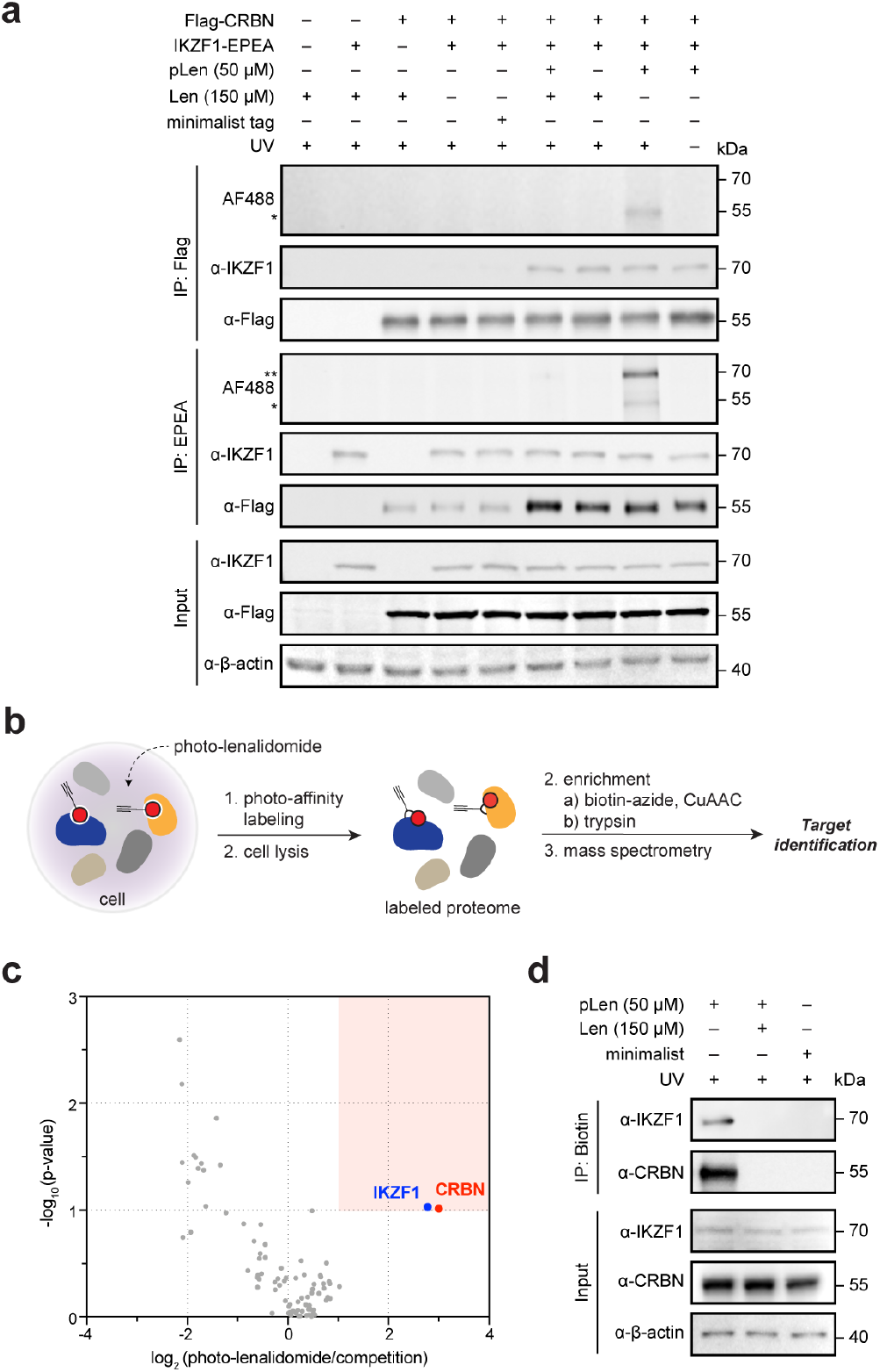
Evaluation of photo-lenalidomide as a chemical proteomics probe in cells. (**a**) In-gel fluorescence imaging and Western blot to visualize photo-affinity labeling of Flag-CRBN or IKZF1-EPEA after immunoprecipitation from treated HEK293T-CRBN cells transfected with IKZF1-EPEA. The asterisk (*) denotes CRBN band and the double asterisks (**) denotes IKZF1 band. (**b**) Schematic of target identification workflow from cells using photo-lenalidomide. (**c**) Volcano plot of lenalidomide targets identified by TMT quantitative proteomics in MM.1S cells treated with 50 *μ*M photo-lenalidomide with or without 150 *μ*M of lenalidomide (competition) for 1 h (n = 4). The significantly enriched region of proteins is shaded in red. (**d**) Western blot of IKZF1 and CRBN following treatment of MM.1S cells with the indicated compound, photo-irradiation, and enrichment. Full immunoblot images are shown in Figure S11.

Labeling of both target proteins was competitively inhibited by co-treatment with lenalidomide, and no labeling was observed in the presence of the minimalist tag (Figure 3a). Additionally, IKZF1 and CRBN were selectively co-immunoprecipitated in the presence of lenalidomide or photo-lenalidomide, which suggests both compounds form similar ternary complexes in cells. Importantly, both members of the ternary complex were labeled by photo-lenalidomide. Labeled CRBN was also observed in the sample immunoprecipitated by the EPEA antibody.

We next sought to utilize photo-lenalidomide to enrich and map lenalidomide targets in live cells using quantitative proteomics (Figure 3b). MM.1S cells were treated with 50 *μ*M photo-lenalidomide with or without lenalidomide competition (150 *μ*M) for 1 h, and photo-irradiated to capture cellular binding partners. After lysis, the labeled proteins were tagged with a biotin-azide probe via CuAAC, enriched on streptavidin-agarose beads, and the enriched proteins were digested with trypsin. The recovered peptides from four biological replicates were then analyzed by quantitative proteomics (Figure 3b, 3c, Table S5).^20^ As expected, CRBN and IKZF1 were significantly enriched binding partners of photo-lenalidomide and selectively competed off by co-treatment with lenalidomide [enrichment ratio(photo-lenalidomide/competition) = 8.01, 6.86, respectively]. The selective enrichment of these target proteins by photo-lenalidomide, but not on competition with lenalidomide, was further validated by Western blot (Figure 3d). Taken together, these data indicate that photo-lenalidomide is a suitable probe that can reveal lenalidomide targets within protein complexes directly from cells.

### Photo-lenalidomide reveals that eIF3i forms a complex with lenalidomide and CRBN

While evaluating photo-lenalidomide via chemical proteomics, we performed a similar competition experiment in HEK293T-CRBN cells in the absence of IKZF1-EPEA transfection. HEK293T-CRBN cells were treated with 50 *μ*M photo-lenalidomide with or without lenalidomide competition for 1 h before photo-irradiation, enrichment, and evaluation by quantitative proteomics. As expected, CRBN was enriched, though the enrichment ratio was reduced relative to that observed in MM.1S cells [Figure 4a, Table S6, enrichment ratio(photo-lenalidomide/competition) = 1.34, p-value = 0.002]. Interestingly, eukaryotic translation initiation factor 3 subunit i (eIF3i), a protein component of the eIF3 complex, was one of the most significantly enriched proteins [enrichment ratio (photo-lenalidomide/competition) = 3.13, p-value = 0.004]. The selective labeling and enrichment of CRBN and eIF3i by direct interaction with photo-lenalidomide was validated by Western blot (Figure 4b).

**Figure 4.**
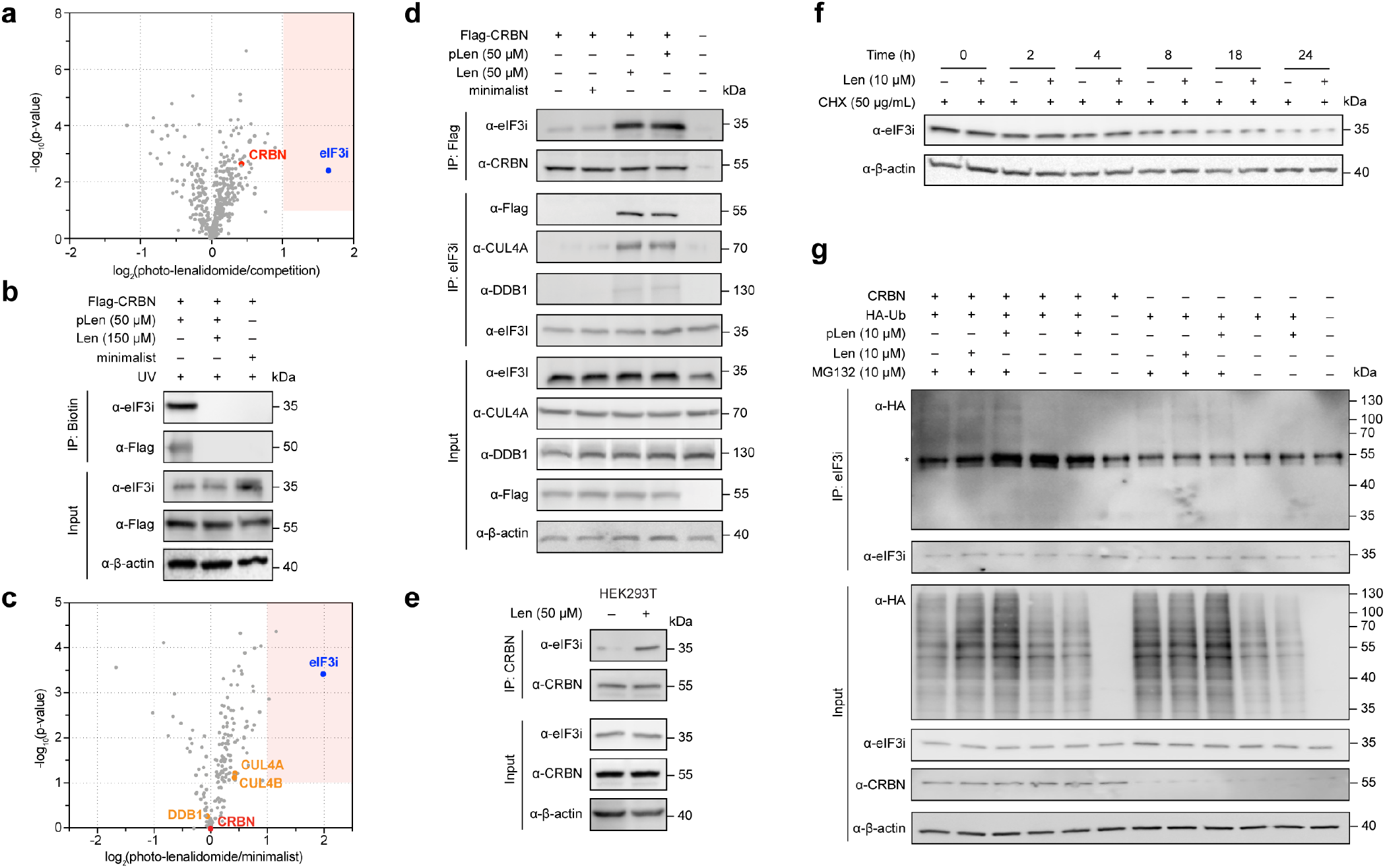
Discovery of the CRBN-lenalidomide-eIF3i ternary complex by chemical proteomics. (**a**) Volcano plot of the lenalidomide interactome identified by TMT quantitative proteomics in HEK293T-CRBN cells treated with 50 *μ*M photo-lenalidomide with or without 150 *μ*M of the parent compound for 1 h (n = 4). The significantly enriched region of proteins is shaded in red. (**b**) Western blot of eIF3i and Flag-CRBN following treatment with the indicated compound, photo-irradiation, and streptavidin enrichment from HEK293T-CRBN cells. (**c**) Volcano plot of proteins co-immunoprecipitated by Flag-CRBN identified by TMT quantitative proteomics in HEK293T-CRBN cells in the presence of 50 *μ*M pLen or the minimalist tag (n = 4). (**d**) Co-immunoprecipitation of eIF3i and Flag-CRBN with or without 50 *μ*M lenalidomide or photo-lenalidomide from HEK293T-CRBN cells. (**e**) Co-immunoprecipitation of eIF3i in HEK293T cells by endogenous CRBN with or without 50 *μ*M lenalidomide. (**f**) Cycloheximide (CHX) chase experiment in HEK293T-CRBN cells. Cells were treated with or without 10 *μ*M Len for 1 h before 50 *μ*g/mL CHX was added. Cells were collected at the indicated time and analyzed by immunoblotting. (**g**) In vivo ubiquitylation assay in HEK293T or HEK293T CRBN-KD cells. Cells transfected with HA-Ub were treated with 10 *μ*M Len or pLen for 1 h before 10 *μ*M MG132 was added. After 6 h incubation, cells were collected, immunoprecipitated with eIF3i antibody, and analyzed by immunoblotting. Input represents whole cell lysate. The asterisk (*) denotes a nonspecific band of IgG HC of eIF3i antibody. Full immunoblot images are shown in Figure S12-13.

Intrigued, we subsequently performed Flag-CRBN co-immunoprecipitation to evaluate whether eIF3i forms a ternary complex with photo-lenalidomide and CRBN. HEK293T-CRBN cells were incubated with photo-lenalidomide (50 *μ*M) or the minimalist tag as a negative control for 1 h. After lysis, the target proteins interacting with Flag-CRBN were co-immunoprecipitated with anti-Flag magnetic beads. After elution by Flag peptide, captured proteins were digested and analyzed by mass spectrometry (Figure 4c, Table S7). eIF3i was one of the most significantly enriched proteins [ratio(photo-lenalidomide/minimalist) = 3.95, p-value = 0.0004], with core components of the cullin-RING-based E3 ubiquitin-protein ligase complex, such as DNA damage-binding protein 1 (DDB1) and Cullin-4A/Cullin-4B (CUL4A/B), also observed.

Co-immunoprecipitation of eIF3i with CRBN in the presence of photo-lenalidomide was additionally validated by Western blot and the reciprocal immunoprecipitation of CRBN and eIF3i, which was extended to lenalidomide (Figure 4d). Additional subunits of CRL4^CRBN^ E3 ligase complex, including DDB1 and CUL4A, were also immunoprecipitated through eIF3i in the presence of photo-lenalidomide and lenalidomide, which suggests eIF3i is recruited to the CRL4^CRBN^ complex by photo-lenalidomide or lenalidomide (Figure 4d).

To further examine the generalizability of the interaction between CRBN and eIF3i, we performed co-immunoprecipitation experiments in wild-type HEK293T cells and found that eIF3i is co-immunoprecipitated by CRBN in the presence of lenalidomide (Figure 4e). However, co-immunoprecipitation of eIF3i by CRBN was not promoted by lenalidomide in MM.1S cells (Figure S8). This implies that the stabilization of eIF3i in a complex with CRBN by lenalidomide is cell-type specific.

### eIF3i is not ubiquitylated or degraded when engaged by lenalidomide in complex with CRBN

We next investigated whether the ternary complex formation between CRBN and eIF3i mediated by lenalidomide would lead to eIF3i degradation. HEK293T-CRBN cells were dosed with 10 *μ*M lenalidomide or photo-lenalidomide for 48 h with or without the neddylation inhibitor MLN4924. To our surprise, eIF3i levels were stable when under these conditions (Figure S9a). To further test the effect of lenalidomide on the half-life of eIF3i, we performed a time course chase experiment with the protein synthesis inhibitor cycloheximide (CHX) for up to 24 h in HEK293T cells (Figure 4f). Following treatment with 50 *μ*g/mL CHX, we found that eIF3i protein levels were unaltered with lenalidomide treatment.

To evaluate the possibility that eIF3i is ubiquitylated but not degraded, we further evaluated the ability of lenalidomide or photo-lenalidomide to ubiquitylate eIF3i. HEK293T cells or HEK293T with CRBN knockdown (KD) cells were transfected with HA-ubiquitin and pre-incubated with lenalidomide or photo-lenalidomide before the addition of the proteasome inhibitor MG132 to facilitate accumulation of ubiquitylated proteins. After lysis, protein lysates were immunoprecipitated by anti-eIF3i antibody for Western blot analysis. In both HEK293T cells and HEK293T CRBN KD cells, ubiquitylation levels on eIF3i did not change in a lenalidomide or photo-lenalidomide dependent manner (Figure 4g). These data indicate that neither lenalidomide treatment nor CRBN engagement is contributing to the ubiquitylation of eIF3i. Instead, we observed a minor stabilization of eIF3i in the presence of lenalidomide or photo-lenalidomide by thermal shift assay in cells (Figure S9b). These data indicate that eIF3i is one of the first identified targets of lenalidomide that is recruited to, but not ubiquitylated or degraded by, the CRL4^CRBN^ complex.

## DISCUSSION

Here we report the development of photo-lenalidomide as a probe for lenalidomide, which is a member of a prominent and growing class of therapeutics that engage CRBN. We examined a total of seven candidates to identify photo-lenalidomide, which has the minimalist tag projecting from the 5-position of the isoindolinone through an amide linkage. Photo-lenalidomide maintains the strong degradation activity against IKZF1 and IKZF3 observed with lenalidomide, and gains activity against CK1α and GSPT1, which may be rationalized by an increased hydrogen bond network predicted by binding site mapping and molecular modeling. Although these additional substrates likely enhance the anti-proliferative activity of photo-lenalidomide relative to lenalidomide in MM.1S cells, the broader phenotype profiles between the two compounds are comparable. As has been observed in other studies with lenalidomide analogs, including the development of bifunctional degraders,^33^ we found that preserving efficient ligand interactions was highly sensitive to both the tag position and the linkage. Gratifyingly, photo-lenalidomide selectively enriched both CRBN and IKZF1 from live cells, which demonstrates that PAL can efficiently report on protein complexes. Taken together, photo-lenalidomide recapitulates the phenotypic properties of lenalidomide and is a versatile probe that may be readily applied to reveal the direct protein interactions of lenalidomide within multi-protein complexes from cells, which will provide a more holistic and unbiased view of protein targets.

The chemoproteomics strategy using photo-lenalidomide to discover targets of lenalidomide as demonstrated here may accelerate the discovery of new small molecule-protein interactions, which may be expanded to additional lenalidomide derivatives in a variety of biological systems. Furthermore, photo-lenalidomide may enable additional experimental applications, including elucidation of CRBN-independent mechanisms^10, 34^ live cell labeling for confirming sub-cellular localization,^35^ and cellular target engagement measurements. Extension of the method to incorporate principles derived from enantioprobes^36^ may additionally enable measurement of stereoselective interactions of lenalidomide in the global proteome. Nonetheless, several limitations using photo-lenalidomide may preclude its application to all biological systems. For example, this method cannot be used with biological systems that cannot be photo-affinity labeled. Additionally, the strategy will not report on target proteins that lack an appropriate amino acid environment due to insufficient labeling of the probe.^22^ Finally, the proteome enriched by photo-lenalidomide requires validation by competition with lenalidomide, as photo-lenalidomide and PAL can potentially label non-specific targets. Evaluation of whether an identified target protein is part of a ternary complex with CRBN, or it is CRBN-independent additionally requires followup studies.

Photo-lenalidomide facilitated the direct identification of lenalidomide targets by chemoproteomics, which resulted in the discovery of eIF3i, a member of the eIF3 complex, as a novel substrate of the CRL4^CRBN^ complex recruited by lenalidomide. Surprisingly, eIF3i is not ubiquitylated or degraded by the CRL4^CRBN^ complex after recruitment. As lenalidomide is reported to stabilize CRBN itself,^37–38^ and alter interactions of CRBN with molecular chaperones MCT1-CD147^15^ and HSP90^39^, in addition to altering substrate degradation, there may be degradation-independent mechanisms of lenalidomide that arises from stabilization of substrates, analogous to other molecular glues (e.g., rapamycin, FK506, cyclosporin, sanglifehrin).^40^ Indeed, some bifunctional compounds that ligand to CRBN (e.g., PROTACs) have been observed to engage the target protein in a ternary complex, but the target protein is not degraded.^41–43^ The findings herein imply that the concept of target engagement without degradation may extend more broadly to lenalidomide and other CRBN ligands.

The discovery of eIF3i as a target of lenalidomide, as reported by photo-lenalidomide, also points to new structural and mechanistic directions in studies with lenalidomide. Structurally, eIF3i is a helical WD40 protein^44^ that does not have a known recognition motif targeted by lenalidomide, such as the C2H2 zinc finger motif.^31^ eIF3i may not be ubiquitylated and degraded due to inefficient transfer of ubiquitin by the CRL4^CRBN^ complex to a lysine on eIF3i, or the presence of additional members of the eIF3 complex may intercept the ubiquitylation event. Additional evaluation of the CRBN-lenalidomide-eIF3i complex will elucidate contributions that eIF3i recruitment may have on the biological outcomes of lenalidomide, and may reveal additional motifs that are engaged by the lenalidomide-CRBN complex. Further application of photo-lenalidomide across biological systems to uncover targets of lenalidomide, which may or may not be degraded, may help connect additional therapeutic outcomes of lenalidomide with their molecular targets.

## SUPPORTING INFORMATION

For SI figures and tables, a full description of the methods, and chemical characterization, see the Supporting Information.

## AUTHOR INFORMATION

**Corresponding Author** *

## Notes

The authors declare no competing financial interest.

## ACKNOWLEDGMENT

We thank D. H. Ramirez, Y. Ge and B. Yang for insightful discussions; N. L. Tran for support with recombinant protein preparation, Z. T. Niziolek and J. A. Nelson from Harvard University Bauer Core Facility, S. A. Trauger, R. A. Robinson and J. Wang from Harvard University Mass Spectrometry and Proteomics Resource Laboratory, and E. Cronin-Furman with Olympus for technical support.

## REFERENCES

1. Ito, T.; Ando, H.; Suzuki, T.; Ogura, T.; Hotta, K.; Imamura, Y.; Yamaguchi, Y.; Handa, H., Identification of a Primary Target of Thalidomide Teratogenicity. Science 2010, 327 (5971), 1345–1350.

2. Krönke, J.; Udeshi, N. D.; Narla, A.; Grauman, P.; Hurst, S. N.; McConkey, M.; Svinkina, T.; Heckl, D.; Comer, E.; Li, X.; Ciarlo, C.; Hartman, E.; Munshi, N.; Schenone, M.; Schreiber, S. L.; Carr, S. A.; Ebert, B. L., Lenalidomide Causes Selective Degradation of IKZF1 and IKZF3 in Multiple Myeloma Cells. Science 2014, 343 (6168), 301–305.

3. Lu, G.; Middleton, R. E.; Sun, H.; Naniong, M.; Ott, C. J.; Mitsiades, C. S.; Wong, K.-K.; Bradner, J. E.; Kaelin, W. G., The Myeloma Drug Lenalidomide Promotes the Cereblon-Dependent Destruction of Ikaros Proteins. Science 2014, 343 (6168), 305–309.

4. Kronke, J.; Fink, E. C.; Hollenbach, P. W.; MacBeth, K. J.; Hurst, S. N.; Udeshi, N. D.; Chamberlain, P. P.; Mani, D. R.; Man, H. W.; Gandhi, A. K.; Svinkina, T.; Schneider, R. K.; McConkey, M.; Jaras, M.; Griffiths, E.; Wetzler, M.; Bullinger, L.; Cathers, B. E.; Carr, S. A.; Chopra, R.; Ebert, B. L., Lenalidomide induces ubiquitination and degradation of CK1[agr] in del(5q) MDS. Nature 2015, 523 (7559), 183–188.

5. Rehman, W.; Arfons, L. M.; Lazarus, H. M., The Rise, Fall and Subsequent Triumph of Thalidomide: Lessons Learned in Drug Development. Ther Adv Hematology 2011, 2 (5), 291–308.

6. Corral, L. G.; Kaplan, G., Immunomodulation by thalidomide and thalidomide analogues. Ann Rheum Dis 1999, 58 Suppl 1, 1107–13.

7. Donovan, K. A.; An, J.; Nowak, R. P.; Yuan, J. C.; Fink, E. C.; Berry, B. C.; Ebert, B. L.; Fischer, E. S., Thalidomide promotes degradation of SALL4, a transcription factor implicated in Duane Radial Ray syndrome. eLife 2018, 7, e38430.

8. Matyskiela, M. E.; Couto, S.; Zheng, X.; Lu, G.; Hui, J.; Stamp, K.; Drew, C.; Ren, Y.; Wang, M.; Carpenter, A.; Lee, C. W.; Clayton, T.; Fang, W.; Lu, C. C.; Riley, M.; Abdubek, P.; Blease, K.; Hartke, J.; Kumar, G.; Vessey, R.; Rolfe, M.; Hamann, L. G.; Chamberlain, P. P., SALL4 mediates teratogenicity as a thalidomide-dependent cereblon substrate. Nat Chem Biol 2018, 14 (10), 981–987.

9. Asatsuma-Okumura, T.; Ando, H.; De Simone, M.; Yamamoto, J.; Sato, T.; Shimizu, N.; Asakawa, K.; Yamaguchi, Y.; Ito, T.; Guerrini, L.; Handa, H., p63 is a cereblon substrate involved in thalidomide teratogenicity. Nat Chem Biol 2019, 15 (11), 1077–1084.

10. Gemechu, Y.; Millrine, D.; Hashimoto, S.; Prakash, J.; Sanchenkova, K.; Metwally, H.; Gyanu, P.; Kang, S.; Kishimoto, T., Humanized cereblon mice revealed two distinct therapeutic pathways of immunomodulatory drugs. Proc Natl Acad Sci U S A 2018, 115 (46), 11802–11807.

11. Lu, L.; Payvandi, F.; Wu, L.; Zhang, L. H.; Hariri, R. J.; Man, H. W.; Chen, R. S.; Muller, G. W.; Hughes, C. C.; Stirling, D. I.; Schafer, P. H.; Bartlett, J. B., The anti-cancer drug lenalidomide inhibits angiogenesis and metastasis via multiple inhibitory effects on endothelial cell function in normoxic and hypoxic conditions. Microvasc Res 2009, 77 (2), 78–86.

12. Winter, G. E.; Buckley, D. L.; Paulk, J.; Roberts, J. M.; Souza, A.; Dhe-Paganon, S.; Bradner, J. E., Phthalimide conjugation as a strategy for in vivo target protein degradation. Science 2015, 348 (6241), 1376–1381.

13. Lu, J.; Qian, Y.; Altieri, M.; Dong, H.; Wang, J.; Raina, K.; Hines, J.; Winkler, James D.; Crew, Andrew P.; Coleman, K.; Crews, Craig M., Hijacking the E3 Ubiquitin Ligase Cereblon to Efficiently Target BRD4. Chem Biol 2015, 22 (6), 755–763.

14. Matyskiela, M. E.; Lu, G.; Ito, T.; Pagarigan, B.; Lu, C. C.; Miller, K.; Fang, W.; Wang, N. Y.; Nguyen, D.; Houston, J.; Carmel, G.; Tran, T.; Riley, M.; Nosaka, L.; Lander, G. C.; Gaidarova, S.; Xu, S.; Ruchelman, A. L.; Handa, H.; Carmichael, J.; Daniel, T. O.; Cathers, B. E.; Lopez-Girona, A.; Chamberlain, P. P., A novel cereblon modulator recruits GSPT1 to the CRL4(CRBN) ubiquitin ligase. Nature 2016, 535 (7611), 252–7.

15. Eichner, R.; Heider, M.; Fernandez-Saiz, V.; van Bebber, F.; Garz, A. K.; Lemeer, S.; Rudelius, M.; Targosz, B. S.; Jacobs, L.; Knorn, A. M.; Slawska, J.; Platzbecker, U.; Germing, U.; Langer, C.; Knop, S.; Einsele, H.; Peschel, C.; Haass, C.; Keller, U.; Schmid, B.; Gotze, K. S.; Kuster, B.; Bassermann, F., Immunomodulatory drugs disrupt the cereblon-CD147-MCT1 axis to exert antitumor activity and teratogenicity. Nat Med 2016, 22 (7), 735–43.

16. Zhu, Y. X.; Braggio, E.; Shi, C. X.; Kortuem, K. M.; Bruins, L. A.; Schmidt, J. E.; Chang, X. B.; Langlais, P.; Luo, M.; Jedlowski, P.; LaPlant, B.; Laumann, K.; Fonseca, R.; Bergsagel, P. L.; Mikhael, J.; Lacy, M.; Champion, M. D.; Stewart, A. K., Identification of cereblon-binding proteins and relationship with response and survival after IMiDs in multiple myeloma. Blood 2014, 124 (4), 536–45.

17. Niphakis, M. J.; Lum, K. M.; Cognetta, A. B., 3rd; Correia, B. E.; Ichu, T. A.; Olucha, J.; Brown, S. J.; Kundu, S.; Piscitelli, F.; Rosen, H.; Cravatt, B. F., A Global Map of Lipid-Binding Proteins and Their Ligandability in Cells. Cell 2015, 161 (7), 1668–80.

18. Parker, C. G.; Galmozzi, A.; Wang, Y.; Correia, B. E.; Sasaki, K.; Joslyn, C. M.; Kim, A. S.; Cavallaro, C. L.; Lawrence, R. M.; Johnson, S. R.; Narvaiza, I.; Saez, E.; Cravatt, B. F., Ligand and Target Discovery by Fragment-Based Screening in Human Cells. Cell 2017, 168 (3), 527–541.e29.

19. Li, Z.; Hao, P.; Li, L.; Tan, C. Y. J.; Cheng, X.; Chen, G. Y. J.; Sze, S. K.; Shen, H.-M.; Yao, S. Q., Design and Synthesis of Minimalist Terminal Alkyne-Containing Diazirine Photo-Crosslinkers and Their Incorporation into Kinase Inhibitors for Cell- and Tissue-Based Proteome Profiling. Angew Chem Int Ed 2013, 52 (33), 8551–8556.

20. Gao, J.; Mfuh, A.; Amako, Y.; Woo, C. M., Small Molecule Interactome Mapping by Photoaffinity Labeling Reveals Binding Site Hotspots for the NSAIDs. J Am Chem Soc 2018, 140 (12), 4259–4268.

21. Miyamoto, D. K.; Flaxman, H. A.; Wu, H.-Y.; Gao, J.; Woo, C. M., Discovery of a Celecoxib Binding Site on Prostaglandin E Synthase (PTGES) with a Cleavable Chelation-Assisted Biotin Probe. ACS Chem Biol 2019, 14 (12), 2527–2532.

22. West, A. V.; Muncipinto, G.; Wu, H.-Y.; Huang, A. C.; Labenski, M. T.; Jones, L. H.; Woo, C. M., Labeling Preferences of Diazirines with Protein Biomolecules. J Am Chem Soc 2021, 143 (17), 6691–6700.

23. Flaxman, H. A.; Chang, C. F.; Wu, H. Y.; Nakamoto, C. H.; Woo, C. M., A Binding Site Hotspot Map of the FKBP12-Rapamycin-FRB Ternary Complex by Photoaffinity Labeling and Mass Spectrometry-Based Proteomics. J Am Chem Soc 2019, 141 (30), 11759–11764.

24. Fischer, E. S.; Bohm, K.; Lydeard, J. R.; Yang, H.; Stadler, M. B.; Cavadini, S.; Nagel, J.; Serluca, F.; Acker, V.; Lingaraju, G. M.; Tichkule, R. B.; Schebesta, M.; Forrester, W. C.; Schirle, M.; Hassiepen, U.; Ottl, J.; Hild, M.; Beckwith, R. E.; Harper, J. W.; Jenkins, J. L.; Thoma, N. H., Structure of the DDB1-CRBN E3 ubiquitin ligase in complex with thalidomide. Nature 2014, 512 (7512), 49–53.

25. Chamberlain, P. P.; Lopez-Girona, A.; Miller, K.; Carmel, G.; Pagarigan, B.; Chie-Leon, B.; Rychak, E.; Corral, L. G.; Ren, Y. J.; Wang, M.; Riley, M.; Delker, S. L.; Ito, T.; Ando, H.; Mori, T.; Hirano, Y.; Handa, H.; Hakoshima, T.; Daniel, T. O.; Cathers, B. E., Structure of the human Cereblon-DDB1-lenalidomide complex reveals basis for responsiveness to thalidomide analogs. Nat Struct Mol Biol 2014, 21 (9), 803–9.

26. Ishoey, M.; Chorn, S.; Singh, N.; Jaeger, M. G.; Brand, M.; Paulk, J.; Bauer, S.; Erb, M. A.; Parapatics, K.; Muller, A. C.; Bennett, K. L.; Ecker, G. F.; Bradner, J. E.; Winter, G. E., Translation Termination Factor GSPT1 Is a Phenotypically Relevant Off-Target of Heterobifunctional Phthalimide Degraders. ACS Chem Biol 2018, 13 (3), 553–560.

27. Zhu, Y. X.; Yin, H.; Bruins, L. A.; Shi, C. X.; Jedlowski, P.; Aziz, M.; Sereduk, C.; Kortuem, K. M.; Schmidt, J. E.; Champion, M.; Braggio, E.; Keith Stewart, A., RNA interference screening identifies lenalidomide sensitizers in multiple myeloma, including RSK2. Blood 2015, 125 (3), 483–91.

28. Mitsiades, N.; Mitsiades, C. S.; Poulaki, V.; Chauhan, D.; Richardson, P. G.; Hideshima, T.; Munshi, N. C.; Treon, S. P.; Anderson, K. C., Apoptotic signaling induced by immunomodulatory thalidomide analogs in human multiple myeloma cells: therapeutic implications. Blood 2002, 99 (12), 4525–30.

29. Mahony, C.; Erskine, L.; Niven, J.; Greig, N. H.; Figg, W. D.; Vargesson, N., Pomalidomide is nonteratogenic in chicken and zebrafish embryos and nonneurotoxic in vitro. Proc Natl Acad Sci 2013, 110 (31), 12703–12708.

30. Lupas, A. N.; Zhu, H.; Korycinski, M., The thalidomide-binding domain of cereblon defines the CULT domain family and is a new member of the beta-tent fold. PLoS Comput Biol 2015, 11 (1), e1004023.

31. Sievers, Q. L.; Petzold, G.; Bunker, R. D.; Renneville, A.; Slabicki, M.; Liddicoat, B. J.; Abdulrahman, W.; Mikkelsen, T.; Ebert, B. L.; Thoma, N. H., Defining the human C2H2 zinc finger degrome targeted by thalidomide analogs through CRBN. Science 2018, 362 (6414), eaat0572.

32. Nguyen, T. V.; Lee, J. E.; Sweredoski, M. J.; Yang, S. J.; Jeon, S. J.; Harrison, J. S.; Yim, J. H.; Lee, S. G.; Handa, H.; Kuhlman, B.; Jeong, J. S.; Reitsma, J. M.; Park, C. S.; Hess, S.; Deshaies, R. J., Glutamine Triggers Acetylation-Dependent Degradation of Glutamine Synthetase via the Thalidomide Receptor Cereblon. Mol Cell 2016, 61 (6), 809–20.

33. Nowak, R. P.; DeAngelo, S. L.; Buckley, D.; He, Z.; Donovan, K. A.; An, J.; Safaee, N.; Jedrychowski, M. P.; Ponthier, C. M.; Ishoey, M.; Zhang, T.; Mancias, J. D.; Gray, N. S.; Bradner, J. E.; Fischer, E. S., Plasticity in binding confers selectivity in ligand-induced protein degradation. Nat Chem Biol 2018, 14 (7), 706–714.

34. Hideshima, T.; Ogiya, D.; Liu, J.; Harada, T.; Kurata, K.; Bae, J.; Massefski, W.; Anderson, K. C., Immunomodulatory drugs activate NK cells via both Zap-70 and cereblon-dependent pathways. Leukemia 2021, 35 (1), 177–188.

35. Tateno, S.; Iida, M.; Fujii, S.; Suwa, T.; Katayama, M.; Tokuyama, H.; Yamamoto, J.; Ito, T.; Sakamoto, S.; Handa, H.; Yamaguchi, Y., Genome-wide screening reveals a role for subcellular localization of CRBN in the anti-myeloma activity of pomalidomide. Sci Rep 2020, 10 (1), 4012.

36. Wang, Y.; Dix, M. M.; Bianco, G.; Remsberg, J. R.; Lee, H.-Y.; Kalocsay, M.; Gygi, S. P.; Forli, S.; Vite, G.; Lawrence, R. M.; Parker, C. G.; Cravatt, B. F., Expedited mapping of the ligandable proteome using fully functionalized enantiomeric probe pairs. Nat Chem 2019, 11 (12), 1113–1123.

37. Liu, Y.; Huang, X.; He, X.; Zhou, Y.; Jiang, X.; Chen-Kiang, S.; Jaffrey, S. R.; Xu, G., A novel effect of thalidomide and its analogs: suppression of cereblon ubiquitination enhances ubiquitin ligase function. FASEB J 2015, 29 (12), 4829–39.

38. Lopez-Girona, A.; Mendy, D.; Ito, T.; Miller, K.; Gandhi, A. K.; Kang, J.; Karasawa, S.; Carmel, G.; Jackson, P.; Abbasian, M.; Mahmoudi, A.; Cathers, B.; Rychak, E.; Gaidarova, S.; Chen, R.; Schafer, P. H.; Handa, H.; Daniel, T. O.; Evans, J. F.; Chopra, R., Cereblon is a direct protein target for immunomodulatory and antiproliferative activities of lenalidomide and pomalidomide. Leukemia 2012, 26 (11), 2326–35.

39. Heider, M.; Eichner, R.; Stroh, J.; Morath, V.; Kuisl, A.; Zecha, J.; Lawatscheck, J.; Baek, K.; Garz, A. K.; Rudelius, M.; Deuschle, F. C.; Keller, U.; Lemeer, S.; Verbeek, M.; Gotze, K. S.; Skerra, A.; Weber, W. A.; Buchner, J.; Schulman, B. A.; Kuster, B.; Fernandez-Saiz, V.; Bassermann, F., The IMiD target CRBN determines HSP90 activity toward transmembrane proteins essential in multiple myeloma. Mol Cell 2021, 81 (6), 1170–1186 e10.

40. Schreiber, S. L., The Rise of Molecular Glues. Cell 2021, 184 (1), 3–9.

41. Donovan, K. A.; Ferguson, F. M.; Bushman, J. W.; Eleuteri, N. A.; Bhunia, D.; Ryu, S.; Tan, L.; Shi, K.; Yue, H.; Liu, X.; Dobrovolsky, D.; Jiang, B.; Wang, J.; Hao, M.; You, I.; Teng, M.; Liang, Y.; Hatcher, J.; Li, Z.; Manz, T. D.; Groendyke, B.; Hu, W.; Nam, Y.; Sengupta, S.; Cho, H.; Shin, I.; Agius, M. P.; Ghobrial, I. M.; Ma, M. W.; Che, J.; Buhrlage, S. J.; Sim, T.; Gray, N. S.; Fischer, E. S., Mapping the Degradable Kinome Provides a Resource for Expedited Degrader Development. Cell 2020, 183 (6), 1714–1731 e10.

42. Smith, B. E.; Wang, S. L.; Jaime-Figueroa, S.; Harbin, A.; Wang, J.; Hamman, B. D.; Crews, C. M., Differential PROTAC substrate specificity dictated by orientation of recruited E3 ligase. Nat Commun 2019, 10 (1), 131.

43. Zeng, M.; Xiong, Y.; Safaee, N.; Nowak, R. P.; Donovan, K. A.; Yuan, C. J.; Nabet, B.; Gero, T. W.; Feru, F.; Li, L.; Gondi, S.; Ombelets, L. J.; Quan, C.; Janne, P. A.; Kostic, M.; Scott, D. A.; Westover, K. D.; Fischer, E. S.; Gray, N. S., Exploring Targeted Degradation Strategy for Oncogenic KRAS(G12C). Cell Chem Biol 2020, 27 (1), 19–31 e6.

44. Simonetti, A.; Brito Querido, J.; Myasnikov, A. G.; Mancera-Martinez, E.; Renaud, A.; Kuhn, L.; Hashem, Y., eIF3 Peripheral Subunits Rearrangement after mRNA Binding and Start-Codon Recognition. Mol Cell 2016, 63 (2), 206–217.

