## Supplementary material for "Development of photo-lenalidomide for cellular target identification": SI

<u>*Supporting Information*</u>

**Index**

Additional Experimental Information ........................................................................... S3

**Scheme S1 |** Synthetic scheme for photo-lenalidomide and analogs **1–6**.

**Figure S1 |** Western blot of IKZF1 and IKZF3 in MM.1S cells.
**Figure S2 |** Western blot of CK1α and GSPT1 in MM.1S cells.
**Figure S3 |** Cell viability (MTT) assay of Len and pLen in Jurkat, K562, RAW264.7 and MOLM13 cells after 5 d treatment.
**Figure S4 |** Immunomodulatory response of interleukin-2 (IL-2), CD69 and TNF-α.
**Figure S5 |** Binding site mapping of photo-lenalidomide analogs with the thalidomide-binding domain of CRBN (CULT domain).
**Figure S6 |** Docked structure of CRBN–analog **3**–IKZF1 ZF2 complex adapted from PDB 6H0F.
**Figure S7 |** In-gel fluorescence imaging of protein targets conjugated with photo-lenalidomide in HEK293T-CRBN cells.
**Figure S8 |** Co-immunoprecipitation of eIF3i in MM.1S cells by endogenous CRBN with or without lenalidomide.
**Figure S9 |** Supporting data related to **Figure 4**.
**Figure S10 |** Uncropped immunoblots related to **Figure 1c**.
**Figure S11 |** Uncropped immunoblots related to **Figure 3a** and **Figure 3d**.
**Figure S12 |** Uncropped immunoblots related to **Figure 4b** and **Figure 4d**.
**Figure S13 |** Uncropped immunoblots related to **Figure 4e**, **Figure 4f** and **Figure 4g**.

**Table S1 |** Complete list of pLen-modified peptide spectral matches identified by LC-MS/MS.
**Table S2 |** Unique pLen-modified peptide assignments identified by LC-MS/MS.
**Table S3 |** Complete list of analog **1**-modified peptide spectral matches identified by LC-MS/MS.

**Table S4 |** Unique analog **1**-modified peptide assignments identified by LC-MS/MS.
**Table S5 |** Complete protein list identified by LC-MS/MS in **Figure 3c**.
**Table S6 |** Complete protein list identified by LC-MS/MS in **Figure 4a**.
**Table S7 |** Complete protein list identified by LC-MS/MS in **Figure 4c**.

General methods and instrumentation ........ S17
General chemical methods ........ S17
Chemical instrumentation ........ S17
Synthetic procedures ........ S19
Antibodies and biological reagents ........ S35
Cell culture and transfection ........ S35
Plasmids and subcloning ........ S35
Overexpression and purification of CULT ........ S36
Structural proteomics mass spectrometry sample preparation ........ S37
Structural proteomics mass spectrometry procedures ........ S37
Structural proteomics data analysis procedures ........ S38
Structural modeling details ........ S38
Immunoprecipitation assays in live cells ........ S38
Cycloheximide (CHX) treatment ........ S39
Photo-affinity labeling experiment and in-gel fluorescent imaging ........ S39
Cell viability assay (MTT) ........ S39
Flow cytometry (FACS) ........ S39
RNA isolation and qRT-PCR experiment ........ S40
Luciferase reporter assay ........ S40
In vivo thermal shift assay ........ S40
Photo-affinity labeling and enrichment of lenalidomide targets in living cells and on-bead digestion ........ S41
Quantitative mass spectrometry sample preparation ........ S41
Quantitative mass spectrometry analysis ........ S41
In vivo ubiquitylation assay ........ S42
Software ........ S42
Statistical analysis ........ S42
References ........ S43
NMR and IR spectrum of compounds ........ S44

### SUPPORTING INFORMATION FIGURES

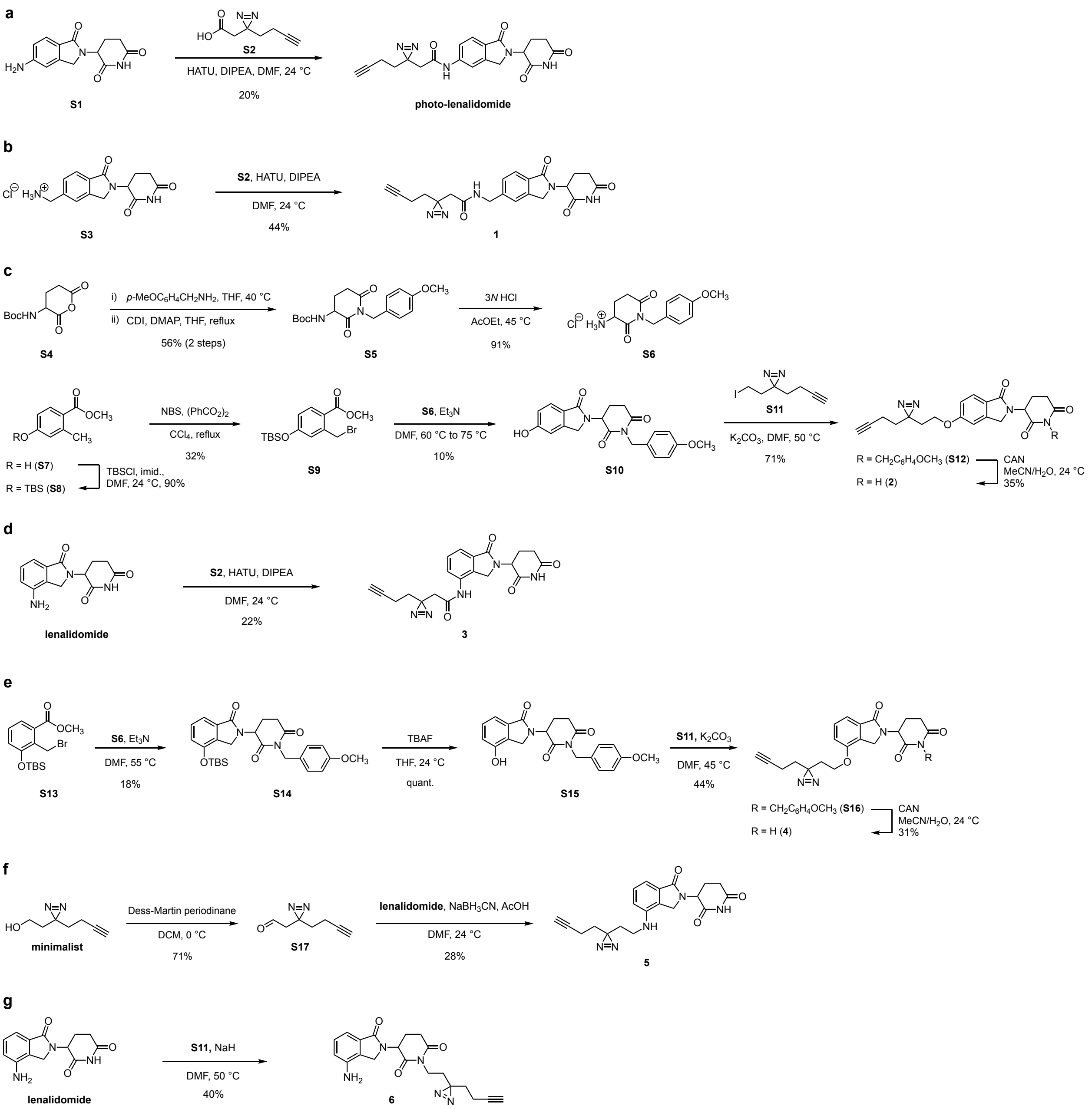


**Scheme S1 |** Synthetic scheme for photo-lenalidomide and analogs **1–6**. Synthesis of (**a**) photo-lenalidomide, (**b**) analog **1**, (**c**) analog **2**, (**d**) analog **3**, (**e**) analog **4**, (**f**) analog **5**, and (**g**) analog **6**.

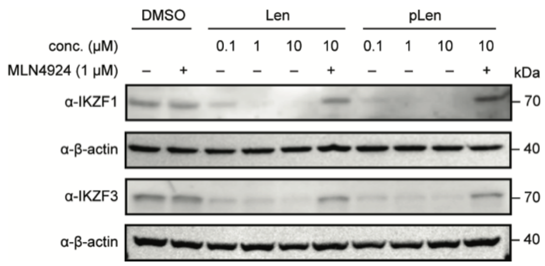


**Figure S1 |** Western blot of IKZF1 and IKZF3 after treatment of MM.1S cells with the indicated concentration of Len or pLen of 8 h with or without 1 μM MLN4924. Results are representative of three biologically independent experiments.

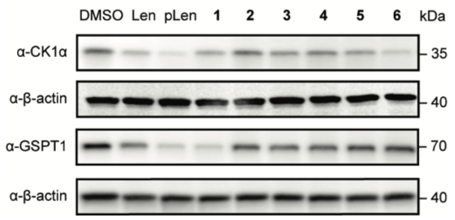


**Figure S2 |** Western blot of CK1α and GSPT1 after treatment of MM.1S cells with 10 μM Len, pLen or analogs **1–6** for 8 h. Results are representative of three biologically independent experiments.

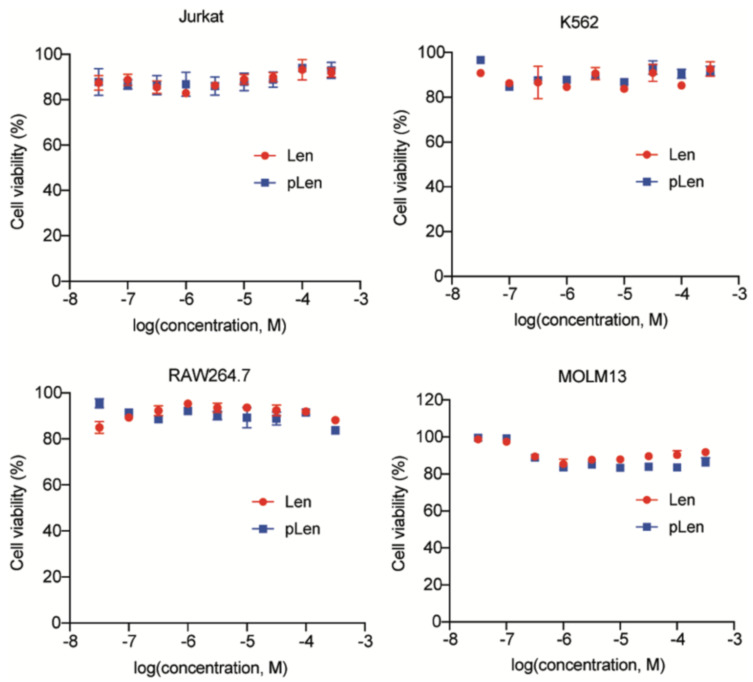


**Figure S3 |** Cell viability (MTT) assay of Len and pLen in Jurkat, K562, RAW264.7 and MOLM13 cells after 5 d treatment. Results are representative of two biologically independent experiments. Data was normalized to the untreated samples. All graphs are shown as mean ± s.d., n = 3.

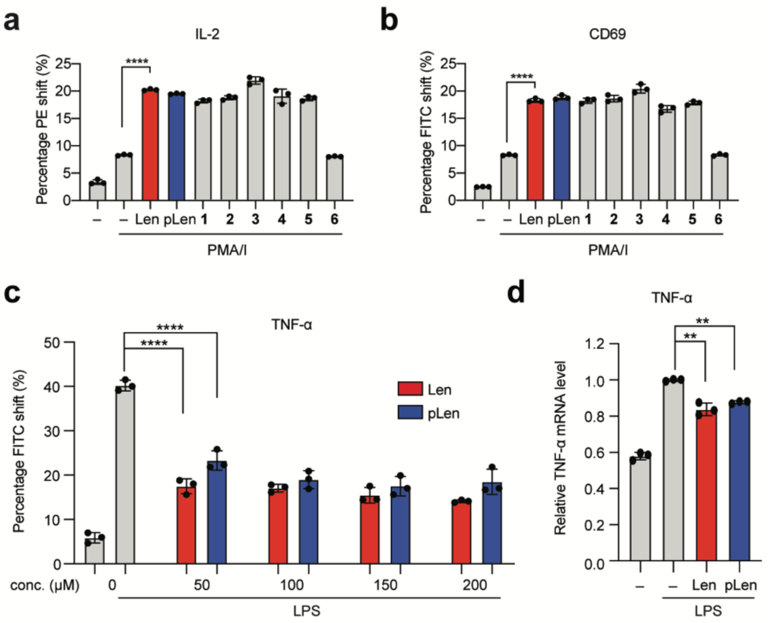


**Figure S4 |** Immunomodulatory response of interleukin-2 (IL-2) (**a**) and CD69 (**b**) in Jurkat cells treated with 1 μM Len, pLen, or analogs **1**–**6**. Jurkat cells were treated with the indicated compound (1 μM) for 1 h before stimulation with 1 ng/mL phorbol 12-myristate 13-acetate and 1 mg/mL ionomycin (PMA/I) for 48 h. Data represents the % IL-2- or CD69-positive cell numbers among total cell counts of 100,000. Results are representative of four biologically independent experiments. (**c**, **d**) TNF-α protein and mRNA level in RAW264.7 cells upon treatment. (**c**) Levels of TNF-α in RAW264.7 cells measured by flow cytometry. The cells were treated with Len or pLen at the indicated concentration for 1 h and stimulated with 1 ng/mL lipopolysaccharide (LPS) for 3 h prior to flow cytometry analysis. (**d**) Levels of TNF-α mRNA in RAW264.7 cells measured by qRT-PCR. The cells were treated with 10 μM Len or pLen for 48 h and stimulated with 1 ng/mL LPS for 3 h prior to qRT-PCR analysis. Results are representative of four biologically independent experiments. Statistical significance was analyzed using a 2-way Student's t-test. *P < 0.05; **P < 0.005; ***P < 0.0005, ****P < 0.0001; n.s. not significant.

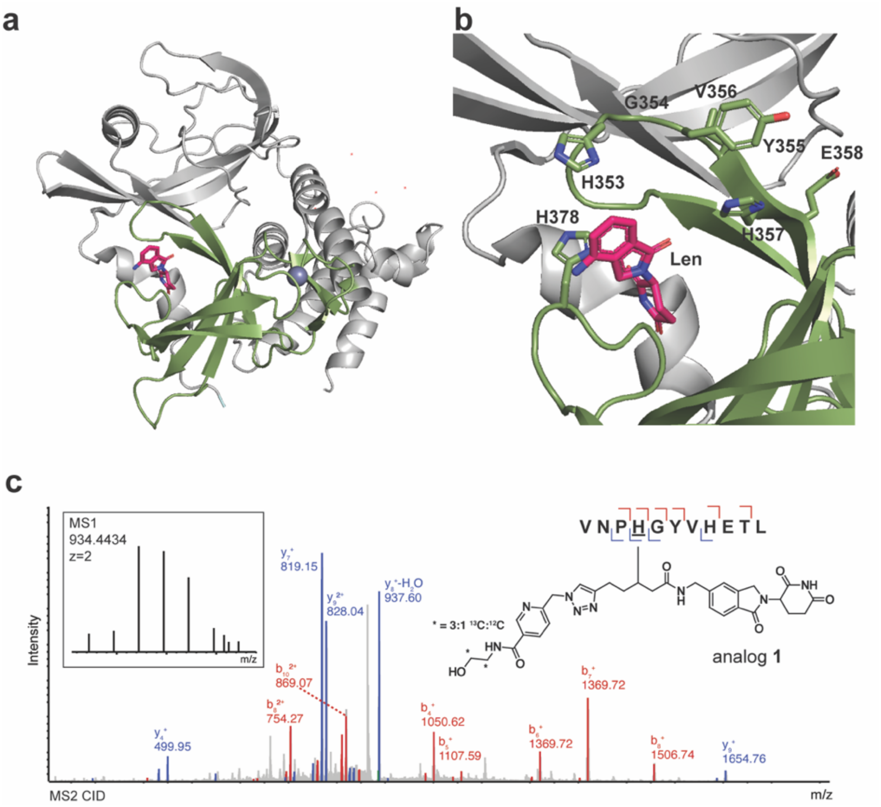


**Figure S5 |** Binding site mapping of photo-lenalidomide analogs with the thalidomide-binding domain of CRBN (CULT domain). (**a**) Structure of the DDB1-CRBN-lenalidomide complex (PDB 4CI2, DDB1 is not shown) with the CULT domain highlighted in green and the rest of CRBN in grey. (**b**) Enlarged view of the tri-tryptophan binding pocket in the thalidomide-binding domain of CRBN. Side chains of amino acids modified by photo-lenalidomide (His378, etc.) or analog **1** (His378, etc.) are shown in stick form. (**c**) Representative assignment of a peptide in CULT conjugated to analog **1**.

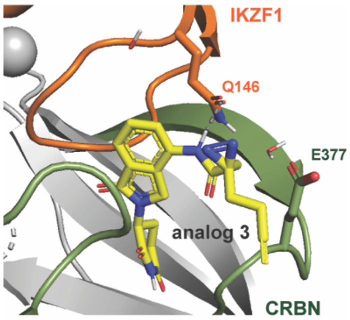


**Figure S6 |** Docked structure of CRBN–analog **3**–IKZF1 ZF2 complex adapted from PDB 6H0F. The conjugated peptides of CRBN are colored green and the rest of the protein is colored gray. IKZF1 ZF2 is colored orange.

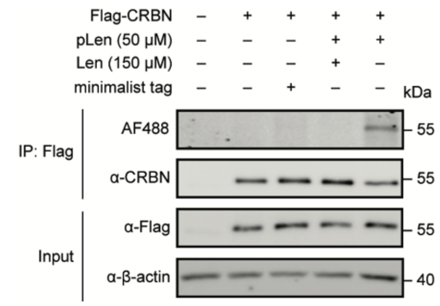


**Figure S7 |** In-gel fluorescence imaging of protein targets conjugated with photo-lenalidomide in HEK293T-CRBN cells. HEK293T-CRBN cells were treated with the indicated compound(s) for 1 h prior to photo-irradiation and lysis. The target proteins were immunoprecipitated, subjected to click chemistry with AF488-azide, and visualized by in-gel fluorescence imaging or Western blot analysis. Results are representative of three biologically independent experiments.

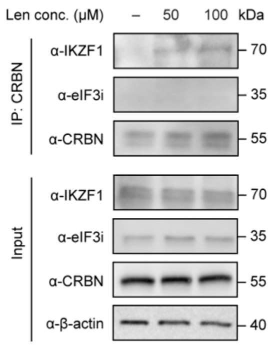


**Figure S8 |** Co-immunoprecipitation of eIF3i in MM.1S cells by endogenous CRBN with or without lenalidomide. MM.1S cells were treated with lenalidomide at the indicated concentration for 1 h prior to lysis. The target proteins were immunoprecipitated and visualized by immunoblotting. Results are representative of three biologically independent experiments.

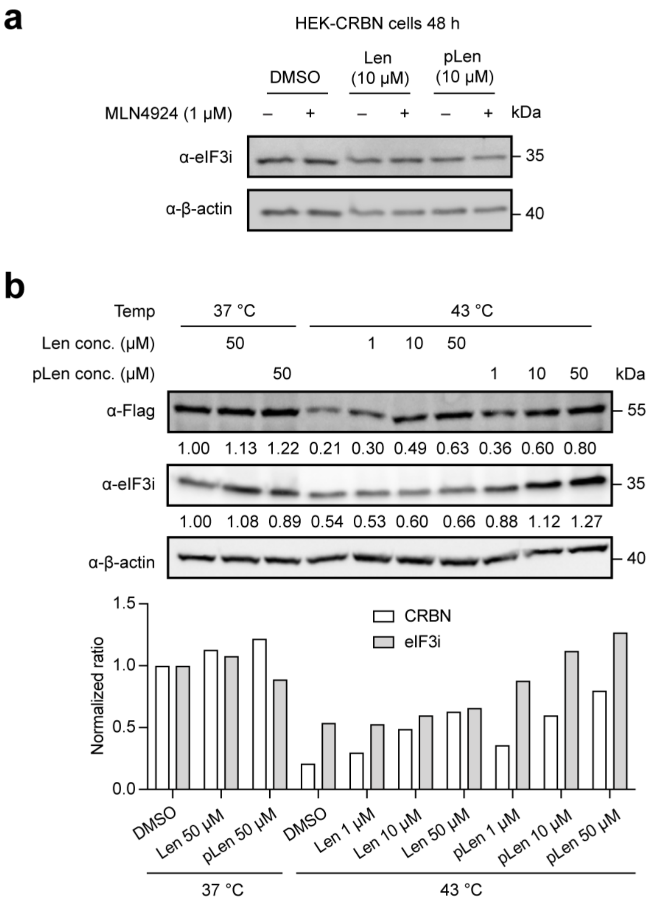


**Figure S9 |** Supporting data related to **Figure 4**. (**a**) Western blot of eIF3i after treatment with Len or pLen in HEK293T-CRBN cells for 48 h with and without 1 μM MLN4924. (**b**) Thermal shift assay for Flag-CRBN and eIF3i in HEK293T-CRBN cells. The cells were incubated with or without Len or pLen for 1 h and then incubated at the indicated temperature for 5 min followed by cell lysis. Cell lysates were cleared by centrifugation before immunoblotting. Results are representative of two biologically independent experiments.

Figure 1c

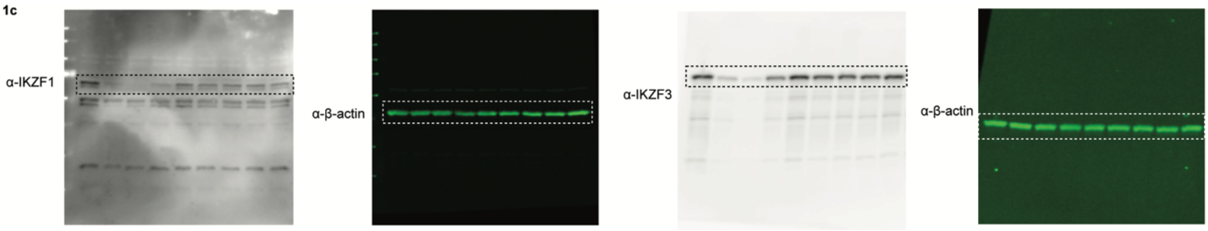


**Figure S10 |** Uncropped immunoblots related to **Figure 1c**.

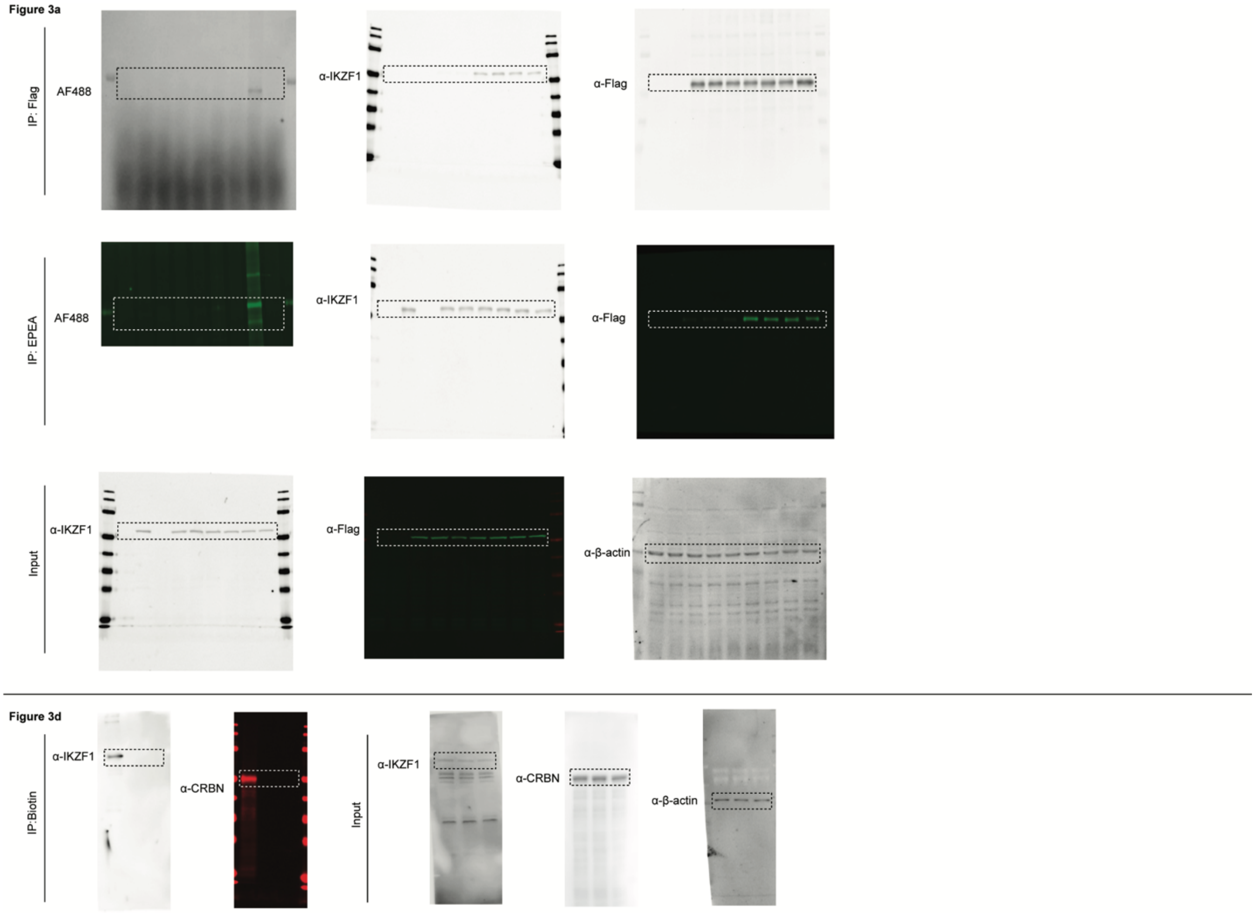


**Figure S11 |** Uncropped immunoblots related to **Figure 3a** and **Figure 3d**.

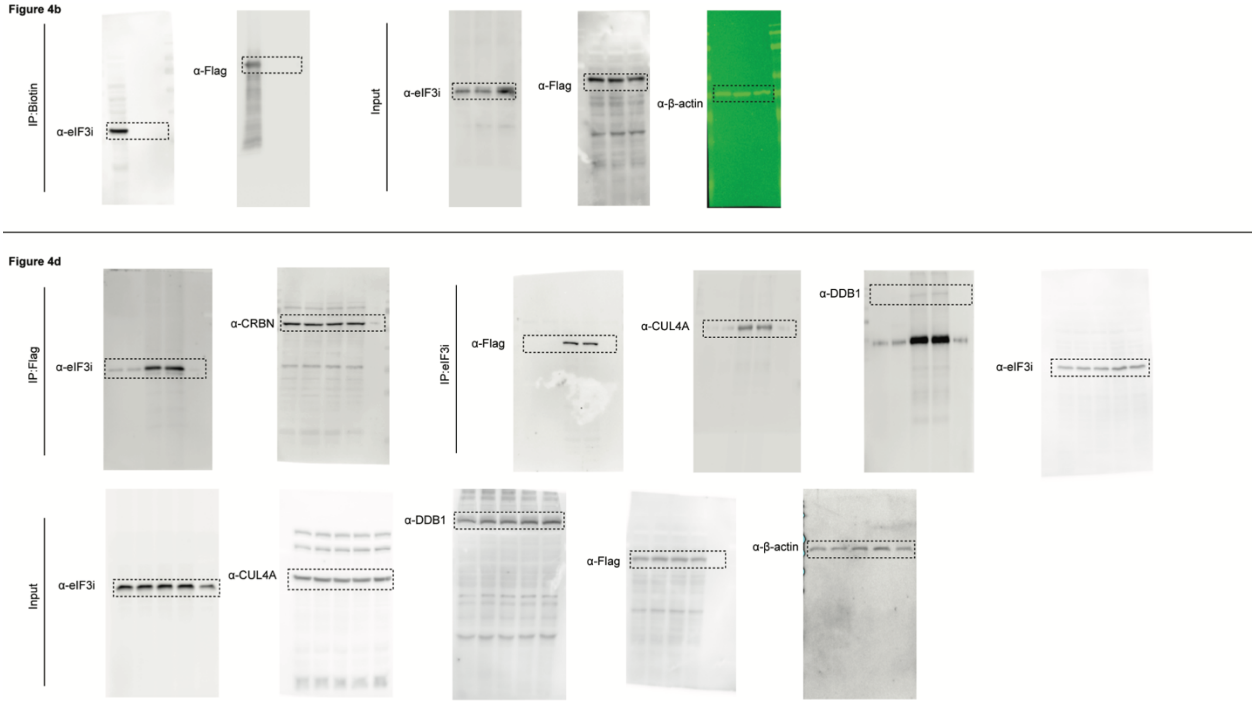


**Figure S12 |** Uncropped immunoblots related to **Figure 4b** and **Figure 4d**.

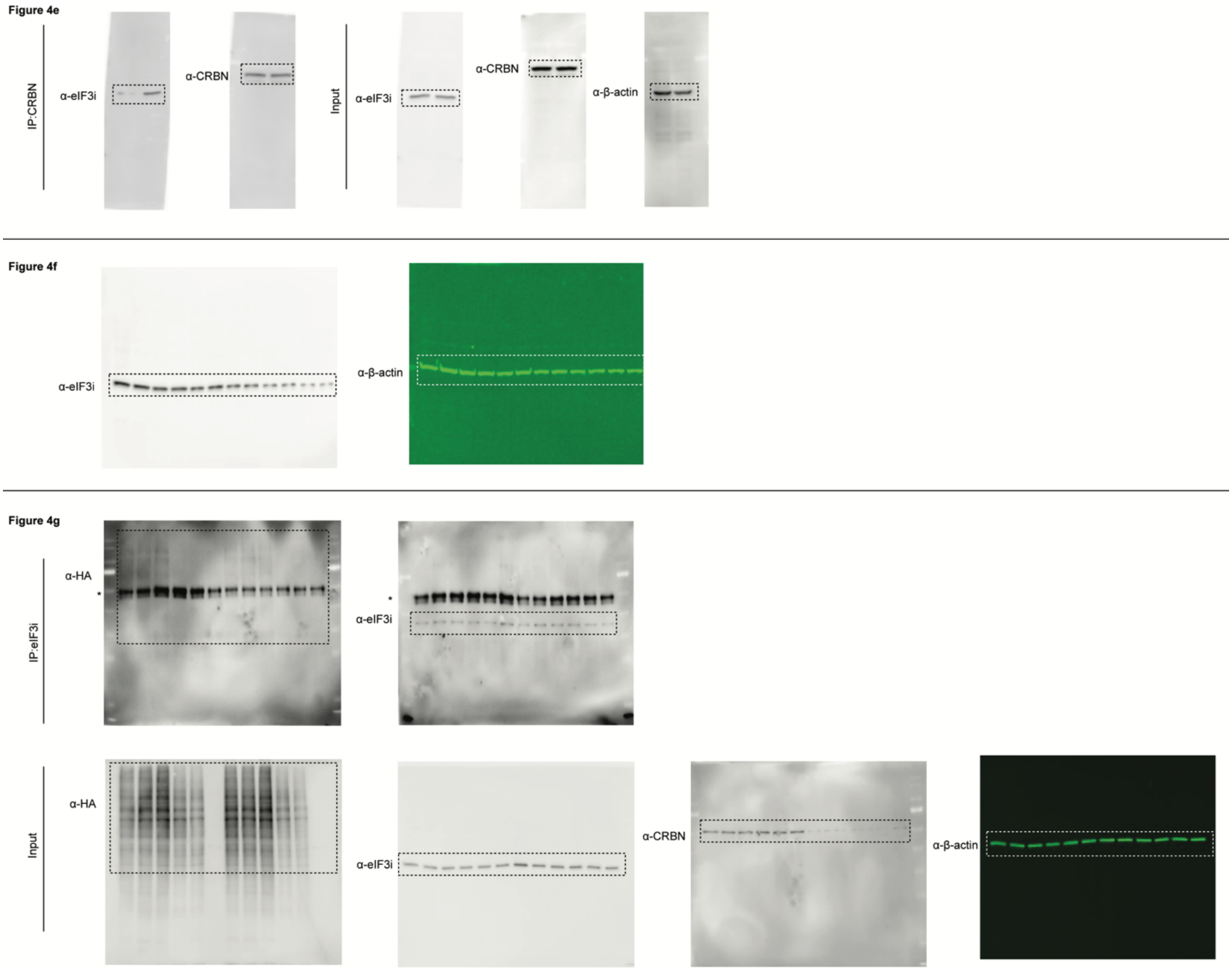


**Figure S13 |** Uncropped immunoblots related to **Figure 4e**, **Figure 4f** and **Figure 4g**.

### Methods

#### General methods and instrumentation.

Photo-irradiation was performed using a Dymax ECE 5000 UV Light-Curing Flood Lamp system (Cat # 41060) in a 4 ºC cold room unless otherwise noted. The lamp was turned on for at least 15 minutes prior to use.

Protein quantification was performed by bicinchoninic acid assay on a multimode microplate reader FilterMax F3 (Molecular Devices LLC, Sunnyvale, CA). Sonication of cells or protein pellets was performed using a Branson SFX250 Sonifier (Branson Ultrasonics, Brookfield, CT). In-gel fluorescence, IR blots and chemiluminescence measurements were detected on an Azure Imager C600 (Azure Biosystems, Inc., Dublin, CA). DNA or RNA concentration was measured with a NanoDrop One UV-vis spectrophotometer (Thermo Fisher, Cat # ND-ONE-W).

Proteomics experiments were performed on a Lumos Tribrid Orbitrap (Thermo Scientific) equipped with an UltiMate 3000 RSLCnano System (Thermo Scientific) within the Mass Spectrometry and Proteomics Resource Laboratory at Harvard University (MSPRL). Peptides were dried using an Eppendorf Vacufuge (Speed-Vac). Cell sorting experiments were performed on a BD FACSAria II cell sorter (BD Bioscience). Flow cytometry quantification were performed on BD LSR Fortessa (BD Bioscience, SORP) and BD LSR II flow cytometric analyzers (BD Biosciences, SORP).

For immunoblotting analysis, proteins were transferred from SDS-PAGE gels to nitrocellulose membranes using iBlot-2 dry blotting system (Thermo Scientific). Membranes were blocked with Tris buffered saline containing 0.1% Tween-20 and 5% BSA and incubated with the primary antibodies and the secondary antibodies sequentially. Immunoblots images were captured by Azure Imager C600 (Azure Biosystems, Dublin, CA).

#### General chemical methods.

All reactions were performed in single-neck, oven-dried, round-bottomed flasks fitted with rubber septa under a positive pressure of argon, unless otherwise noted. Air- and moisture-sensitive liquids were transferred via syringe or stainless-steel cannula. Organic solutions were concentrated by rotary evaporation at 30–33 °C. Normal phase flash-column chromatography was performed as described by Still and co-workers.[1] Normal phase purifications employ silica gel (60 Å, 40–63 µm particle size) purchased from Silicycle (Quebec, Canada). Analytical thin-layer chromatography (TLC) was performed using glass plates pre-coated with silica gel (0.25 mm, 60 Å pore size) containing a fluorescent indicator (254 nm). TLC plates were visualized by exposure to ultraviolet light (UV), iodine ($I_2$), and/or submersion in *p*-anisaldehyde or ninhydrin followed by brief heating with a heat gun (10–15 s).

Commercial solvents and reagents were used as received with the following exceptions. Lenalidomide was obtained from BioVision (Cat # 1862-25). Minimalist tag was prepared according to the method of Yao and co-workers.[2] The cleavable biotin azide probe was synthesized according to the method of Woo and co-workers[3] and the cleavable biotin picolyl-azide probe was synthesized according to the method of Woo and co-workers.[4] Cycloheximide (CHX) was purchased from Sigma-Aldrich (Cat # C4859). Dichloromethane, *N*,*N*-dimethylformamide and tetrahydrofuran were purified according to the method of Grubbs and co-workers.[5] Triethylamine was distilled over calcium hydride under an atmosphere of argon.

#### Chemical instrumentation.

Proton nuclear magnetic resonance spectra ($^1$H NMR) were recorded at 500 MHz at 25 °C unless otherwise noted. Chemical shifts are expressed in parts per million (ppm, δ scale) downfield from tetramethylsilane and are referenced to residual protium in the NMR solvent [$CHCl_3$, δ 7.26; $CHD_2OD$, δ 3.31; $(CHD_2)(CD_3)SO$, δ 2.49]. Data are represented as follows: chemical shift, multiplicity (s = singlet, d = doublet, t = triplet, q = quartet, quin = quintet, m = multiplet and/or multiple resonances, br = broad, app = apparent), integration, coupling constant in Hertz, and assignment. Proton-decoupled carbon nuclear magnetic resonance spectra ($^{13}C$ NMR) were recorded at 125 MHz at 25 °C unless otherwise noted. Chemical shifts are expressed in parts per million (ppm, δ scale) downfield from tetramethylsilane and are referenced to the carbon resonances of the solvent [$CDCl_3$, δ 77.2; $CD_3OD$, δ 49.0; $(CD_3)_2SO$, δ 39.5]. $^{13}C$ NMR and data are represented as follows: chemical shift, carbon type. Infrared (IR) spectra were obtained using a Shimadzu 8400S FT-IR spectrometer referenced to a polystyrene standard. Data are represented as follows: frequency of absorption ($cm^{-1}$), intensity of absorption (s = strong, m = medium, w = weak, br = broad). Small molecule high-resolution mass spectrometry (HRMS) measurements were obtained at the Harvard University Chemistry and Chemical Biology Department Mass Spectrometry Facility using an Agilent 1260 UPLC-MS Bruker connected in line with a microTOF-Q II hybrid quadrupole-time of flight MS or at the Harvard Center for Mass Spectrometry using a Thermo q-Exactive Plus. Preparative HPLC was performed on Waters Prep 150 LC System with XBridge BEH $C_{18}$ OBD Prep Column, 130 Å, 5 μm, 19 mm x 100 mm.

### Synthetic procedures.

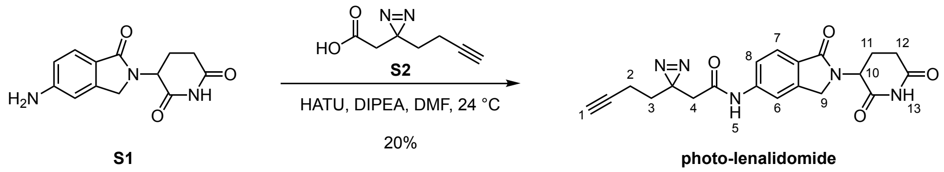


### Synthesis of photo-lenalidomide:

The glutarimide **S1**[6] (19.2 mg, 74.1 µmol, 1.20 equiv.) was added in a single portion to a stirred solution of the carboxylic acid **S2**[7] (9.4 mg, 61.8 µmol, 1.00 equiv.). Sodium sulfate (26.3 mg, 185 µmol, 3.00 equiv.), HATU (70.5 mg, 197 µmol, 3.60 equiv.) and *N,N*-diisopropylethylamine (32.4 µL, 185 µmol, 3.60 equiv.) were sequentially added to the mixture and the reaction mixture was stirred for 24 h at 25 ˚C. The product mixture was diluted with ethyl acetate (5.0 mL) and transferred to a separatory funnel. The diluted product mixture was washed with saturated aqueous sodium chloride (5.0 mL) and 10% aqueous citric acid (5.0 mL). The layers were mixed, and the aqueous layer was extracted by ethyl acetate (3 x 5.0 mL). The combined organic layers were dried over sodium sulfate and concentrated to dryness. The crude product was purified by flash-column chromatography (eluting with 2% methanol–dichloromethane, grading to 4% methanol–dichloromethane, three steps). The obtained product mixture was further purified by HPLC (grading 95% to 5% water–acetonitrile over 25 min, flow rate = 10 mL/min, retention time = 12.3 min) to afford the titled compound as a colorless solid (**photo-lenalidomide**, 4.8 mg, 20%).

$R_f$ = 0.25 (4% methanol–dichloromethane; UV). $^1$H NMR (500 MHz, methanol-$d_4$): δ 8.03 (s, 1H, $H_6$), 7.74 (d, *J* = 8.3 Hz, 1H, $H_7$), 7.52 (dd, *J* = 8.3, 1.9 Hz, 1H, $H_8$), 5.14 (dd, *J* = 13.4, 5.2 Hz, 1H, $H_{10}$), 4.51 (d, *J* = 17.0 Hz, 1H, $H_9$), 4.45 (d, *J* = 17.0 Hz, 1H, $H_9$), 2.91 (ddd, *J* = 17.5, 13.5, 5.3 Hz, 1H, $H_{12}$), 2.78 (ddd, *J* = 17.5, 4.5, 2.4 Hz, 1H, $H_{12}$), 2.50 (dddd, *J* = 13.5, 13.5, 13.5, 4.5 Hz, 2H, $H_{11}$), 2.52 (s, 2H, $H_4$), 2.29 (t, *J* = 2.6 Hz, 1H, $H_1$), 2.17 (dddd, *J* = 13.5, 5.3, 5.3, 2.4 Hz, 1H, $H_{11}$), 2.11 (td, *J* = 7.4, 2.6 Hz, 2H, $H_2$), 1.79 (t, *J* = 7.4 Hz, 2H, $H_3$). $^{13}$C NMR (126 MHz, methanol-$d_4$): δ 173.3 (C), 170.8 (C), 169.7 (C), 167.9 (C), 143.6 (C), 142.2 (C), 126.7 (C), 123.6 (CH), 119.3 (CH), 113.7 (CH), 82.1 (C), 69.0 (CH), 52.2 (CH), 47.9 ($CH_2$), 41.0 ($CH_2$), 32.2 ($CH_2$), 31.0 ($CH_2$), 25.9 (C), 22.7 ($CH_2$), 12.4 ($CH_2$). IR (ATR-FTIR), cm$^{-1}$: 3277 (br), 3087 (br), 1671 (s), 1553 (m), 1491 (m), 1434 (m), 1370 (m), 1259 (m), 1234 (m), 1198 (m), 773 (w), 668 (w). HRMS-ESI (*m/z*): $[M+H]^+$ calculated for $C_{20}H_{20}N_5O_4$, 394.1510; found, 394.1512.

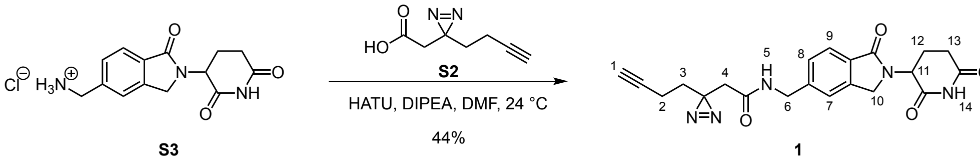

### Synthesis of analog 1:

The compound **S3**[8] (6.4 mg, 20.8 μmol, 1.00 equiv.) was added in a single portion to a stirred solution of **S2** (3.8 mg, 25.0 μmol, 1.20 equiv.) in *N,N*-dimethylformamide (210 μL). HATU (11.9 mg, 31.2 μmol, 1.50 equiv.) and *N,N*-diisopropylethylamine (10.9 μL, 62.4 μmol, 3.00 equiv.) were added in sequence to the reaction mixture at 25 ˚C. The reaction mixture was stirred for 24 h at 25 ˚C. The product mixture was diluted with ethyl acetate (5.0 mL) and transferred to a separatory funnel. The diluted product mixture was washed with saturated aqueous sodium chloride (5.0 mL) and 1*N* hydrochloric acid (5.0 mL). The aqueous layers were combined, and the aqueous layer was extracted by chloroform (3 x 5.0 mL). The organic layers were combined, and the combined organic layers were dried over sodium sulfate. The dried solution was filtered, and the filtrate was concentrated to dryness. The crude product was purified by flash-column chromatography (eluting with 2% methanol–dichloromethane, grading to 4% methanol–dichloromethane, three steps). The obtained product mixture was further purified by HPLC (grading 95% to 5% water–acetonitrile over 25 min, flow rate = 10 mL/min, retention time = 10.8 min), to afford the titled compound as a colorless solid (**1**, 4.4 mg, 44%).

$R_f$ = 0.25 (4% methanol–dichloromethane; UV, anis aldehyde). $^1$H NMR (400 MHz, dimethylsulfoxide-$d_6$): δ 10.97 (s, 1H, $H_{14}$), 8.54 (t, *J* = 6.0 Hz, 1H, $H_5$), 7.68 (d, *J* = 7.8 Hz, 1H, $H_9$), 7.46 (s, 1H, $H_7$), 7.39 (dd, *J* = 7.8, 1.4 Hz, 1H, $H_8$), 5.10 (dd, *J* = 13.2, 5.2 Hz, 1H, $H_5$), 4.44 (d, *J* = 17.3 Hz, 1H, $H_{10}$), 4.40 – 4.34 (m, 2H, $H_6$), 4.31 (d, *J* = 17.3 Hz, 1H, $H_{10}$), 2.91 (ddd, *J* = 17.3, 13.2, 5.2 Hz, 1H, $H_{13}$), 2.82 (t, *J* = 2.7 Hz, 1H, $H_1$), 2.60 (ddd, *J* = 17.3, 4.4, 2.0 Hz, 1H, $H_{13}$), 2.39 (dddd, *J* = 13.2, 13.2, 13.2, 4.4 Hz, 1H, $H_{12}$), 2.31 (s, 2H, $H_4$), 2.04 (dt, *J* = 7.5, 2.7 Hz, 2H, $H_2$), 2.00 (dddd, *J* = 13.2, 5.2, 5.2, 2.0 Hz, 1H, $H_{12}$), 1.67 (t, *J* = 7.5 Hz, 2H, $H_3$). $^{13}$C NMR (MHz,): $^{13}$C NMR (126 MHz, dimethylsulfoxide-$d_6$) δ 172.9 (C), 171.1 (C), 167.9 (C), 167.4 (C), 143.5 (C), 142.4 (C), 130.5 (C), 127.1 (CH), 123.0 (CH), 122.1 (CH), 83.1 (C), 71.8 (CH), 51.6 (CH), 47.1 ($CH_2$), 42.2 ($CH_2$), 40.0 ($CH_2$), 31.9 ($CH_2$), 31.2 ($CH_2$), 26.7 (C), 22.5 ($CH_2$), 12.6 ($CH_2$). δ. IR (ATR-FTIR), $cm^{-1}$: 3258 (br), 3095 (br), 1707 (s), 1400 (m), 1263 (m), 1178 (m), 746 (w), 725 (w). HRMS-ESI (*m/z*): $[M+H]^+$ calculated for $C_{21}H_{21}N_5O_4$, 408.1666; found, 408.1662.

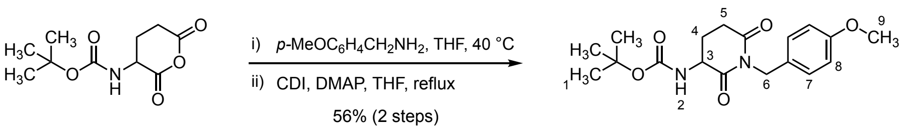


**S4** **S5**

### Synthesis of S5:

*Step 1*

*N*-Boc-glutamic acid anhydride (**S4**, 502 mg, 2.19 mmol, 1.00 equiv.) and *p*-methoxybenzyl amine (300 µL, 2.30 mmol, 1.05 equiv.) were dissolved in tetrahydrofuran (3.64 mL) at 25 °C. The resulting mixture was stirred for 24 h at 40 °C. The product mixture was cooled to 25 °C and acidified to pH 1 with 1*N* hydrochloric acid. The acidic aqueous layer was extracted by ethyl acetate (3 x 10 mL). The organic layers were combined, and the combined organic layers were dried over sodium sulfate and filtered. The filtrate was concentrated under vacuum and used directly in the following step.

*Step 2*

The reaction mixture obtained from the previous step was dissolved in tetrahydrofuran (21.9 mL) and carbonyl diimidazole (1.06 g, 6.57 mmol, 3.00 equiv.) and 4-dimethylaminopyridine (53.5 mg, 438 µmol, 0.20 equiv.) were added in sequence at 25 ˚C. The resulting mixture was heated at reflux for 24 h. The product mixture was cooled down to 25 °C and acidified to pH 1 with 1*N* hydrochloric acid. The acidic aqueous layer was extracted by ethyl acetate (3 x 20 mL). The organic layers were combined, and the combined organic layer were dried over sodium sulfate. The dried solution was filtered, and the filtrate was concentrated under vacuum. The crude mixture was purified by flash-column chromatography (eluting with 20% ethyl acetate–hexanes, grading to 40% ethyl acetate–hexanes, two steps) to afford the titled compound (**S5**, 428 mg, 56% over 2 steps).

$R_f$ = 0.25 (40% ethyl acetate–hexane; ninhydrin). $^1$H NMR (500 MHz, chloroform-*d*): δ 7.31 (d, *J* = 8.5 Hz, 2H, $H_8$), 6.81 (d, *J* = 8.5 Hz, 2H, $H_7$), 5.41 (bs, 1H, $H_2$), 4.90 (d, *J* = 13.5 Hz, 1H, $H_6$), 4.86 (d, *J* = 13.5 Hz, 1H, $H_6$), 4.30-4.22 (m, 1H, $H_3$), 3.77 (s, 3H, $H_9$), 2.86 (ddd, *J* = 18.0, 4.5, 2.0 Hz, 1H, $H_5$), 2.70 (ddd, *J* = 18.0, 13.5, 5.5 Hz, 1H, $H_5$), 2.51-2.43 (m, 1H, $H_4$), 1.77 (dddd, *J* = 13.5, 13.5, 13.5, 4.5 Hz, 1H, $H_4$), 1.45 (s, 9H, $H_1$). $^{13}$C NMR (125 MHz, chloroform-*d*): δ 171.9 (C), 171.2 (C), 159.0 (C), 155.6 (C), 130.4 (CH x 2), 129.1 (C), 113.8 (CH x 2), 80.2 (CH), 55.2 ($CH_2$), 52.5 (C), 43.1 ($CH_2$), 31.8 ($CH_2$), 28.3 ($CH_3$ x 3), 24.7 ($CH_2$). IR (ATR-FTIR), cm$^{-1}$ 3369 (br), 2975 (w), 1674 (m), 1513 (m), 1246 (m), 1156 (s), 729 (m). HRMS-ESI (*m/z*): $[M+H]^+$ calculated for $C_{18}H_{25}N_2O_5$, 349.1758; found, 349.1759.

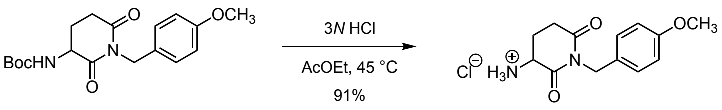


**Synthesis of S6:**

3*N*-Hydrochloric acid (2.4 mL, 7.20 mmol, 6.00 equiv.) was added to a stirred solution of the compound **S5** (418.5 mg, 1.20 mmol, 1.00 equiv.) in ethyl acetate (6.0 mL). The reaction mixture was stirred at 45 °C for 24 h. The product mixture was concentrated to dryness. The obtained residue was dissolved in benzene (5.0 mL) and concentrated to dryness to afford 1-(4-methoxybenzyl)-2,6-dioxopiperidin-3-aminium chloride as a white solid (**S6**, 310 mg, 91%). The product was used directly in the following step.

**Synthesis of S8:**

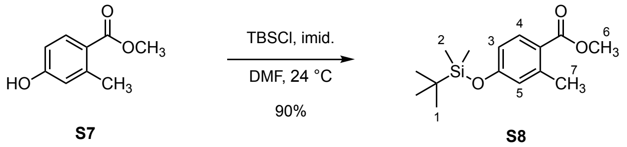


Imidazole (3.04 g, 44.8 mmol, 2.50 equiv.) was added in a single portion to a stirred solution of the compound **S7**[9] (2.98 g, 17.9 mmol, 1.00 equiv.) in *N,N*-dimethylformamide (9.9 mL) at 25 ˚C. The reaction mixture was cooled to 0 ˚C and *tert*-butyldimethylsilyl chloride (2.97 g, 19.7 mmol, 1.1 equiv.) was added in a single portion to the stirred mixture. The reaction mixture was warmed up to 25 ˚C and stirred for 24 h. The product mixture was diluted with ethyl acetate (20.0 mL) and transferred to a separatory funnel. The diluted product mixture was washed with saturated aqueous sodium chloride (20.0 mL) and the aqueous layer was extracted into ethyl acetate (20.0 mL). The organic layers were combined, and the combined organic layers were dried over sodium sulfate. The dried solution was filtered, and the filtrate was concentrated to dryness. The crude product was purified by flash-column chromatography (eluting with 5% ethyl acetate–hexane) to afford the titled compound as a colorless oil (**S8**, 4.52 g, 90%).

$R_f$ = 0.25 (2% ethyl acetate–hexane; UV). $^1$H NMR (500 MHz, chloroform-*d*): δ 7.86 (d, *J* = 8.0 Hz, 1H, $H_4$), 6.75 – 6.64 (m, 2H, $H_3$, $H_5$), 3.85 (s, 3H, $H_6$), 2.56 (s, 3H, $H_7$), 0.98 (s, 9H, $H_1$), 0.22 (s, 6H, $H_2$). $^{13}$C NMR (126 MHz, chloroform-*d*): δ 167.8 (C), 159.1 (C), 143.1 (C), 132.9 (CH), 123.2 (CH), 122.5 (C), 117.3 (CH), 51.7 ($CH_3$), 25.8 ($CH_3$ x3), 22.3, 18.38, -4.21 ($CH_3$). IR (ATR-FTIR), cm$^{-1}$ 2952 (w), 2930 (w), 2858 (w), 1719 (m), 1600 (m), 1246 (s), 840 (m), 808 (m). HRMS-ESI (*m/z*): $[M+H]^+$ calculated for $C_{15}H_{24}O_3Si$, 281.1567; found, 281.1568.

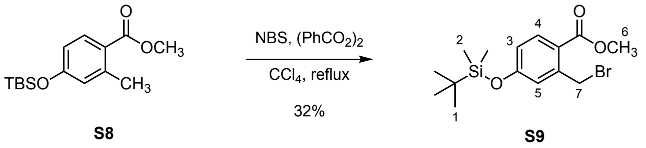


### Synthesis of S9:

*N*-Bromosuccinimide (762 mg, 4.28 mmol, 1.20 equiv.) and benzoyl peroxide (86.4 mg, 357 µmol, 0.10 equiv.) were added in sequence to a stirred solution of the compound **S8** (1.00 g, 3.57 mmol, 1.00 equiv.) in carbon tetrachloride (12.0 mL) at 25 ˚C. The reaction mixture was heated at reflux for 24 h. The product mixture was cooled down to 25 ˚C and filtrated through cotton. The filtrate was concentrated to dryness and purified by flash-column chromatography (eluting with 2% ethyl acetate–hexane) to afford the titled compound as a colorless oil (**S9**, 412 mg, 32%).

$R_f$ = 0.25 (2% ethyl acetate–hexane; UV). $^1$H NMR (500 MHz, chloroform-*d*) δ 7.91 (d, *J* = 8.5 Hz, 1H, $H_4$), 6.92 (d, *J* = 2.5 Hz, 1H, $H_5$), 6.79 (dd, *J* = 8.5, 2.5 Hz, 1H, $H_3$), 4.93 (s, 2H, $H_7$), 3.90 (s, 3H, $H_6$), 0.99 (s, 9H, $H_1$), 0.24 (s, 6H, $H_2$). $^{13}$C NMR (126 MHz, chloroform-*d*) δ 166.7 (C), 159.5 (C), 141.8 (C), 133.7 (CH), 123.4 (CH), 121.7 (C), 119.9 (CH), 52.1 ($CH_3$), 31.8 ($CH_2$), 25.7 ($CH_3$ x3), 18.4 (C), -4.2 ($CH_3$ x2). IR (ATR-FTIR), cm$^{-1}$ 2954 (w), 2930 (w), 2858 (w), 1719 (m), 1601 (m), 1307 (m), 1275 (m), 1253 (s), 843 (m), 783 (m). HRMS-ESI (*m/z*): $[M+H]^+$ calculated for $C_{15}H_{24}BrO_3Si$, 359.0673; found, 359.0671.

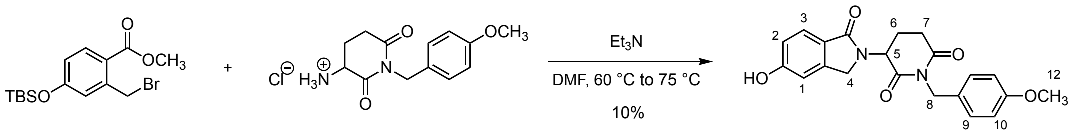


**Synthesis of S10:**

The compound **S6** (100 mg, 351 µmol, 1.00 equiv.) and triethylamine (147 µL, 1.05 µmol, 3.00 equiv.) were added in sequence to the stirred solution of the compound **S9** (126.2 mg, 351 µmol, 1.00 equiv.) in *N,N*-dimethylformamide (1.8 mL) at 25 ˚C. The reaction mixture was transferred to an oil bath and stirred at 60 ˚C for 12 h, and additionally stirred at 75 ˚C for 8 h. The product mixture was cooled down to 25 ˚C and diluted with ethyl acetate (5.0 mL). The diluted mixture was transferred to a separatory funnel and washed with 1*N*-hydrochloric acid (5.0 mL). The aqueous layer was extracted into ethyl acetate (5.0 mL). The organic layers were combined, and the combined organic layers were dried over sodium sulfate. The dried solution was filtered, and the filtrate was concentrated to dryness. The residue obtained was purified by flash-column chromatography (grading from 60% ethyl acetate–hexane to 90% ethyl acetate–hexane over 30 min) to afford the titled compound as a green solid (**S10**, 13.1 mg, 10%).

$R_f$ = 0.33 (90% ethyl acetate–hexane; UV). $^1$H NMR (500 MHz, chloroform-*d*): δ 7.65 (d, *J* = 8.0 Hz, 1H, $H_3$), 7.30 (d, *J* = 8.5 Hz, 2H, $H_{10}$), 6.90 (d, *J* = 8.0 Hz, 1H, $H_2$), 6.86 (s, 1H, $H_1$), 6.79 (d, *J* = 8.5 Hz, 2H, $H_9$), 5.12 (dd, *J* = 13.5, 5.0 Hz, 1H, $H_5$), 4.87 (s, 2H, $H_8$), 4.30 (d, *J* = 16.0 Hz, 1H, $H_4$), 4.21 (d, *J* = 16.0 Hz, 1H, $H_4$), 3.75 (s, 3H, $H_{12}$), 2.97 (ddd, *J* = 17.5, 4.5, 2.5 Hz, 1H, $H_7$), 2.82 (ddd, *J* = 17.5, 13.5, 5.5 Hz, 1H, $H_7$), 2.26 (dddd, *J* = 13.5, 13.5, 13.5, 4.5 Hz, 1H, $H_6$), 2.13 (dddd, *J* = 13.5, 5.0, 5.0, 2.5 Hz, 1H, $H_6$). $^{13}$C NMR (126 MHz, chloroform-*d*): δ 171.1 (C), 170.3 (C), 170.1 (C), 160.7 (C), 159.2 (C), 144.2 (C), 130.7 (CH x2), 129.1 (C), 125.8 (CH), 123.4 (C), 116.5 (CH), 114.0 (CH x2), 109.9 (CH), 55.4 ($CH_3$), 52.8 (CH), 47.3 ($CH_2$), 43.4 ($CH_2$), 32.3 ($CH_2$), 22.8 ($CH_2$). IR (ATR-FTIR), cm$^{-1}$ 2954 (w), 2930 (w), 2858 (w), 1719 (m), 1246 (s), 840 (s), 780 (s). HRMS-ESI (*m/z*): $[M+H]^+$ calculated for $C_{21}H_{21}N_2O_5$, 381.1445; found, 381.1445.

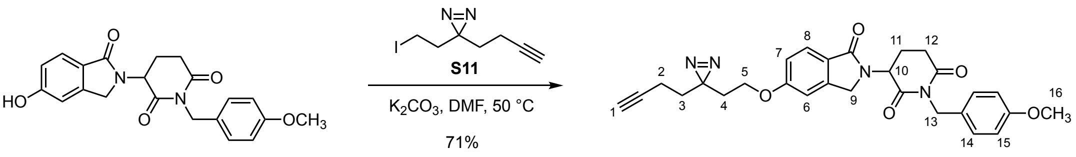


### Synthesis of S12:

Potassium carbonate (9.0 mg, 65.4 µmol, 3.00 equiv.) and the compound **S11**[2] (14.8 mg, 52.0 µmol, 2.40 equiv.) were added in sequence to a solution of the compound **S10** (8.3 mg, 21.8 µmol, 1.00 equiv.) in *N,N*-dimethylformamide (220 µL) at 25 ˚C. The reaction mixture was stirred for 48 h at 50 ˚C. The product mixture was diluted with ethyl acetate (5.0 mL) and transferred to a separatory funnel. The diluted product mixture was washed with water (5.0 mL) and the aqueous layer was extracted by ethyl acetate (3 x 5.0 mL). The organic layers were combined, and the combined organic layers were dried over sodium sulfate. The dried solution was filtered, and the filtrate was concentrated to dryness. The crude product was purified by flash-column chromatography (eluting with 50% ethyl acetate–hexane) to afford the titled compound as a colorless solid (**S12**, 7.7 mg, 71%).

$R_f$ = 0.25 (50% ethyl acetate–hexane; UV). $^1$H NMR (500 MHz, chloroform-*d*) δ 7.79 (d, *J* = 8.5 Hz, 1H, $H_8$), 7.33 (d, *J* = 8.5 Hz, 2H, $H_{14}$), 6.99 (dd, *J* = 8.5, 2.0 Hz, 1H, $H_7$), 6.92 (s, 1H, $H_6$), 6.81 (d, *J* = 8.5 Hz, 2H, $H_{15}$), 5.15 (dd, *J* = 13.5, 5.0 Hz, 1H, $H_{10}$), 4.89 (s, 2H, $H_{13}$), 4.37 (d, *J* = 16.0 Hz, 1H, $H_9$), 4.25 (d, *J* = 16.0 Hz, 1H, $H_9$), 3.88 (t, *J* = 6.0 Hz, 3H, $H_5$), 3.78 (s, 3H, $H_{16}$), 2.98 (ddd, *J* = 17.5, 4.5, 2.5 Hz, 1H, $H_{12}$), 2.84 (ddd, *J* = 17.5, 13.5, 5.5 Hz, 1H, $H_{12}$), 2.26 (dddd, *J* = 13.5, 13.5, 13.5, 4.5 Hz, 1H, $H_{11}$), 2.14 (dddd, *J* = 13.5, 5.5, 5.5, 2.5 Hz, 1H, $H_{11}$), 2.08 (d, *J* = 7.5, 2.5 Hz, 2H, $H_2$), 1.99 (t, *J* = 2.5 Hz, 1H, $H_1$), 1.94 (t, *J* = 6.0 Hz, 2H, $H_4$), 1.74 (t, *J* = 7.5 Hz, 2H, $H_3$). $^{13}$C NMR (126 MHz, chloroform-*d*) δ 171.1 (C), 170.2 (C), 169.3 (C), 162.1 (C), 159.2 (C), 143.9 (C), 130.8 (CH x2), 129.3 (C), 125.7 (CH), 124.7 (C), 115.4 (CH), 113.9 (CH x2), 108.6 (CH), 82.8 (C), 69.4 (CH), 63.2 ($CH_2$), 55.4 ($CH_3$), 52.6 (C), 47.0 ($CH_2$), 43.3 ($CH_2$), 32.9 ($CH_2$), 32.8 ($CH_2$), 32.4 ($CH_2$), 26.7 (C), 22.9 ($CH_2$), 13.4 ($CH_2$). IR (ATR-FTIR), $cm^{-1}$ 2920 (w), 1729 (s), 1230 (m), 1141 (m). HRMS-ESI (*m/z*): $[M+H]^+$ calculated for $C_{28}H_{29}N_4O_5$, 501.2132; found, 501.2140.

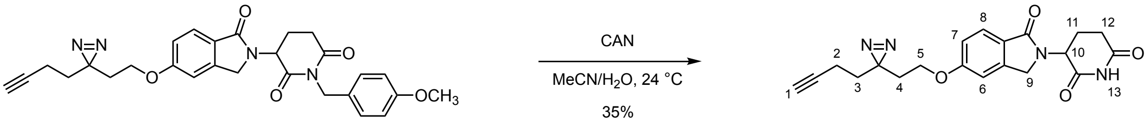


**S12** **2**

### Synthesis of analog 2:

Ammonium cerium(IV) nitrate (32.8 mg, 61.6 µmol, 4.00 equiv.) was added in a single portion to a solution of the compound **S12** (7.7 mg, 30.0 µmol, 1.00 equiv.) in acetonitrile (170 µL) and water (25.0 µL) at 25 ˚C. The reaction mixture was stirred for 6 h at 25 ˚C. The product mixture was diluted with ethyl acetate (5.0 mL) and transferred to a separatory funnel. The diluted product mixture was washed with water (5.0 mL) and saturated aqueous sodium chloride (5.0 mL). The combined aqueous layers were extracted with chloroform (3 x 5.0 mL). The organic layers were combined, and the combined organic layers were dried over sodium sulfate. The dried solution was filtered, and the filtrate was concentrated to dryness. The crude product was purified by flash-column chromatography (eluting with 1% methanol–dichloromethane). The product mixture was further purified by recrystallization from dichloromethane to afford the titled compound as a colorless solid (**2**, 2.1 mg, 35%).

$R_f$ = 0.25 (2% methanol–dichloromethane; UV). $^1$H NMR (500 MHz, chloroform-*d*) δ 7.89 (s, 1H, $H_{13}$), 7.80 (d, *J* = 8.4 Hz, 1H, $H_8$), 7.00 (dd, *J* = 8.4, 2.2 Hz, 1H, $H_7$), 6.94 (d, *J* = 2.2 Hz, 1H, $H_{16}$), 5.21 (dd, *J* = 13.2, 5.2 Hz, 1H, $H_{10}$), 4.45 (d, *J* = 15.8 Hz, 1H, $H_9$), 4.29 (d, *J* = 15.8 Hz, 1H, $H_9$), 3.88 (t, *J* = 6.2 Hz, 2H, $H_5$), 2.94-2.91 (m, 1H, $H_{12}$), 2.83 (ddd, *J* = 17.9, 13.2, 5.2 Hz, 1H, $H_{12}$), 2.35 (dddd, *J* = 13.2, 13.2, 13.2, 4.8 Hz, 1H, $H_{11}$), 2.22 (dddd, *J* = 13.2, 5.2, 5.2, 2.2 Hz, 1H, $H_{11}$), 2.08 (td, *J* = 7.5, 2.6 Hz, 2H, $H_2$), 2.00 (t, *J* = 2.6 Hz, 1H, $H_1$), 1.95 (t, *J* = 6.2 Hz, 2H, $H_4$), 1.75 (t, *J* = 7.5 Hz, 2H, $H_3$). $^{13}$C NMR (126 MHz, chloroform-*d*) δ 171.1 (C), 169.7 (C), 169.2 (C), 162.1 (C), 143.9 (C), 125.7 (CH), 124.5 (C), 115.5 (CH), 108.6 (CH), 82.8 (C), 69.5 (CH), 63.2 ($CH_2$), 52.0 (CH), 47.0 ($CH_2$), 32.9 ($CH_2$), 32.8 ($CH_2$), 31.7 ($CH_2$), 26.7 (C), 23.6 ($CH_2$), 13.4 ($CH_2$). IR (ATR-FTIR), 3285 (br), 3100 (br), 2924 (br), 1682 (s), 1614 (s), 1454 (m), 1368 (m), 1258 (s), 1233 (s), 1199 (s), 1178 (s), 770 (w), 687 (w) $cm^{-1}$. HRMS-ESI (*m/z*): $[M+H]^+$ calculated for $C_{20}H_{21}N_4O_4$, 381.1557; found, 381.1560.

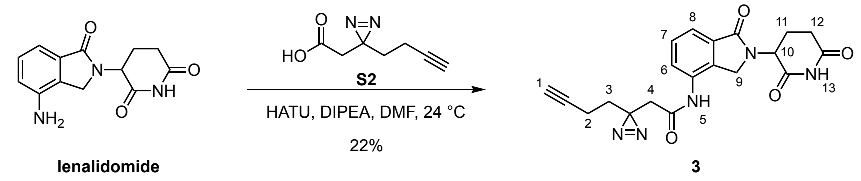

### Synthesis of analog 3:

Lenalidomide (14.2 mg, 54.8 μmol, 1.00 equiv.), sodium sulfate (23.3 mg, 164 μmol, 3.00 equiv.), HATU (75.0 mg, 197 μmol, 3.60 equiv.) and *N,N*-diisopropylethylamine (26.0 μL, 197 μmol, 3.60 equiv.) were added in a single portion to a stirred solution of the carboxylic acid **S2** (10.0 mg, 65.7 μmol, 1.20 equiv.) sequentially at 25 ˚C. The reaction mixture was stirred for 24 h at 25 ˚C. The product mixture was diluted with ethyl acetate (5.0 mL) and transferred to a separatory funnel. The diluted product mixture was washed with saturated aqueous sodium chloride (5.0 mL) and 10% aqueous citric acid (5.0 mL). The layers were mixed, and the aqueous layer was extracted by ethyl acetate (3 x 5 mL). The combined organic layers were dried over sodium sulfate and concentrated to dryness. The crude product was purified by flash-column chromatography (eluting with 2% methanol–dichloromethane, grading to 4% methanol–dichloromethane, three steps). The obtained product mixture was further purified by HPLC (grading 95% to 5% water-acetonitrile over 25 min, flow rate = 10 mL/min, retention time = 11.8 min), to afford the titled compound as a white solid (**3**, 4.6 mg, 22%).

$R_f$ = 0.25 (4% methanol-dichloromethane; UV). $^1$H NMR (500 MHz, DMSO-$d_6$) δ 11.03 (s, 1H, $H_{13}$), 9.90 (s, 1H, $H_5$), 7.79 (dd, *J* = 7.6, 1.4 Hz, 1H, $H_6$/$H_8$), 7.57 – 7.47 (m, 2H, $H_6$/$H_8$, $H_7$), 5.15 (dd, *J* = 13.5, 5.0 Hz, 1H, $H_{10}$), 4.37 (d, *J* = 17.5 Hz, 1H, $H_9$), 4.31 (d, *J* = 17.5 Hz, 1H, $H_9$), 2.92 (ddd, *J* = 17.4, 13.5, 5.0 Hz, 1H, $H_{12}$), 2.85 (t, *J* = 2.6 Hz, 1H, $H_1$), 2.61 (ddd, *J* = 17.4, 5.0, 2.2 Hz, 1H, $H_{12}$), 2.54 (s, 2H, $H_4$), 2.34 (dddd, *J* = 13.5, 13.5, 13.5, 4.6 Hz, 1H, $H_{11}$), 2.11 – 1.99 (m, 3H, $H_{11}$, $H_2$), 1.73 (t, *J* = 7.4 Hz, 2H, $H_3$). $^{13}$C NMR (126 MHz, DMSO-$d_6$): δ. IR (ATR-FTIR), cm$^{-1}$ 3272 (br), 2920 (m), 2851 (m), 1686 (s), 1677 (s), 1432 (m), 1418 (m), 1235 (m), 1204 (m), 750 (m). HRMS-ESI (*m/z*): $[M+H]^+$ calculated for $C_{20}H_{20}N_5O_4$ 394.1510; found, 394.1505.

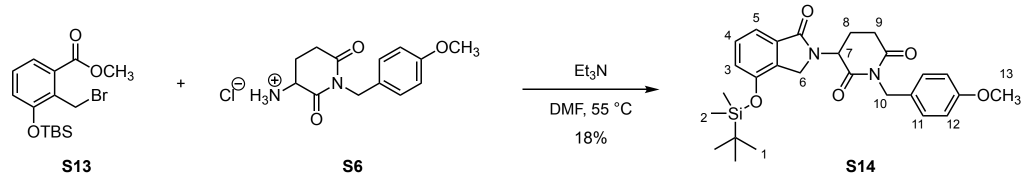


### Synthesis of S14:

The compound **S6** (159 mg, 557 µmol, 1.00 equiv.) and triethylamine (232 µL, 1.67 mmol, 3.00 equiv.) were added in sequence to a stirred solution of the compound **S13**[10] (200 mg, 557 µmol, 1.00 equiv.) in *N,N*-dimethylformamide (2.8 mL) at 25 ˚C. The reaction mixture was stirred for 24 h at 55 ˚C. The product mixture was diluted with ethyl acetate (10.0 mL) and transferred to a separatory funnel. The diluted product mixture was washed with 1*N* hydrochloric acid (5.0 mL) and the aqueous layer was extracted by ethyl acetate (3 x 5.0 mL). The organic layers were combined, and the combined organic layers were dried over sodium sulfate. The dried solution was filtered, and the filtrate was concentrated to dryness. The crude product was purified by flash-column chromatography (eluting with 50% ethyl acetate–hexane) to afford the titled compound as a white solid (**S14**, 50.3 mg, 18%).

$R_f$ = 0.25 (40% ethyl acetate–hexane; UV). $^1$H NMR (500 MHz, chloroform-*d*): δ 7.49 (dd, *J* = 8.0, 0.5 Hz, 1H, $H_5$), 7.39 – 7.30 (m, 3H, $H_4$, $H_{11}$), 6.96 (dd, *J* = 8.0, 0.5 Hz, 1H, $H_3$), 6.81 (d, *J* = 8.5 Hz, 2H, $H_{12}$), 5.17 (dd, *J* = 13.5, 5.0 Hz, 1H, $H_7$), 4.93 (d, *J* = 14.0 Hz, 1H, $H_{10}$), 4.89 (d, *J* = 14.0 Hz, 1H, $H_{10}$), 4.31 (d, *J* = 16.0 Hz, 1H, $H_6$), 4.21 (d, *J* = 16.0 Hz, 1H, $H_6$), 3.78 (s, 3H, $H_{13}$), 2.98 (ddd, *J* = 17.5, 4.5, 2.5 Hz, 1H, $H_9$), 2.84 (ddd, *J* = 17.5, 13.5, 5.0, 1H, $H_9$), 2.33 (dddd, *J* = 13.5, 13.5, 13.5, 5.0 Hz, 1H, $H_8$), 2.15 (dddd, *J* = 13.5, 5.0, 5.0, 2.5 Hz, 1H, $H_8$), 1.00 (s, 9H, $H_1$), 0.26 (s, 6H, $H_2$). $^{13}$C NMR (126 MHz, chloroform-*d*): δ 171.1 (C), 170.1 (C), 169.6 (C), 159.2 (C), 151.0 (C), 133.7 (C), 132.1 (C), 130.8 (CH x2), 129.8 (CH), 129.3 (C), 121.7 (CH), 117.1 (CH), 113.9 (CH x2), 55.4 ($CH_3$), 52.7 (CH), 45.3 ($CH_2$), 43.3 ($CH_2$), 32.4 ($CH_2$), 25.8 ($CH_3$ x3), 22.8 ($CH_2$), 18.3 (C), -4.06 ($CH_3$), -4.09 ($CH_3$). IR (ATR-FTIR), cm$^{-1}$ 2054 (w), 2930 (w), 1678 (s), 1288 (m), 1248 (m), 831 (m). HRMS-ESI (*m/z*): $[M+H]^+$ calculated for $C_{27}H_{35}N_2O_5Si$, 495.2310; found, 495.2315.

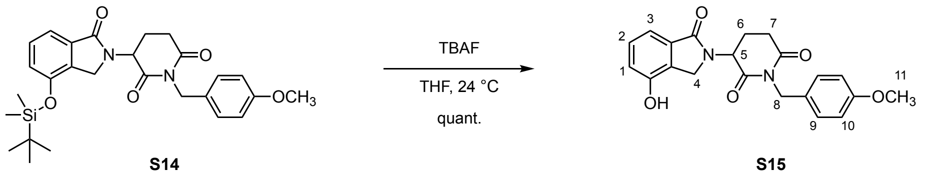

### Synthesis of S15:

Tetrabutylammonium fluoride solution in tetrahydrofuran (1.0 M, 153 µL, 153 mmol, 1.50 equiv.) was added dropwise to a stirred solution of the compound **S14** (50.3 mg, 102 µmol, 1.00 equiv.) in tetrahydrofuran (510 µL) over 5 min at 0 ˚C. The reaction mixture was stirred for 1 h at 25 ˚C. The product mixture was diluted with ethyl acetate (5.0 mL) and saturated aqueous ammonium chloride solution (5.0 mL). The biphasic product mixture was transferred to a separatory funnel and the layers were separated. The aqueous layer was extracted by ethyl acetate (3 x 5 mL). The organic layers were combined, and the combined organic layers were dried over sodium sulfate. The dried solution was filtered, and the filtrate was concentrated to dryness. The crude product was purified by flash-column chromatography (eluting with 60% ethyl acetate–hexane, grading to 80% ethyl acetate–hexane, three steps) to afford the titled compound as a colorless solid (**S15**, 38.6 mg, quantitative yield).

$R_f$ = 0.25 (70% ethyl acetate–hexane; UV). $^1$H NMR (500 MHz, chloroform-*d*) δ 7.43 (d, *J* = 8.0, 1H, $H_3$), 7.32 (d, *J* = 8.5, 2H, $H_9$), 7.29 (dd, *J* = 8.0, 1H, $H_2$), 6.95 (d, *J* = 8.0 Hz, 1H, $H_1$), 6.81 (d, *J* = 8.5 Hz, 2H, $H_{10}$), 5.16 (dd, *J* = 13.5, 5.0 Hz, 1H, $H_5$), 4.89 (s, 2H, $H_8$), 4.38 (d, *J* = 16.0 Hz, 1H, $H_4$), 4.28 (d, *J* = 16.0 Hz, 1H, $H_4$), 3.77 (s, 3H, $H_{11}$), 2.99 (ddd, *J* = 18.0, 4.5, 2.5 Hz, 1H, $H_7$), 2.84 (ddd, *J* = 18.0, 13.5, 5.5 Hz, 1H, $H_7$), 2.29 (dddd, *J* = 13.5, 13.5, 13.5, 4.5 Hz, 1H, $H_6$), 2.15 (dddd, *J* = 13.5, 13.5, 5.5, 2.5 Hz, 1H, $H_7$). $^{13}$C NMR (126 MHz, chloroform-*d*) δ 171.1 (C), 170.2 (C), 169.8 (C), 159.2 (C), 151.3 (C), 133.5 (C), 130.7 (CH x2), 130.0 (CH), 129.2 (C), 128.0 (C), 118.6 (CH), 116.4 (CH), 114.0 (CH x2), 55.4 ($CH_3$), 52.9 (CH), 45.3 ($CH_2$), 43.4 ($CH_2$), 32.4 ($CH_2$), 22.8 ($CH_2$). IR (ATR-FTIR), cm$^{-1}$ 3171 (br), 2958 (w), 2924 (w), 2853 (w), 1678 (s), 1608 (m), 1513 (m), 1291 (m), 1248 (m), 1164 (m), 733 (m). HRMS-ESI (*m/z*): $[M+H]^+$ calculated for $C_{21}H_{21}N_2O_5$, 381.1445; found, 381.1442.

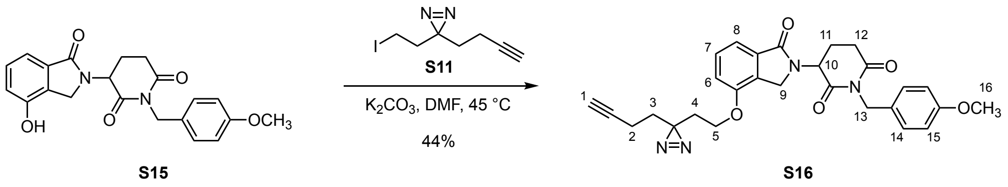

### Synthesis of S16:

Potassium carbonate (42.1 mg, 304 µmol, 3.00 equiv.) and minimalist iodide **S11**[2] (25.7 mg, 101 µmol, 1.00 equiv.) were added in sequence to a solution of the compound **S15** (38.6 mg, 101 µmol, 1.00 equiv.) in *N,N*-dimethylformamide (500 µL) at 25 ˚C. The reaction mixture was stirred for 24 h at 45 ˚C. The product mixture was diluted with ethyl acetate (10.0 mL) and transferred to a separatory funnel. The diluted product mixture was washed with water (5.0 mL) and the aqueous layer was extracted by ethyl acetate (3 x 5.0 mL). The organic layers were combined, and the combined organic layers were dried over sodium sulfate. The dried solution was filtered, and the filtrate was concentrated to dryness. The crude product was purified by flash-column chromatography (eluting with 50% ethyl acetate–hexane, grading to 60% ethyl acetate–hexane, one step) to afford the titled compound as a colorless solid (**S16**, 16.8 mg, 44%).

$R_f$ = 0.25 (60% ethyl acetate–hexane; UV). $^1$H NMR (500 MHz, chloroform-*d*): δ 7.50 (d, *J* = 8.0 Hz, 1H, $H_6$/$H_8$), 7.42 (t, *J* = 8.0 Hz, 1H, $H_7$), 7.34 (d, *J* = 8.5 Hz, 2H, $H_{14}$), 6.97 (d, *J* = 8.0 Hz, 1H, $H_6$/$H_8$), 6.81 (d, *J* = 8.5 Hz, 2H, $H_{15}$), 5.18 (dd, *J* = 13.5, 5.0 Hz, 1H, $H_{10}$), 4.90 (s, 2H, , $H_{13}$), 4.41 (d, *J* = 16.5 Hz, 1H, $H_9$), 4.32 (d, *J* = 16.5 Hz, 1H, $H_9$), 3.98-3.94 (m, 2H, $H_5$), 3.77 (s, 3H, $H_{16}$), 2.99 (ddd, *J* = 17.5, 4.5, 2.5 Hz, 1H, $H_{12}$), 2.85 (ddd, *J* = 17.5, 13.5, 5.5 Hz, 1H, , $H_{12}$), 2.34 (dddd, *J* = 13.5, 13.5, 13.5, 4.5 Hz, 1H, $H_{11}$), 2.16 (dddd, *J* = 13.5, 4.5, 2.5 Hz, 1H, $H_{11}$), 2.06 (td, *J* = 7.5, 2.6 Hz, 2H, $H_2$), 2.00 – 1.87 (m, 3H, $H_1$, $H_4$), 1.72 (t, *J* = 7.4 Hz, 2H, $H_3$). $^{13}$C NMR (126 MHz, chloroform-*d*): δ 171.1 (C), 170.0 (C), 169.5 (C), 159.2(C), 153.6 (C), 133.5 (C), 130.8 (CH x2), 130.1 (C), 130.0 (CH), 129.2 (C), 116.7 (CH), 113.9 (CH x2, CH), 82.8 (C), 69.5 (CH), 63.1 ($CH_2$), 55.4 ($CH_3$), 52.7 (CH), 45.2 ($CH_2$), 43.3 ($CH_2$), 32.9 ($CH_2$), 32.7 ($CH_2$), 32.4 ($CH_2$), 26.7 (C), 22.9 ($CH_2$), 13.4 ($CH_2$). IR (ATR-FTIR), cm$^{-1}$: 3287 (br), 2923 (m)m, 2853 (m), 1677 (s), 1606 (m), 1513 (m), 1275 (m), 1246 (m), 1161 (m), 749 (m). HRMS-ESI (*m/z*): $[M+H]^+$ calculated for $C_{28}H_{28}N_4O_5$, 501.2132; found, 501.2077.

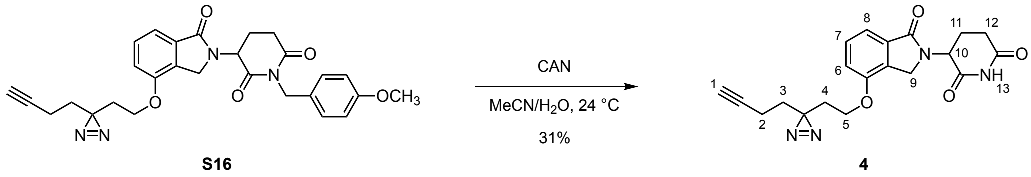

### Synthesis of analog 4:

Ammonium cerium(IV) nitrate (65.7 mg, 120 µmol, 4.00 equiv.) was added in a single portion to a solution of the compound **S16** (15.0 mg, 30.0 µmol, 1.00 equiv.) in acetonitrile (330 µL) and water (67.0 µL) at 25 ˚C. The reaction mixture was stirred for 2 h at 25 ˚C. The product mixture was diluted with ethyl acetate (5.0 mL) and transferred to a separatory funnel. The diluted product mixture was washed with water (5.0 mL) and saturated aqueous sodium chloride (5.0 mL). The organic layers were combined, and the combined organic layers were dried over sodium sulfate. The dried solution was filtered, and the filtrate was concentrated to dryness. The crude product was purified by flash-column chromatography (eluting with 1% methanol–dichloromethane). The product mixture was further purified by HPLC (grading 95% to 5% water–acetonitrile over 25 min, flow rate = 10 mL/min, retention time = 14.6 min) to afford the titled compound as a colorless solid (**4**, 2.8 mg, 31%).

$R_f$ = 0.25 (2% methanol–dichloromethane; UV). $^1$H NMR (600 MHz, chloroform-*d*) δ 7.97 (s, 1H, $H_{13}$), 7.51 (d, *J* = 7.8 Hz, 1H, $H_6$/$H_8$), 7.44 (dd, *J* = 7.8, 7.8 Hz, 1H, $H_7$), 6.98 (d, *J* = 7.8 Hz, 1H, $H_6$/$H_8$), 5.23 (dd, *J* = 13.2, 5.2 Hz, 1H, $H_{10}$), 4.48 (d, *J* = 16.4 Hz, 1H, $H_9$), 4.36 (d, *J* = 16.4 Hz, 1H, $H_9$), 3.96 (ddd, *J* = 9.5, 6.6, 5.2 Hz, 1H, $H_5$), 3.92 (ddd, *J* = 9.5, 7.2, 4.8 Hz, 1H, $H_5$), 2.97 – 2.89 (m, 1H, $H_{12}$), 2.84 (ddd, *J* = 18.1, 13.2, 5.2 Hz, 1H, $H_{12}$), 2.42 (dddd, *J* = 13.2, 13.2, 13.2, 4.7 Hz, 1H, $H_{11}$), 2.23 (dddd, *J* = 13.2, 5.2, 5.2, 2.5 Hz, 1H, $H_{11}$), 2.07 (td, *J* = 7.5, 2.4 Hz, 2H, $H_2$), 1.98 (t, *J* = 2.4 Hz, 1H, $H_1$), 1.97 (ddd, *J* = 15.6, 7.2, 5.2 Hz, 1H, $H_4$), 1.92 (ddd, *J* = 15.6, 6.6, 4.8 Hz, 1H, $H_4$), 1.73 (t, *J* = 7.5 Hz, 2H, $H_3$). $^{13}$C NMR (126 MHz, chloroform-*d*) δ 171.0 (C), 169.4 (C), 169.3 (C), 153.4 (C), 133.2 (C), 129.9 (C), 129.9 (CH), 116.6 (CH), 113.9 (CH), 82.7 (C), 69.4 (CH), 62.9 ($CH_2$), 51.9 (CH), 45.0 ($CH_2$), 32.7 ($CH_2$), 32.6 ($CH_2$), 31.6 ($CH_2$), 26.6 (C), 23.4 ($CH_2$), 13.3 ($CH_2$). IR (ATR-FTIR), cm$^{-1}$ 3243 (br), 2925 (br), 1694 (s), 1606 (m), 1493 (m), 1277 (m), 1145 (m), 750 (m). HRMS-ESI (*m/z*): $[M+H]^+$ calculated for $C_{20}H_{21}N_4O_4$, 381.1557; found, 381.1566.

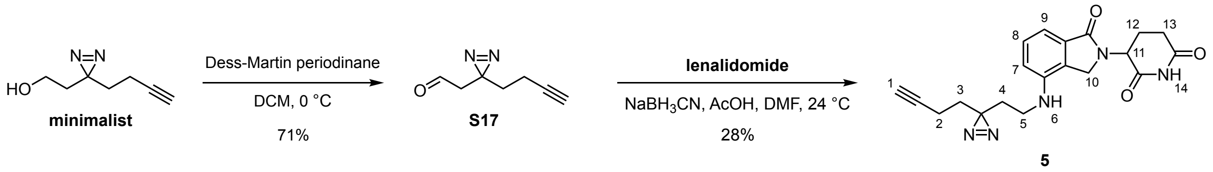


### Synthesis of analog 5:

*Step 1*
Sodium bicarbonate (164.2 mg, 1.95 mmol, 15.0 equiv.) was added in a single portion to a stirred solution of minimalist[2] (18.0 mg, 130 µmol, 1.00 equiv.) in dichloromethane (1.3 mL). The reaction mixture was cooled down to 0 ˚C and Dess-Martin periodinane (82.9 mg, 195 µmol, 1.5 equiv.) was added to the solution in a single portion. The reaction mixture was stirred for 2 h at 0 ˚C. Additional Dess-Martin periodinane (26.7 mg, 65 µmol, 0.5 equiv.) was added to the reaction mixture in a single portion at 0 ˚C and the reaction mixture was stirred for 30 min at 0 ˚C. Saturated aqueous sodium thiosulfate (5.0 mL) was added to the product mixture at 0 ˚C and stirred for 30 min at 25 ˚C. The product mixture was transferred to a separatory funnel and washed with water (5.0 mL). The aqueous layer was extracted with diethyl ether (3 x 5.0 mL). The organic layers were combined, and the combined organic layers were dried over sodium sulfate. The crude product was filtrated through silica gel (eluting with 50% diethyl ether–hexane) to remove residual reagents. The aldehyde **S17** was afforded as colorless oil (12.6 mg, 71%). The product was used in the following step without further purification.

*Step 2*
Lenalidomide (24.0 mg, 92.5 µmol, 1.00 equiv.) and acetic acid (6.4 µL, 111 µmol, 1.20 equiv.) were added in a single portion to a stirred solution of the aldehyde **S17** (12.6 mg, 92.5 µmol, 1.00 equiv.) in *N,N*-dimethylformamide (310 µL) at 25 ˚C. The reaction mixture was stirred for 20 min at 25 ˚C. Sodium cyanoborohydride (8.7 mg, 139 µmol, 1.50 equiv.) was added in a single portion to the reaction mixture and the reaction mixture was stirred for an additional 24 h at 25 °C. The product mixture was diluted with ethyl acetate (5.0 mL) and transferred to a separatory funnel. The diluted product mixture was washed with saturated aqueous sodium bicarbonate (5.0 mL) and saturated aqueous sodium chloride (5.0 mL). The layers were mixed, and the aqueous layer was extracted by ethyl acetate (3 x 5 mL). The organic layers were combined, and the combined organic layers were dried over sodium sulfate. The dried solution was filtered, and the filtrate was concentrated to dryness. The crude product was purified by flash-column chromatography (eluting with 2% methanol–dichloromethane, grading to 3% methanol–dichloromethane, two steps). The obtained product mixture was further purified by HPLC (grading 95% to 48% water–acetonitrile over 28 min, flow rate = 10 mL/min, retention time = 25.2 min), to afford the titled compound as a colorless solid (**5**, 9.8 mg, 28%).

$R_f$ = 0.30 (4% methanol–dichloromethane; UV, anis aldehyde). $^1$H NMR (500 MHz, methanol-$d_4$): δ 7.34 (dd, $J$ = 7.5, 7.5 Hz, 1H, $H_8$), 7.10 (d, $J$ = 7.5 Hz, 1H, $H_9$), 6.79 (d, $J$ = 7.5 Hz, 1H, $H_7$), 5.16 (dd, $J$ = 13.5, 5.0 Hz, 1H, $H_{11}$), 4.33 (d, $J$ = 17.0 Hz, 1H, $H_{10}$), 4.28 (d, $J$ = 17.0 Hz, 1H, $H_{10}$), 3.08 (t, $J$ = 7.0 Hz, 2H, $H_5$), 2.92 (ddd, $J$ = 17.5, 13.5, 5.5 Hz, 1H, $H_{13}$), 2.79 (ddd, $J$ = 17.5, 4.5, 2.5 Hz, 1H, $H_{13}$), 2.48 (dddd, $J$ = 13.5, 13.5, 13.5, 4.5 Hz, 1H, $H_{12}$), 2.29 (t, $J$ = 2.5 Hz, 1H, $H_1$), 2.23 – 2.13 (m, 1H, $H_{12}$), 2.03 (td, $J$ = 7.5, 2.5 Hz, 2H, $H_2$), 1.80 (t, $J$ = 7.0 Hz, 2H, $H_4$), 1.65 (t, $J$ = 7.5 Hz, 2H, $H_3$). $^{13}$C NMR (126 MHz, methanol-$d_4$): δ 174.7 (C), 172.4 (C), 172.3 (C), 144.5 (CH), 133.1 (C), 130.7 (CH), 128.4 (C), 113.8 (CH), 112.4 (CH), 83.7 (C), 70.4 (CH), 53.6 (CH), 47.3 ($CH_2$), 39.0 ($CH_2$), 33.6 ($CH_2$), 33.0 ($CH_2$), 32.4 ($CH_2$), 28.1 (C), 24.3 ($CH_2$), 13.8 ($CH_2$). IR (ATR-FTIR), cm$^{-1}$: 3341 (br), 3290 (br), 2920 (br), 2854 (br), 1672 (s), 1607 (s), 1349 (m), 1201 (m), 748 (m). HRMS-ESI (*m/z*): $[M+Na]^+$ calculated for $C_{20}H_{21}N_5NaO_3$, 402.1536; found, 402.1531.

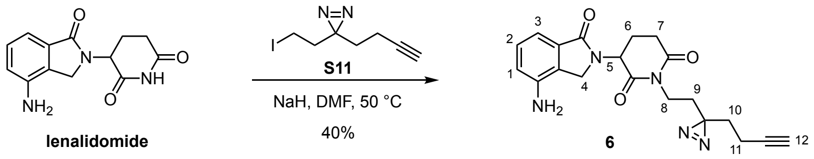


### Synthesis analog 6:

Sodium hydride (3.5 mg, 146 µmol, 1.50 equiv.) was added in a single portion to a stirred solution of lenalidomide (25.2 mg, 97.0 µmol, 1.00 equiv.) in *N,N*-dimethylformamide (770 µL) at 0 ˚C. The reaction mixture was stirred for 1 h at 25 ˚C. A solution of minimalist iodide **S11** (28.9 mg, 117 µmol, 1.20 equiv.) in *N,N*-dimethylformamide (200 µL) was added to the reaction mixture and the reaction mixture was stirred for an additional 24 h at 50 °C. The product mixture was cooled over 30 min to 25 °C. The cooled mixture was diluted with ethyl acetate (5.0 mL) and transferred to a separatory funnel. The mixture was washed with saturated aqueous sodium chloride (5.0 mL) and the aqueous layer was extracted by ethyl acetate (3 x 5.0 mL). The organic layers were combined, and the combined organic layers were dried over sodium sulfate. The dried solution was filtered, and the filtrate was concentrated to dryness. The crude product was purified by flash-column chromatography (eluting with 50% ethyl acetate–hexanes, grading to 80% ethyl acetate–hexanes, two steps). The product was further purified with HPLC (eluting with 5% acetonitrile–water, grading to 95% acetonitrile–water over 25 min, flow rate = 10 mL/min, retention time = 14.0 min) afford the titled compound as a white solid (**6**, 14.8 mg, 40%).

$R_f$ = 0.24 (80% ethyl acetate–hexane; UV, ninhydrin). $^1$H NMR (500 MHz, methanol-$d_4$) δ 7.26 (dd, *J* = 7.5, 7.5 Hz, 1H, $H_2$), 7.13 (dd, *J* = 7.5, 1.0 Hz, 1H, $H_3$), 6.92 (dd, *J* = 7.5, 1.0 Hz, 1H, $H_1$), 5.16 (dd, *J* = 13.5, 5.0 Hz, 1H, , $H_5$), 4.34 (d, *J* = 16.5 Hz, 1H, $H_4$), 4.27 (d, *J* = 16.5 Hz, 1H, $H_4$), 3.86 (dt, *J* = 13.5, 7.5 Hz, 1H, $H_8$), 3.81 (dt, *J* = 13.5, 7.5 Hz, 1H, $H_8$), 3.02 – 2.88 (m, 2H, $H_7$), 2.48 (dddd, *J* = 13.0, 13.0, 13.0, 5.5 Hz, 1H, $H_6$), 2.27 (t, *J* = 2.5 Hz, 1H, $H_{12}$), 2.18 (dddd, *J* = 13.0, 13.0, 5.0, 3.0 Hz, 1H, $H_6$), 2.03 (dt, *J* = 7.5, 2.5 Hz, 2H, $H_{11}$), 1.68 (dt, *J* = 15.0, 7.5 Hz, 1H, $H_{10}$), 1.65 (dt, *J* = 15.0, 7.5 Hz, 1H, $H_{10}$), 1.60 (ddd, *J* = 14.5, 7.5, 7.5 Hz, 1H, $H_9$), 1.56 (ddd, *J* = 14.5, 7.5, 7.5 Hz, 1H, $H_9$). $^{13}$C NMR (125 MHz, chloroform-*d*): δ 171.1 (C), 170.2 (C), 169.9 (C), 141.3 (C), 132.4 (C), 129.7 (CH), 126.4 (C), 118.3 (CH), 114.6 (CH), 82.8 (C), 69.5 (CH), 52.7 (CH), 45.1 ($CH_2$), 36.1 ($CH_2$), 32.3 ($CH_2$), 31.6 ($CH_2$), 31.5 ($CH_2$), 27.2 (C), 22.8 ($CH_2$), 13.4 ($CH_2$). IR (ATR-FTIR), cm$^{-1}$: 3352 (br), 3290 (br), 2921 (br), 1672 (s), 1491 (m), 1349 (m), 1146 (m), 750 (m). HRMS-ESI (*m/z*): $[M+Na]^+$ calculated for $C_{20}H_{21}N_5NaO_3$, 402.1537; found, 402.1539.

**Antibodies and biological reagents.**

Antibodies included anti-IKZF1 (Cell Signaling Technology, Cat # 9034S), anti-IKZF3 (Novus Bio, Cat # NBP2-16938), anti-CRBN (Cell Signaling Technology, Cat # 71810S, Sigma-Aldrich, Cat # HPA045910), anti-Flag (Sigma-Aldrich, Cat # F3165), anti-β-actin (Santa Cruz, Cat # sc-47778), anti-GSPT1 (Proteintech, Cat # 10763-1-AP), anti-CK1α (Abcam, Cat # ab206652), anti-eIF3i (Proteintech, Cat # 10086-380), anti-DDB1 (Bethyl Laboratories, Cat # A300-462A), anti-CUL4A (Bethyl Laboratories, Cat # A300-739A).

Antibody-conjugated beads for immunoprecipitation included Anti-Flag M2 magnetic beads (Sigma-Aldrich, Cat # M8823), anti-EPEA CaptureSelect™ C-tag affinity matrix (Thermo Scientific, Cat # 191307005), and Pierce Anti-HA magnetic beads (Pierce, Cat # PI88836). Protein A/G magnetic beads (Bimake, Cat # B23202) were used for endogenous protein immunoprecipitation. For flow cytometry staining, PE rat anti-human IL-2 antibody (BD Biosciences, 559334), FITC mouse anti-human CD69 antibody (BD Biosciences, 555530, Clone FN50) and FITC rat anti-mouse TNF-α antibody (eBioscience™, MP6-XT22) were used. Cell lysis buffer was prepared by diluting from 10X cell lysis buffer (Cell Signaling Technology, Cat # 9803S). TBS buffer was prepared by diluting from 10X TBS buffer solution (Quality Biological, Cat # 10128-546). Sequencing-grade modified trypsin (Promega, Cat # V5111), sequencing-grade chymotrypsin (Promega, Cat # V1061) and mini Bio-Spin columns (Bio-Rad, Cat # 7326207) were purchased. For qRT-PCR, RNeasy Plus Mini kit (Qiagen, Cat # 74134), PrimeScript RT reagent kit (TakaraBio Cat # RR037A), and QuantiTect SYBR Green PCR kit (Qiagen, Cat # 204143) were used.

**Cell culture and transfection.**

HEK293T cells (a gift from the Bertozzi lab, Stanford University), HEK293T cells stably expressing Flag-CRBN (HEK293T-CRBN) and HEK293T cells with CRBN knock-down by lentiviral shRNA (a gift from the Deshaies lab, California Institute of Technology), HEK293T cells stably transfected with NLucP/NF-κB vector (HEK293T-NF-κB, a gift from the Choudhary lab, Broad Institute), and RAW264.7 cells (a gift from the Mitchison lab, Harvard Medical School) were cultured in Dulbecco's Modified Eagle Medium (DMEM, Cat # 11995073), supplemented with penicillin (50 μg/mL), streptomycin (50 μg/mL), and 10% (v/v) FBS. HEK293T-CRBN cell line was established by transducing cells with the corresponding lentiviral particles with transduction efficiency monitored by GFP signals via flow cytometry. HEK293T-NF-κB were maintained under the selection of 1 μg/μL hygromycin. MM.1S cells (ATCC), U937 cells (ATCC), Jurkat cells (ATCC) and MOLM13 cells (a gift from the Shair lab, Harvard University) were cultured in RPMI1640 (Cat #45000-396) supplemented with penicillin (50 μg/mL), streptomycin (50 μg/mL) and 10% (v/v) FBS.

Transfection was performed using TransIT-PRO (Mirus Bio, Cat # MIR 5740) according to the manufacturer's instructions. Cells were lysed with 1X cell lysis buffer diluted from 10X cell lysis buffer supplemented with 1X cOmplete protease inhibitor cocktail (PI, Cat # 11697498001) unless otherwise noted.

**Plasmids and subcloning.**

Human CRBN with a Flag tag, human eIF3i with a His tag, and HA-tagged ubiquitin were obtained as detailed in the table below. IKZF1 with an EPEA tag was subcloned into the pcDNA3.1 vector. Empty pcDNA3.1 vector was used as a negative control for transfection.

Genetic constructs and primers used in this study are listed below:

| No. | Plasmid name | Abbreviated name | Details | Source |
|---|---|---|---|---|
| 1 | pcDNA3.1 | Ctrl | Empty vector for mammalian expression | Invitrogen, Cat # V79020 |
| 2 | pCDH-Flag-CRBN-WT-T2AcGFP-MSCV | Flag-CRBN | Full length CRBN with a Flag tag | A gift from Deshaies lab |
| 3 | pcDNA3.1-IKZF1-EPEA | IKZF1-EPEA | | Made in house |
| 4 | pNL3.2.NF-κB-RE[NlucP/NF-κB-RE/Hygro] Vector | | NanoLuc NF-κB reporter gene | Promega, Cat # N111A |
| 5 | pCMV3-His-eIF3i | His-eIF3i | N-His tagged human eIF3i | Sino Biological, Cat # HG16651-NH |
| 6 | pcDNA3.1-HA-Ubiquitin | | | Addgene, Cat # 18712 |
| 7 | pET28a-His-CULT | | His tagged human CULT for bacterial expression | Made in house |

| No. | Primer name | Details | Sequence |
|---|---|---|---|
| 1 | ZL_hRT001F | qRT-PCR primer for human IRF4 | GAACGAGGAGAAGAGCATCTTCC |
| 2 | ZL_hRT001R | qRT-PCR primer for human IRF4 | CGATGCCTTCTCGGAACTTTCC |
| 3 | ZL_hRT002F | qRT-PCR primer for human NF-κB | GCAGCACTACTTCTTGACCACC |
| 4 | ZL_hRT002R | qRT-PCR primer for human NF-κB | TCTGCTCCTGAGCATTGACGTC |
| 5 | ZL_hRT003F | qRT-PCR primer for human TNF-α | CTCTTCTGCCTGCTGCACTTTG |
| 6 | ZL_hRT003R | qRT-PCR primer for human TNF-α | ATGGGCTACAGGCTTGTCACTC |
| 7 | ZL_hRT022F | qRT-PCR primer for human eIF3i | CTGTGGCCAAGGACCCTATC |
| 8 | ZL_hRT022R | qRT-PCR primer for human eIF3i | CCAGTGAGGACATGCTTGGT |
| 9 | ZL_hRT025F | qRT-PCR primer for human GAPDH | GTCTCCTCTGACTTCAACAGCG |
| 10 | ZL_hRT025R | qRT-PCR primer for human GAPDH | ACCACCCTGTTGCTGTAGCCAA |
| 11 | ZL_mRT004F | qRT-PCR primer for mouse NF-κB | GCTGCCAAAGAAGGACACGACA |
| 12 | ZL_mRT004R | qRT-PCR primer for mouse NF-κB | GGCAGGCTATTGCTCATCACAG |
| 13 | ZL_mRT005F | qRT-PCR primer for mouse TNF-α | GGTGCCTATGTCTCAGCCTCTT |
| 14 | ZL_mRT005R | qRT-PCR primer for mouse TNF-α | GCCATAGAACTGATGAGAGGGAG |
| 15 | ZL_mRT006F | qRT-PCR primer for mouse GAPDH | CATCACTGCCACCCAGAAGACTG |
| 16 | ZL_mRT006R | qRT-PCR primer for mouse GAPDH | ATGCCAGTGAGCTTCCCGTTCAG |
| 17 | ZL_mRT007F | qRT-PCR primer for mouse IRF4 | GAACGAGGAGAAGAGCGTCTTC |
| 18 | ZL_mRT007R | qRT-PCR primer for mouse IRF4 | GTAGGAGGATCTGGCTTGTCGA |
| 19 | ZL_mRT024F | qRT-PCR primer for mouse eIF3i | AACAGAGCGTCCTGTCAACTCG |
| 20 | ZL_mRT024R | qRT-PCR primer for mouse eIF3i | CTTGCCAATCCTGGTGGAGGTT |
| 21 | CULT-F | pET28a-CULT cloning | GGAATTCCATATGATGAACAAGTGTACGAGCCTGT |
| 22 | CULT-R | pET28a-CULT cloning | GAGCTCGAATTCTTACAGGCAC |

#### Overexpression and purification of CULT.

pET28a-CULT was constructed by extracting the region encoding the CULT domain from a E. coli codon-optimized gene encoding human cereblon (synthesized by Atum Bio) using Primers 1 (CULT-F, GGAATTCCATATGATGAACAAGTGTACGAGCCTGT) and 2 (CULT-R, GAGCTCGAATTCTTACAGGCAC). The fragment was then introduced into pET28a using restriction enzyme cloning with NdeI and EcoRI. BL21 (DE3) cells transformed with pET28a-CULT were used to inoculate overnight cultures of LB + 50 µg/mL kanamycin, which were incubated at 30 °C with shaking at 200 rpm for approximately 16 h. For each large-scale overexpression, kanamycin was added to 750 mL autoclaved LB to a final concentration of 50 µg/mL and the culture was inoculated with overnight culture diluted 1:100. The overexpression cultures were incubated at 30 °C with shaking at 200 rpm until the OD600 was approximately

0.4, at which point isopropyl isopropyl β-D-1-thiogalactoside (IPTG) was added to a final concentration of 0.1 mM and the temperature was reduced to 25 °C. The cultures were incubated for 4-6 h prior to collecting the cells by centrifugation, flash freezing with liquid nitrogen, and storing at –80 °C.

Up to 6 pellets, each from a 750 mL overexpression culture, were purified simultaneously. To purify protein, cell pellets were thawed on ice or in cool water, then resuspended in 5 mL of 25 mM imidazole, 1× protease inhibitor, 1% Triton-X 100/PBS per pellet. Up to 3 pellets were combined and sonicated (30 s on, 10 s off, 5 min total, 25% amplitude) on ice together. Lysates were clarified by centrifugation (15,000 × g, 4 °C, 10 min) and syringe filtration (0.45 µm). The His-tagged protein was crudely purified on a Ni-bound 1 mL HiTrap Chelating HP column according to the manufacturer's instructions. The column was equilibrated and washed with 25 mM imidazole/PBS before the proteins were eluted with a gradient to 500 mM imidazole/PBS. Protein-containing fractions, as determined by A280, were taken directly to further purification on a HiLoad 16/600 Superdex 200 pg column, pre-equilibrated and run with PBS. Protein-containing fractions not eluting with the dead volume were collected and concentrated to approximately 100 µM with a 10 kDa MWCO spin concentrator, with protein concentration was determined by measuring the A280 (ε = 27,960 $M^{-1}$ $cm^{-1}$ with reduced cysteines). The protein was aliquoted, flash frozen with liquid nitrogen, and stored at –80 ˚C.

**Structural proteomics mass spectrometry sample preparation.**

For each analysis, 20 µM CULT and 22 µM of the indicated compound in 0.1% Triton-X 100 in PBS was prepared (34 µL, 10 µg of CULT) with the compounds added from 4 mM stock in DMSO. The sample was incubated for 30 min at 25℃, followed by photo-irradiation for 30 s at 4℃ via Dymax. A stock solution of 10% RapiGest/PBS was added to the photo-irradiated sample to a final concentration of 1 % (3.8 µL). The denatured proteins were tagged with an isotopically coded, cleavable biotin picolyl-azide probe[4] via CuAAC by addition of a pre-mixed cocktail (final concentrations: 100 µM cleavable biotin picolyl-azide probe, 250 µM THPTA, 250 µM $CuSO_4$, 2.5 mM freshly dissolved sodium ascorbate) and incubated for 1.5 h at 25℃ with inversion. Protein was precipitated by the addition of cold acetone (160 µL) and incubated for 30 min at −80℃. The precipitated protein was pelleted by centrifugation (4℃, 10 min, 21,300 x g). The supernatant was discarded, and the protein pellet was air dried for 10 min at 25℃. The protein was resuspended in 100 mM TEAB (50 µL, pH=8.5) and 100 mM DTT/PBS (1.5 µL, final concentration 3 mM) was added before incubating for 30 min at 25℃ with inversion. 500 mM iodoacetamide/PBS (1.0 µL, final concentration 10 mM) was then added to the sample and incubated for 30 min at 25℃ in the dark with inversion. The protein was then digested by adding sequence-grade chymotrypsin (0.5 µg) in 1 mM HCl and incubating for 15–18 h at 25℃ with inversion. The digested protein was acidified by formic acid (1.0 µL) and incubated for 30 min at 25℃ with inversion to cleave the cleavable biotin picolyl-azide probe. The sample was desalted using a C18 Ziptip according to the manufacture's protocol with elution in 0.1% formic acid/70% acetonitrile/water and the desalted peptides were dried on a Speed-Vac. The dried sample was re-suspended in 20 µL 0.1% formic acid/water for analysis.

**Structural proteomics mass spectrometry procedures.**

The sample was loaded onto a microcapillary trapping column (C18 Reprosil resin, Dr. Maisch, 5 µm particle size, 100 Å pore size, 30 mm length, 100 µm internal diameter) and then separated on an analytical column (50 cm µPAC™ column, PharmaFluidics) at 200 nL/min. The column temperature was maintained at 40℃. Peptides were eluted with a water/acetonitrile gradient (buffer A = 0.1% formic acid/water, buffer B = 0.1% formic acid/acetonitrile; flow rate 200 nL/min; gradient: hold at 2% B for 5 min, increase to 6% B over 1 min, increase to 35% B over 29 min, increase to 95% B over 5 min, hold at 95% B for 20 min). Electrospray ionization was enabled through applying a voltage of 1.8 kV using a home-made electrode junction at the end of the microcapillary column and sprayed from stainless-steel needle (PepSep, Denmark). The Lumos Orbitrap was operated in data-dependent mode and survey scans of peptide precursors were performed at 120K FWHM resolution over a *m/z* range of 400-1800, an AGC setting of 1,000,000, and the maximum ion accumulation time of 50 ms. Tandem MS was performed on the 10 most abundant precursors with an AGC setting of 10,000 and fragmentation energy of 35% for CID, at 15K FWHM resolution, an AGC setting of 10,000 and fragmentation energy of 37% for HCD. An isolation window of 3 *m/z* used to ensure capture of isotopically coded species. With a mass tolerance of 10 ppm, selected precursors were excluded from further fragmentation for 60 s.

**Structural proteomics data analysis procedures**.

The raw data were analyzed using Proteome Discoverer 2.4.1.15. Assignment of MS/MS spectra was performed using the Sequest HT algorithm by searching the data against a protein sequence database of CULT and common contaminant proteins. Search parameters included: mass tolerance of 10 ppm for the precursor, 0.02 Da for HCD fragment ions, 0.6 Da for CID fragment ions, semi-specific digestion, 2 missed cleavages, static modification of carbamidomethylation on cysteine residues (+57.0214 Da), a dynamic oxidation on methionine residues (+15.9949 Da), a dynamic deamidation on asparagine and glutamine residues (+0.9847 Da) and a dynamic modification of the compounds on any amino acid residues (photo-lenalidomide: +588.2355 Da, analog **1**: +602.2512 Da). Peptide spectral matches (PSMs) were validated with the Target Decoy PSM Validator. Spectra assigned as photo-lenalidomide/analog **1**-conjugated peptides were manually validated by evaluating the isotopic coding embedded in the MS1 precursor.

**Structural modeling details.**

Docking study was performed with MOE version 2020.09. Pomalidomide in the crystal structure of DDB1-CRBN-pomalidomide-IKZF1 ZF2 complex (PDB 6H0F)[11] was adapted to the indicated compounds and the adapted complex was subjected to energy minimization with Amber10:EHT force field followed by preparation with Protonate 3D. Ligand conformations were generated with the bond rotation method and the docking was performed with the Triangle Matcher placement method and scored with the London dG scoring function. The lowest energy conformations were rendered and 30 of them were submitted to a refinement step with the Induced Fit method and rescored with the GBVI/WSA dG scoring function. A conformation of each complex with the lowest S score given by the GBVI/WSA dG scoring function was analyzed for the ligand interactions.

**Immunoprecipitation assays in live cells.**

For immunoprecipitation of proteins with the Flag tag, cell lysates with equal amounts of protein were diluted with the lysis buffer and incubated with Anti-Flag M2 magnetic beads (pre-washed three times with TBS) for 2 h at 4 ˚C with rotation. The beads were washed three times with TBS buffer (50 mM Tris HCl pH 7.4, 150 mM NaCl). The enriched proteins were eluted with SDS sample buffer for SDS-PAGE or 50 µL 3X Flag peptide solution (final concentration 150 ng/µl), according to the manufacturer's instructions.

For immunoprecipitation of proteins with the EPEA tag, cell lysates with equal amounts of protein were diluted with PBS and incubated with C-tag affinity matrix (pre-washed three times with PBS) for 2 h at 25 ˚C. The beads were washed three times with PBS buffer. The enriched proteins were eluted with SDS sample buffer for SDS-PAGE or 50 µL neutral pH-based elution buffer (20 mM Tris, 2 M $MgCl_2$, pH 7.0), according to the manufacturer's instructions.

For endogenous protein immunoprecipitation using protein A/G beads, cell lysates with equal amounts of protein were diluted with PBS and incubated with protein A/G beads (pre-washed three times with binding buffer, 50 mM Tris, 150 mM NaCl, 0.2% Triton, pH=7.5) for 2 h at 4 ˚C along with the protein-specific antibody at the vendor-suggested dilution. The beads were washed three times with binding buffer (50 mM Tris, 150 mM NaCl, 0.2% Triton, pH=7.5). The enriched proteins were eluted with acidic elution buffer (100 mM glycine, 0.1% Triton, pH=2.8) before neutralizing with 1M Tris (pH=8), according to the manufacturer's instructions.

**Cycloheximide (CHX) treatment.**

HEK293T cells were treated with DMSO or indicated drug 1 h before the addition of 50 µg/mL CHX for up to 24 h. At the indicated time points, cells were harvested and washed with PBS twice before lysis with 1X cell lysis buffer with 1X PI. The samples were then subjected to Western blot analysis.

**Photo-affinity labeling experiment and in-gel fluorescent imaging.**

Cells were plated on a 6-well plate and treated with the indicated drugs for 1 h at 37 ˚C before photo-irradiation for 120 s at 4 ˚C via Dymax. After immunoprecipitation, the eluents were then tagged with AlexaFluor® 488 azide via CuAAC by addition of a pre-mixed cocktail (final concentrations: 25 µM AlexaFluor® 488 azide, 100 µM THPTA, 1 mM $CuSO_4$, 2 mM freshly dissolved sodium ascorbate) and incubated for 1 h at 25 ˚C with inversion. The reactions were quenched by the addition of pre-chilled acetone and incubated for 20 min at −80 ˚C. The protein precipitates were isolated by centrifugation (10 min, 4 ˚C, 21,300 x g) and the supernatant was discarded. The protein pellet was dried for 10 min at 25 ˚C and then re-suspended in 50 µL 1% RapiGest/PBS by sonication (2 s on, 5 s off, 10% amplitude, 10 s total) before visualization by SDS-PAGE.

**Cell viability assay (MTT).**

Cells were treated with the indicated compound in triplicate on a 96-well plate at $1 \times 10^6$ cells/mL in 100 µL per well for 5 d at 37 ˚C. 3-(4,5-Dimethylthiazol-2-yl)-2,5-diphenyltetrazolium bromide (MTT) solution (10 µL, freshly dissolved in PBS at 4 mg/mL) was added to each well and the cells were incubated for 4 h at 37 °C. After incubation, MTT solubilization solution (100 µL, 10% SDS with 0.04 N HCl) was added to each well and incubated for 16 h at 37 °C. The absorbance was measured at a wavelength of 570 nm on a multimode microplate reader FilterMax. Background absorbance signal from media-only blank wells was subtracted and the absorbance signal was normalized to that of untreated cells.

**Flow cytometry (FACS).**

Cells were fixed with 4% formaldehyde for 10 min at 37 °C, and permeabilized by 90% MeOH at 4 °C for 30 min. The samples were then washed three times with PBS, blocked in 1% BSA for 15 min at 25 °C, and incubated with FITC- or PE-conjugated antibodies in 1% BSA for 30 min at 4 °C covered with foil. The samples were washed three times with PBS and analyzed. Cells were gated with SSC-A/FSC-A for live population and FSC-H/FSC-A for singlets, and 100,000 cells were analyzed from each sample. Relative number of FITC- or PE-expressing cells for different treatment conditions was calculated by BD FACSDiva Software and analyzed by Flowjo.

**RNA isolation and qRT-PCR experiment.**

Cells were treated as indicated before RNA isolation. Qiagen RNeasy-Plus Mini kits were used to purify RNA from cells, according to the instructions from the manufacturer. Isolated RNA then underwent two-step real-time RT-PCR by using PrimeScript RT reagent kit for reverse transcription and QuantiTect SYBR Green PCR kit.

**Luciferase reporter assay.**

HEK293T cells transfected with NLucP/NF-κB vector were incubated for 6 h in medium without serum prior to use. The cells were treated with either DMSO or the indicated compound and induced with 10 μM TNF-α for 5 h at 37 °C in the incubator before luciferase activity detection. Luciferase reporter assays were performed using the Luciferase Assay System (Promega, Cat # N1111) according to manufacturer's protocols.

**In vivo thermal shift assay.**

Cells were harvested and suspended in PBS supplemented with 1X PI at 2 × $10^7$ cells/mL. The cells were treated with either DMSO or the indicated compound for 1 h at 37 °C. Each fraction (90 μL) was incubated at the indicated temperature for 5 min, followed by a 3 min incubation on ice.[12] Cells were then lysed by addition of 10 μL 10X cell lysis buffer with 50X PI to a final concentration of 1X before the samples were centrifuged in a mini centrifuge at 21,000 g for 10 min at 4 °C. Supernatants were separated from the pellet and collected for Western blot analysis.

**Photo-affinity labeling and enrichment of lenalidomide targets in living cells and on-bead digestion.**

Cells were plated on a 6-well plate and treated with the indicated compound for 1 h at 37 ˚C before photo-irradiation for 120 s at 4 ˚C via Dymax. After washing with PBS, cells were collected and lysed using 1X cell lysis buffer with 1X PI. The cleared lysates were then tagged with a cleavable biotin azide probe via CuAAC by addition of a pre-mixed cocktail (final concentrations: 100 μM cleavable biotin azide probe, 100 μM THPTA, 1 mM $CuSO_4$, 2 mM freshly dissolved sodium ascorbate) and incubated for 3 h at 25 ˚C with inversion. The reactions were quenched by the addition of pre-chilled acetone and incubated for 20 min at −20 ˚C to precipitate the proteins. Followed by centrifugation (10 min, 4 ˚C, 21,300 x g), the protein was pelleted with supernatant discarded, and dried for 10 min at 25 ˚C to full dryness. The protein pellet was re-suspended in 1% RapiGest/PBS by sonication (2 s on, 5 s off, 10% amplitude, 10 s total). The protein solutions were then diluted to the same concentrations after BCA assay, and 50 μL of the solutions were taken and saved as input samples for Western blot analysis. Streptavidin beads slurry (80 μL for each sample) was pre-washed three times with 400 μL PBS in Bio-Spin columns before the protein solutions were added, followed by incubation for 16 h at 25 ˚C with inversion. The beads were then washed with 1 mL freshly dissolved 8 M urea for three times, 1 mL 1% RapiGest/PBS for three times, and 1 mL PBS for three times. After the wash, 200 μL PBS was added and 25 μL of the bead slurry was taken out from the columns and saved as biotin enrichment samples for Western blot analysis. Rest of the bead slurry were subjected to on-bead digestion for mass spectrometry analysis.

In order to perform on-bead digestion, 25 μL of 10 mM DTT/PBS was added to each sample for 30 min at 25 ˚C with inversion. The solution was discarded using the vacuum manifold and another 200 μL of PBS was added together with 10 μL of 500 mM iodoacetamide/PBS to each Bio-Spin column to alkylate reduced cysteines for 30 min at 25 ˚C with inversion. The solutions were removed using the vacuum manifold and the beads were washed with 1 mL PBS and 1 mL 50 mM HEPES (pH=8.5) for one time before 100 μL 0.5 M guanidinium chloride (GdnHCl) in 50 mM HEPES (pH=8.5) was added into each column. Proteins on the beads were then digested by adding sequence-grade trypsin (3 μg, 6 μL of a 0.5 mg/mL stock solution) for 16 h at 37 ˚C with inversion. The flow-through of the Bio-Spin columns after centrifugation containing the digested peptides were collected. The beads were washed with 200 μL 0.5 M GdnHCl in 50 mM HEPES once and 200 μL 50% acetonitrile/ water once with the flow-through all combined with the digested peptides. The eluted peptides were then dried on the Speed-Vac to dryness and subjected to mass spectrometry sample preparation.

**Mass spectrometry sample preparation.**

Digested peptides were resuspended in 50 µL of 50 mM TEAB buffer (pH=8.5), treated with 5 µL of the TMT 16-plex reagents (10 µg/µL) and then incubated for 1 h at 25 ˚C. The TMT labeling was quenched with 2 µL of 5% hydroxylamine solution for 15 min at 25 ˚C. The TMT-labeled peptides were then combined and dried on the Speed-Vac. The peptides were resuspended in 100 µL 1% formic acid/water and desalted with C18 Tips (Zip-Tip) according to the manufacturer's instructions and stored at -20 ˚C until analysis.

**ERLIC separation and mass spectrometry analysis.**

The sample was separated using the Pierce™ High pH Reversed-Phase Peptide Fractionation Kit (Thermo Fischer) according to manufacturer's instructions. After separation, each fraction was submitted for a single LC-MS/MS experiment that was performed on a Lumos Tribrid Orbitrap equipped with an UltiMate 3000 RSLCnano System. Peptides were separated onto a 100 µm inner diameter microcapillary trapping column packed first with approximately 3 cm of C18 Reprosil resin (5 µm, 100 Å, Dr. Maisch GmbH, Germany) followed by analytical column 50 cm microcapillary based PharmaFluidics (Belgium). Separation was achieved through applying a gradient from 5–27% acetonitrile in 0.1% formic acid in water over 90 min at 200 nl $min^{-1}$. Electrospray ionization was enabled through applying a voltage of 1.8 kV using a home-made electrode junction at the end of the microcapillary column and sprayed from stainless-steel needle (PepSep, Denmark). The Lumos Orbitrap was operated in data-dependent mode for the mass spectrometry methods. The mass spectrometry survey scan was performed in the Orbitrap in the range of 400–1,800 m/z at a resolution of $1.2 \times 10^5$, followed by the selection of the 10 most intense ions (TOP10) for CID-MS2 fragmentation in the Ion trap using a precursor isolation width window of 2 m/z, an AGC setting of 10,000, and a maximum ion accumulation of 200 ms. Singly charged ion species were not subjected to CID fragmentation. Normalized collision energy was set to 35 V and an activation time of 10 ms. Ions in a 10-ppm m/z window around ions selected for MS2 were excluded from further selection for fragmentation for 60 s. The same TOP10 ions were subjected to HCD MS2 event in Orbitrap part of the instrument. The fragment ion isolation width was set to 0.8 Da, with 0.3 Da offset, AGC was set to 50,000, the maximum ion time was 100 ms, normalized collision energy was set to 34 V and an activation time of 1 ms for each HCD MS2 scan.

**Mass spectrometry analysis**.

The raw data were analyzed using Proteome Discoverer 2.4.1.15. Assignment of MS/MS spectra was performed using the Sequest HT algorithm by searching the data against a protein sequence database including all entries from the Human Uniprot database (Aug 19, 2016, 20,156 SwissProt) and a list of common contaminant proteins. Search parameters included: a 10-ppm precursor ion tolerance and 0.02 Da fragment ion tolerance for HCD or 0.6 Da fragment ion tolerance for CID, full tryptic protease specificity with up to two missed cleavages, a static modification of 16-plex TMTpro tags on peptide N-termini and lysine residues (+304.2071 Da), a static modification of cysteine carbamidomethylation (+57.0214 Da), a dynamic modification of methionine oxidation (+15.9949 Da), a dynamic modification of deamidation on asparagine and glutamine residues (+0.9847 Da) and a dynamic modification of acetylation on peptide N-termini (+42.0106 Da). Peptide spectral matches (PSMs) were filtered for a false discovery rate (FDR) of 1% following a target-decoy analysis with Percolator. Reporter ions were quantified using a 0.02 m/z window centered on the theoretical m/z value and the intensity of the signal closest to the theoretical m/z value was recorded. Reporter ion intensities were exported in result file of Proteome Discoverer search engine as an excel tables. The total signal intensity across all peptides quantified was summed and normalized for each TMT channel. Quantified proteins were required to have at least three unique peptides and four separate PSMs. All TMT-based experiments were performed with four replicates.

### In vivo ubiquitylation assay.

In vivo ubiquitylation assay was performed according to the method of Jaffrey and co-workers.[13] HEK293T or HEK293T CRBN-KD cells[14] transiently transfected with HA-Ub were treated with either DMSO or the indicated compound for 1 h before 10 μM MG132 was added. Treated cells were incubated for 6 h at 37 ˚C and collected. Ubiquitylated proteins were immunoprecipitated by the indicated antibody according to the manufacturer's instructions. Samples were then eluted and subjected to Western blot analysis.

### Software.

Data was analyzed and visualized using Microsoft Excel (v16.22) and GraphPad Prism (v8.0.1), in addition to software listed by experiment. NMR data were analyzed using MestReNova (v14.0.1). DNA and protein sequences were analyzed using Geneious (v10.0.7). FACS data were analyzed by FlowJo (v10.6.1). Proteomics data were analyzed by Proteome Discoverer (v2.4.1.15). Images were made using ImageJ (NIH, v1.50i), ImageStudioLite (v5.2.5), and Adobe Illustrator (v22.1). Docking studies were performed with Molecular Operating Environment (MOE, v2020.09) and the obtained structures were analyzed with PyMOL (v2.3.1).

### Statistical analysis.

Statistical analyses (unpaired Student's t-tests) were performed using GraphPad Prism. Data were derived from at least three biological replicate experiments and presented as the mean ± s.d., *P< 0.0332, **P< 0.0021, ***P < 0.0002, ****P < 0.0001 and n.s., not significant.

### REFERENCES


1. Still, W. C.; Kahn, M.; Mitra, A., Rapid chromatographic technique for preparative separations with moderate resolution. *J Org Chem* **1978,** *43* (14), 2923-2925.
2. Li, Z.; Hao, P.; Li, L.; Tan, C. Y.; Cheng, X.; Chen, G. Y.; Sze, S. K.; Shen, H. M.; Yao, S. Q., Design and synthesis of minimalist terminal Alkyne-Containing diazirine photo-crosslinkers and their incorporation into kinase inhibitors for cell-and tissue-based proteome profiling. *Angew Chem Int Ed* **2013,** *125* (33), 8713-8718.
3. Gao, J.; Mfuh, A.; Amako, Y.; Woo, C. M., Small molecule interactome mapping by photoaffinity labeling reveals binding site hotspots for the NSAIDs. *J Am Chem Soc* **2018,** *140* (12), 4259-4268.
4. Miyamoto, D. K.; Flaxman, H. A.; Wu, H. Y.; Gao, J.; Woo, C. M., Discovery of a Celecoxib Binding Site on Prostaglandin E Synthase (PTGES) with a Cleavable Chelation-Assisted Biotin Probe. *ACS Chem Biol* **2019,** *14* (12), 2527-2532.
5. Pangborn, A. B.; Giardello, M. A.; Grubbs, R. H.; Rosen, R. K.; Timmers, F. J., Safe and convenient procedure for solvent purification. *Organometallics* **1996,** *15* (5), 1518-1520.
6. Muller, G. W.; Chen, R.; Huang, S.-Y.; Corral, L. G.; Wong, L. M.; Patterson, R. T.; Chen, Y.; Kaplan, G.; Stirling, D. I., Amino-substituted thalidomide analogs: potent inhibitors of TNF-α production. *Bioorganic Med Chemi Lett* **1999,** *9* (11), 1625-1630.
7. Kleiner, P.; Heydenreuter, W.; Stahl, M.; Korotkov, V. S.; Sieber, S. A., A whole proteome inventory of background photocrosslinker binding. *Angew Chem Int Ed* **2017,** *56* (5), 1396-1401.
8. Matyskiela, M. E.; Lu, G.; Ito, T.; Pagarigan, B.; Lu, C.-C.; Miller, K.; Fang, W.; Wang, N.-Y.; Nguyen, D.; Houston, J., A novel cereblon modulator recruits GSPT1 to the CRL4 CRBN ubiquitin ligase. *Nature* **2016,** *535* (7611), 252-257.
9. Sebej, P.; Lim, B. H.; Park, B. S.; Givens, R. S.; Klán, P., The power of solvent in altering the course of photorearrangements. *Org Lett* **2011,** *13* (4), 644-647.
10. Man, H.-W.; Muller, G. W.; Ruchelman, A. L.; Khalil, E. M.; Chen, R. S.-C.; Zhang, W. Arylmethoxy isoindoline derivatives and compositions comprising and methods of using the same. US2011/0196150A1, 2011.
11. Sievers, Q. L. P., G., Bunker, R. D.; Renneville, A.; Slabicki, M.; Liddicoat, B. J.; Abdulrahman, W.; Mikkelsen, T.; Ebert, B. L.; Thoma, N. H., Defining the human C2H2 zinc finger degrome targeted by thalidomide analogs through CRBN. *Science* **2018,** *362* (6414), eaat0572.
12. de Wispelaere, M.; Du, G.; Donovan, K. A.; Zhang, T.; Eleuteri, N. A.; Yuan, J. C.; Kalabathula, J.; Nowak, R. P.; Fischer, E. S.; Gray, N. S.; Yang, P. L., Small molecule degraders of the hepatitis C virus protease reduce susceptibility to resistance mutations. *Nat Commun* **2019,** *10* (1), 3468.
13. Xu, G.; Jiang, X.; Jaffrey, S. R., A mental retardation-linked nonsense mutation in cereblon is rescued by proteasome inhibition. *J Biol Chem* **2013,** *288* (41), 29573-85.
14. Nguyen, T. V.; Lee, J. E.; Sweredoski, M. J.; Yang, S. J.; Jeon, S. J.; Harrison, J. S.; Yim, J. H.; Lee, S. G.; Handa, H.; Kuhlman, B.; Jeong, J. S.; Reitsma, J. M.; Park, C. S.; Hess, S.; Deshaies, R. J., Glutamine Triggers Acetylation-Dependent Degradation of Glutamine Synthetase via the Thalidomide Receptor Cereblon. *Mol Cell* **2016,** *61* (6), 809-20.

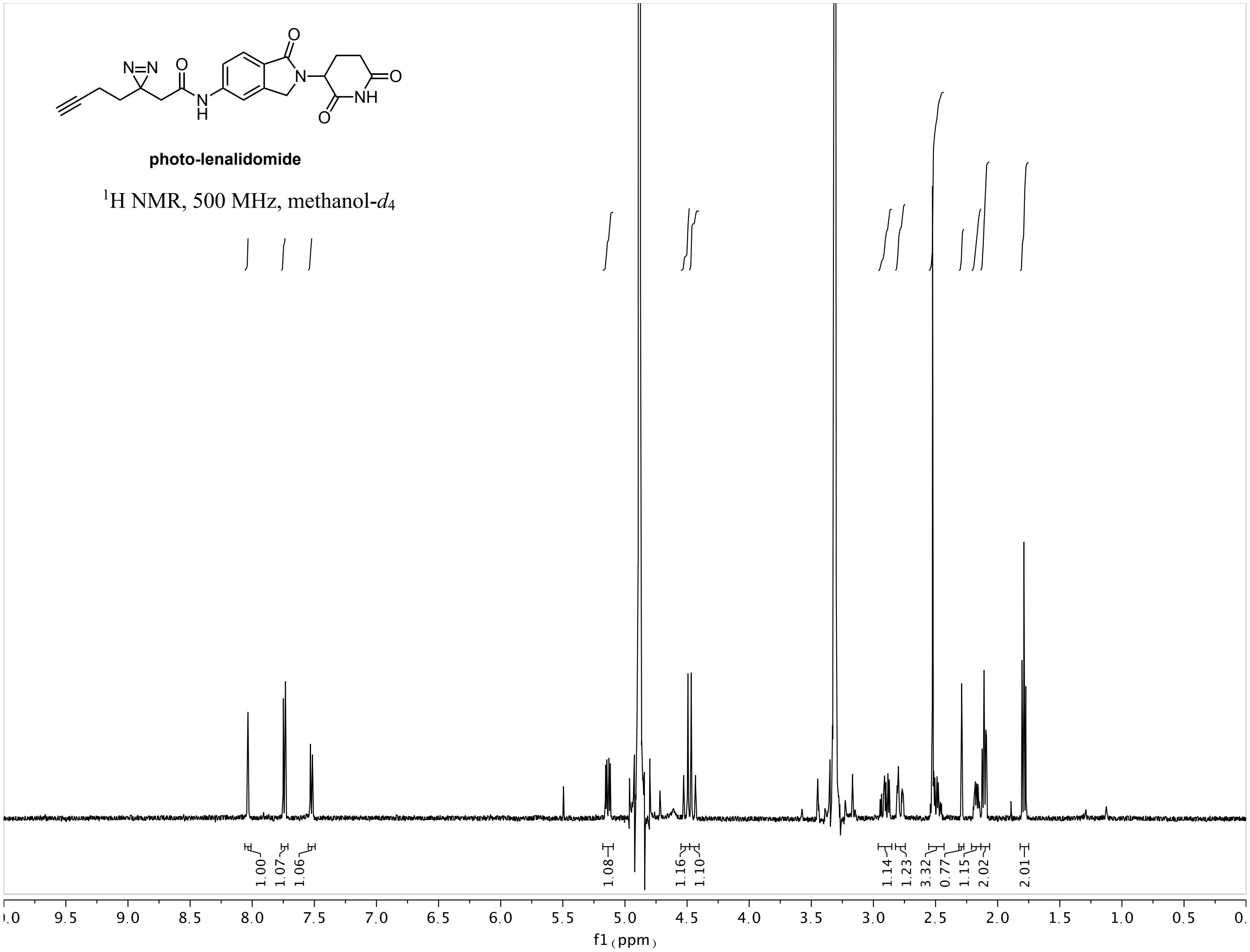

photo-lenalidomide
$^1$H NMR, 500 MHz, methanol-$d_4$
1.00
1.07
1.06
1.08
1.16
1.10
1.14
1.23
3.32
0.77
1.15
2.02
2.01
f1 (ppm)

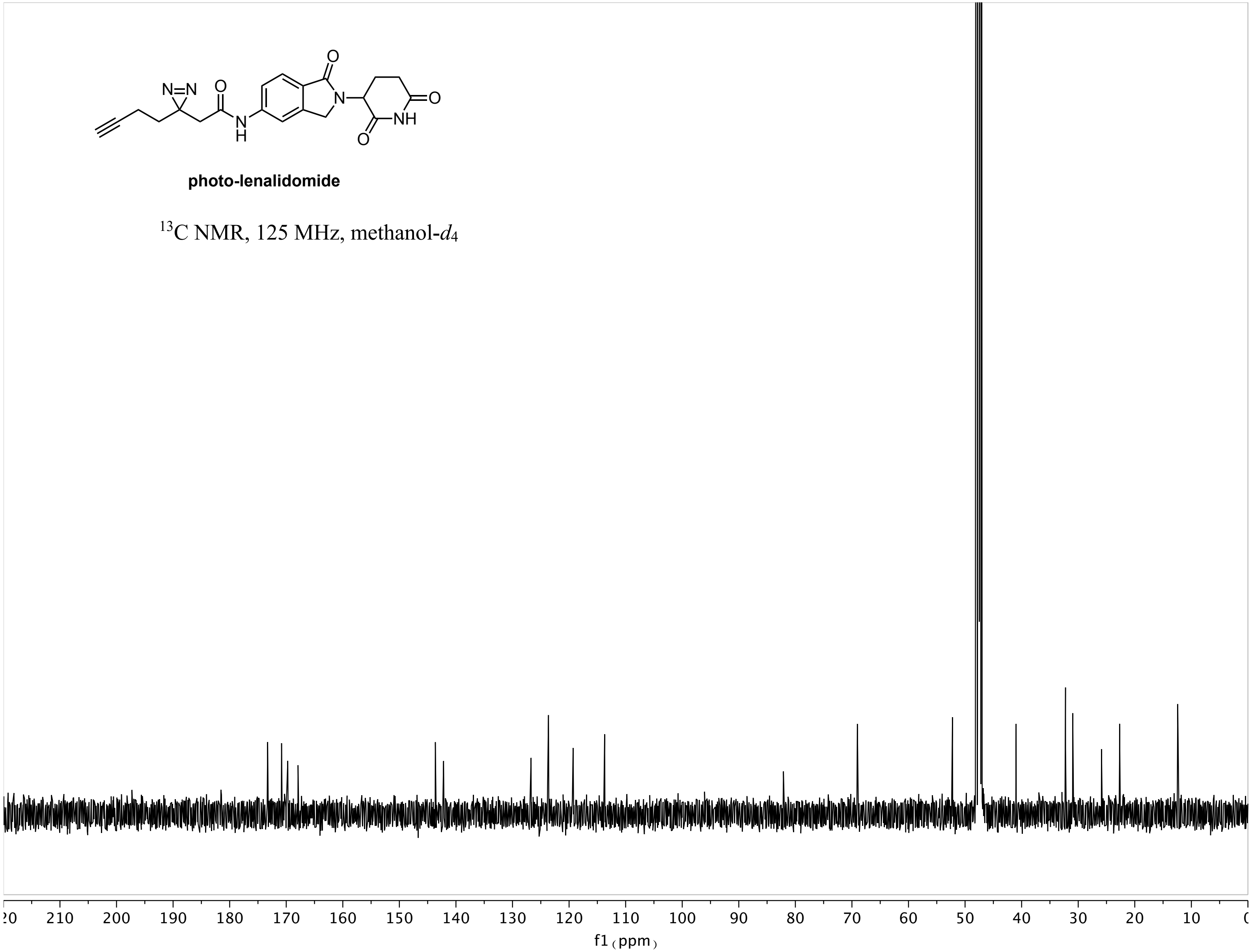
N=N
O
O
N
H
N
NH
O
O
photo-lenalidomide
$^{13}$C NMR, 125 MHz, methanol-$d_4$
210 200 190 180 170 160 150 140 130 120 110 100 90 80 70 60 50 40 30 20 10
f1 (ppm)

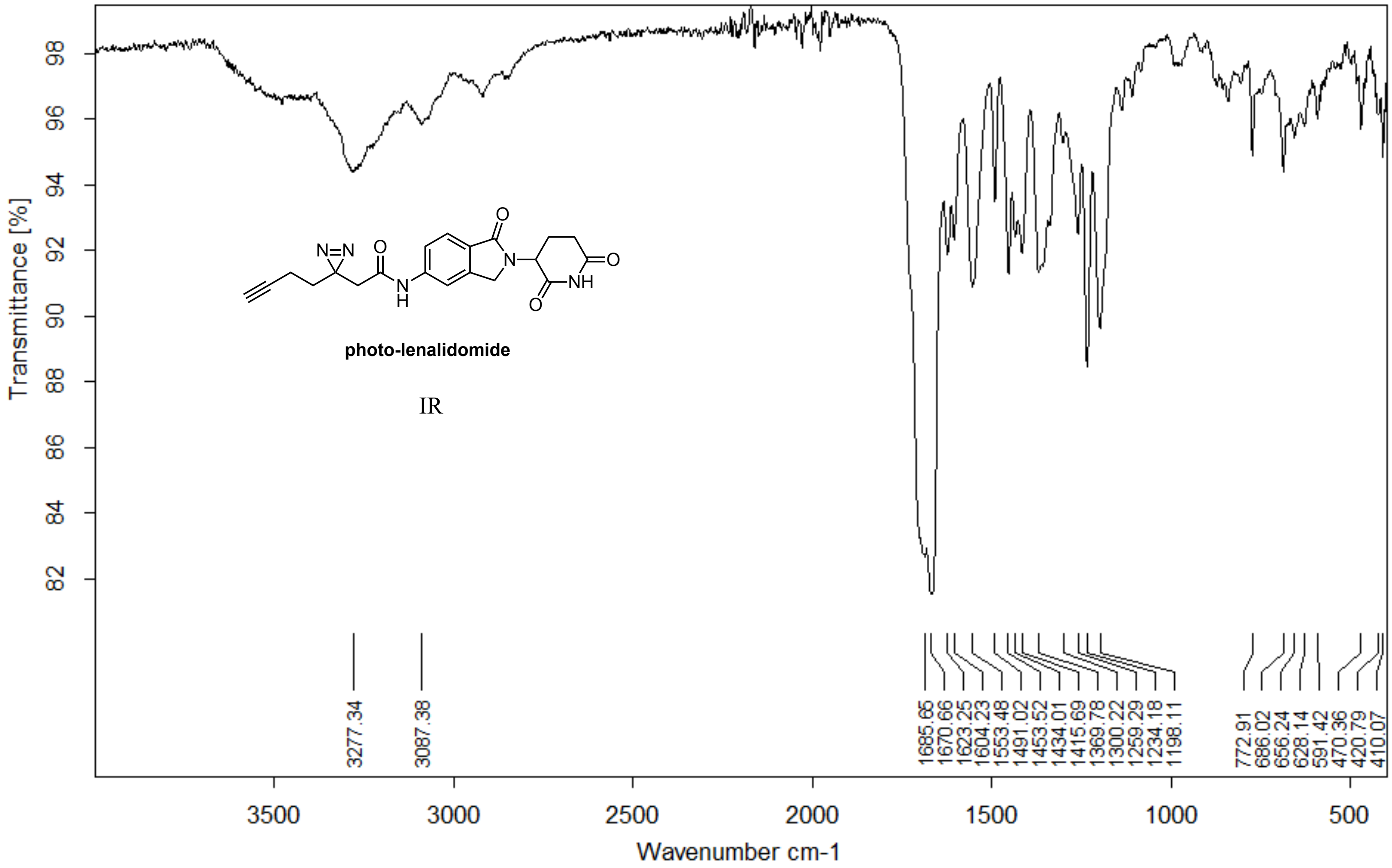


| | | |
|---|---|---|
| Z:\Alpha FTIR Data\Woo\Yuka\YA338.0 | YA338 | Diamond ATR |

07/06/2018

**Page 1/2**

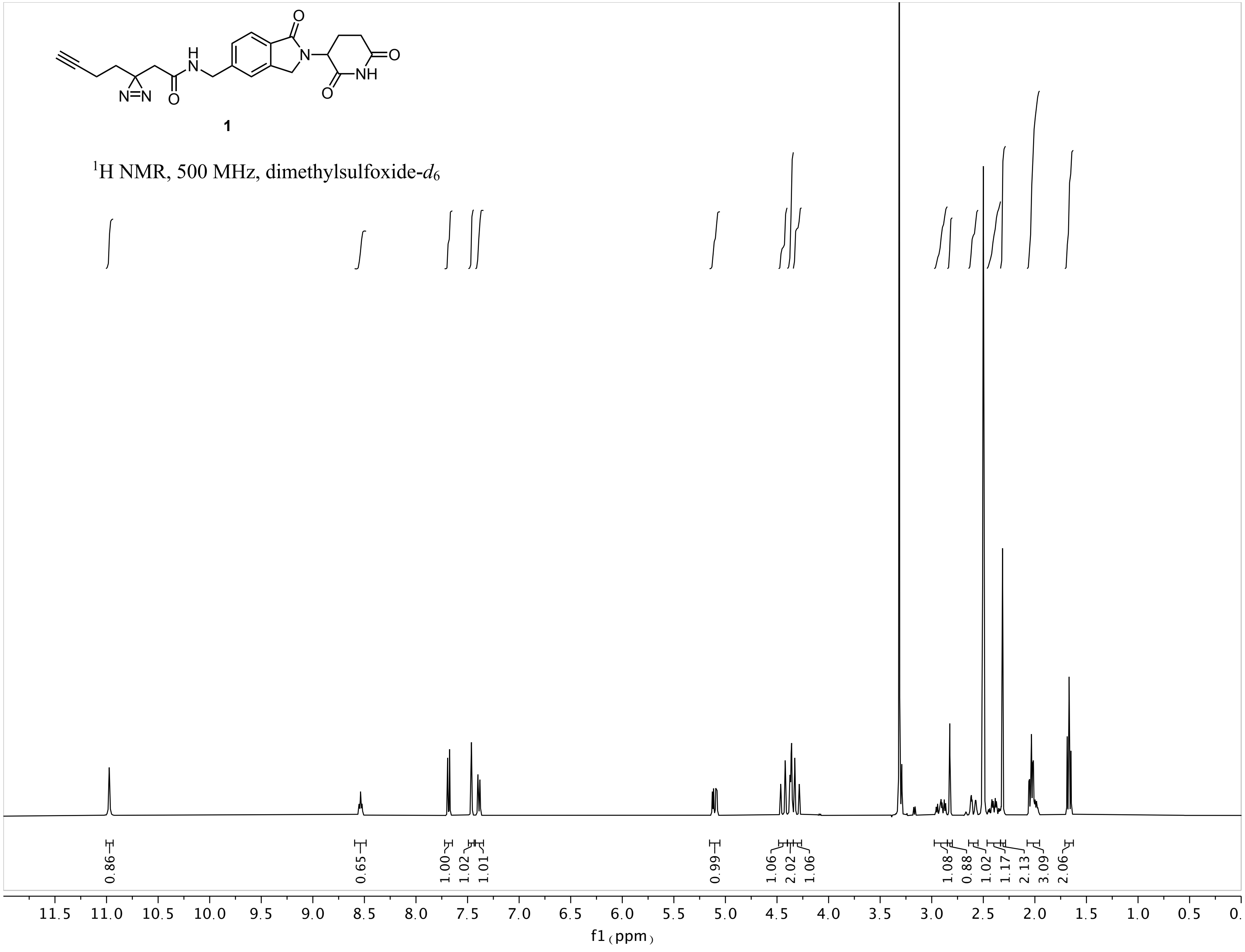

1
$^1$H NMR, 500 MHz, dimethylsulfoxide-$d_6$
0.86
0.65
1.00
1.02
1.01
0.99
1.06
2.02
1.06
1.08
0.88
1.02
1.17
2.13
3.09
2.06
11.5 11.0 10.5 10.0 9.5 9.0 8.5 8.0 7.5 7.0 6.5 6.0 5.5 5.0 4.5 4.0 3.5 3.0 2.5 2.0 1.5 1.0 0.5 0.
f1 (ppm)

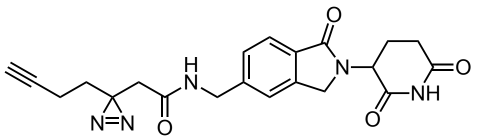

**1**

$^{13}$C NMR, 125 MHz, dimethylsulfoxide-$d_6$

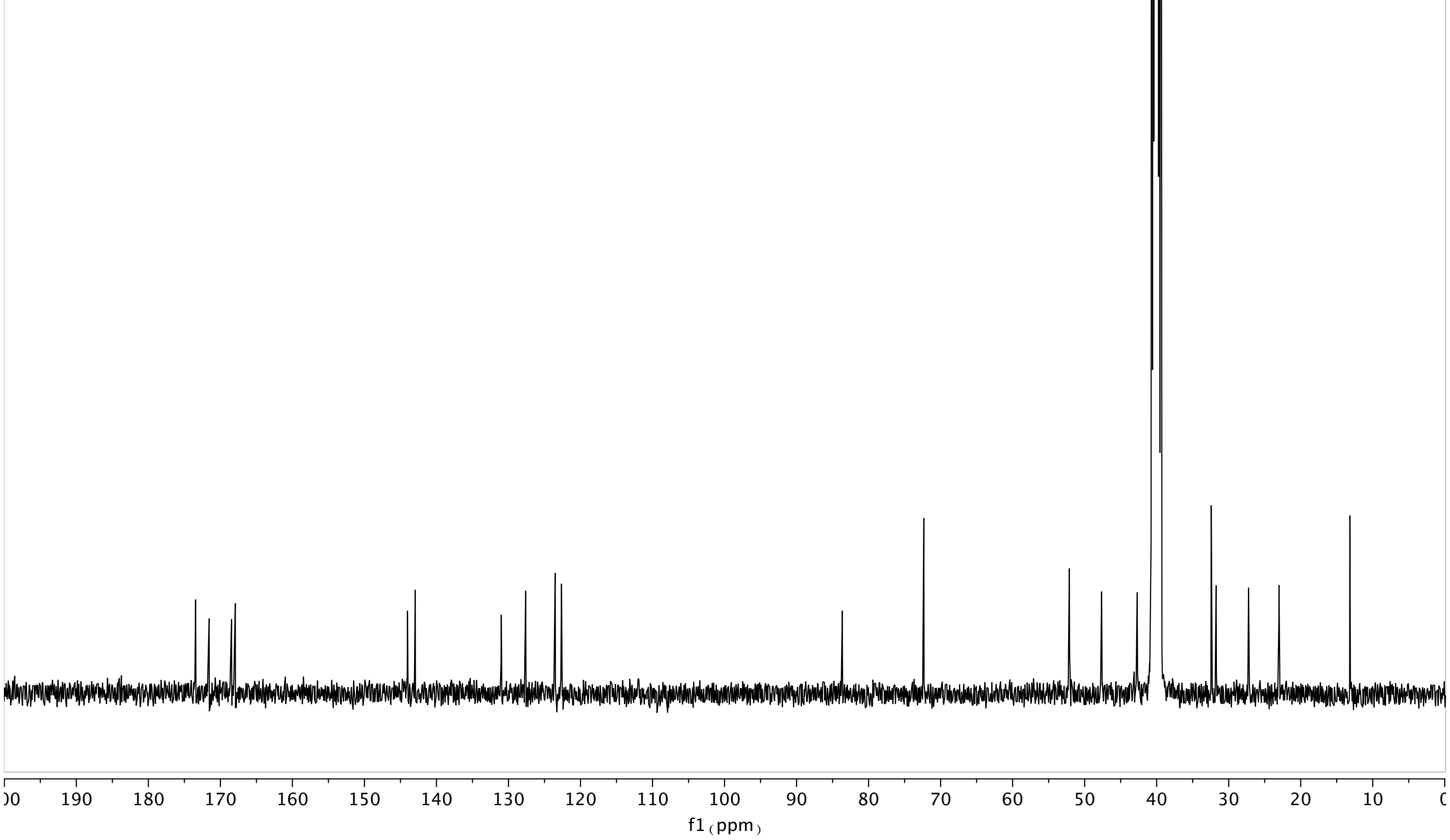

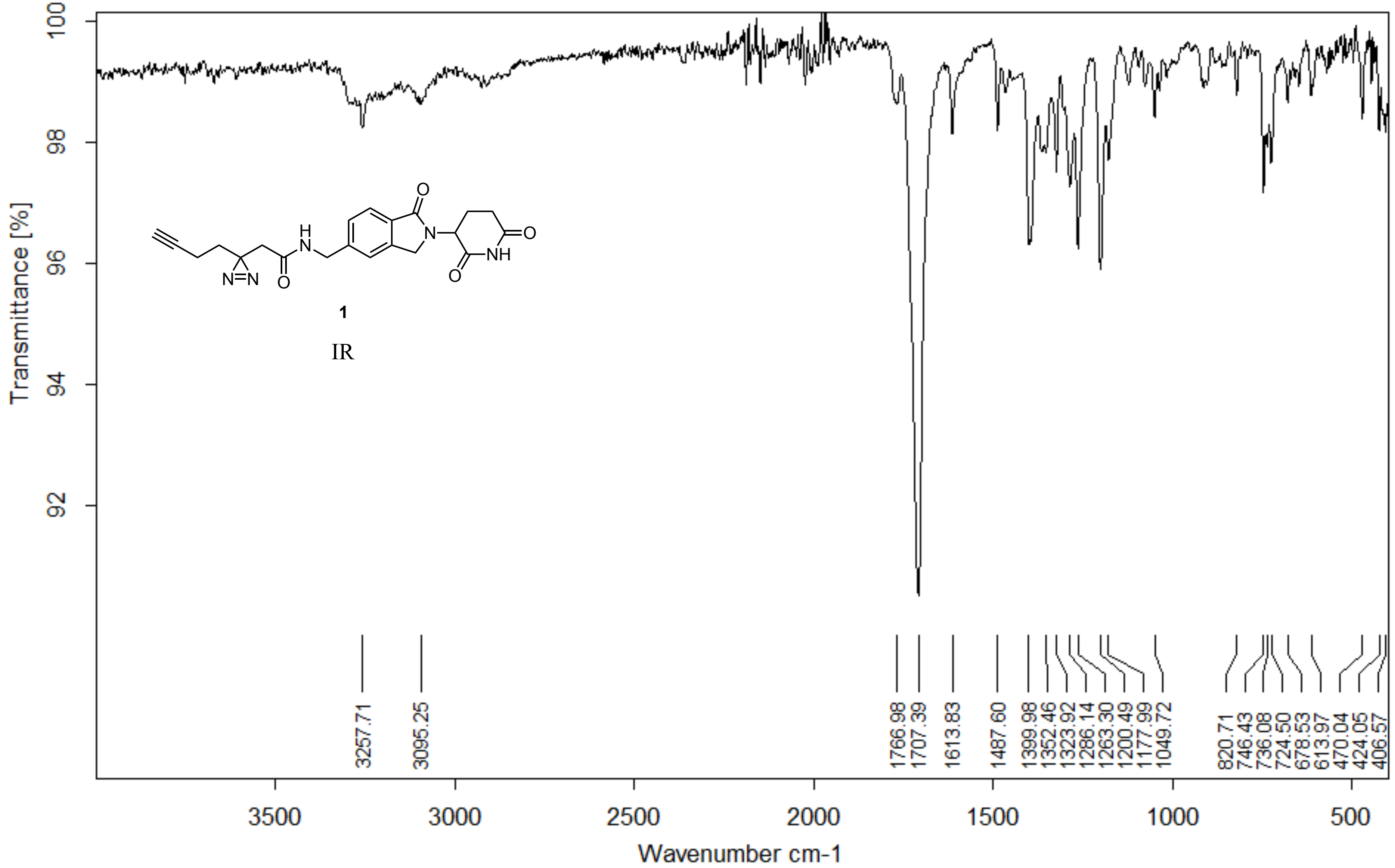


| | | | |
|---|---|---|---|
| Z:\Alpha FTIR Data\Woo\Yuka\YA322.0 | YA322 | Diamond ATR | 28/05/2018 |

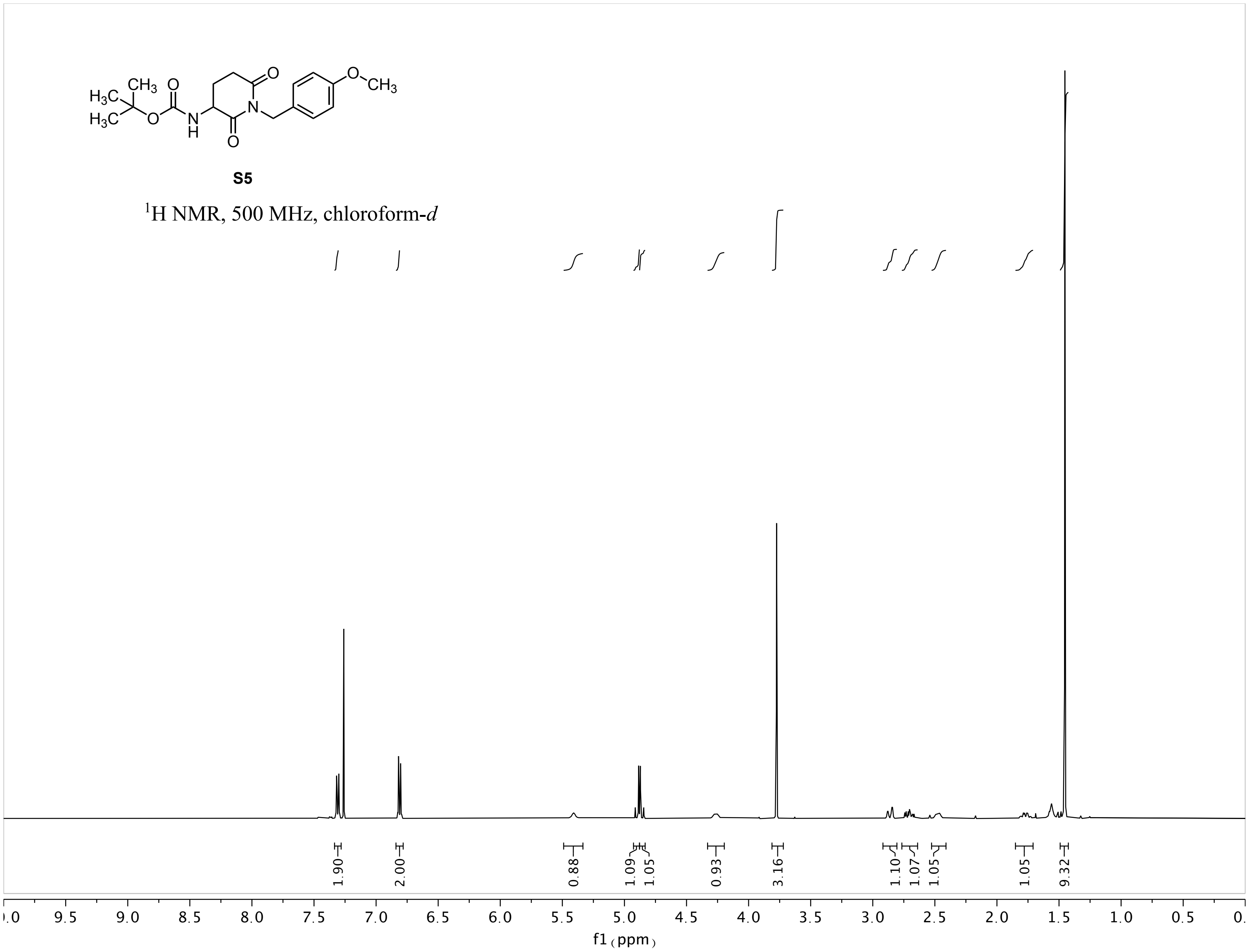
S5
$^{1}$H NMR, 500 MHz, chloroform-*d*
1.90
2.00
0.88
1.09
1.05
0.93
3.16
1.10
1.07
1.05
1.05
9.32
f1 (ppm)

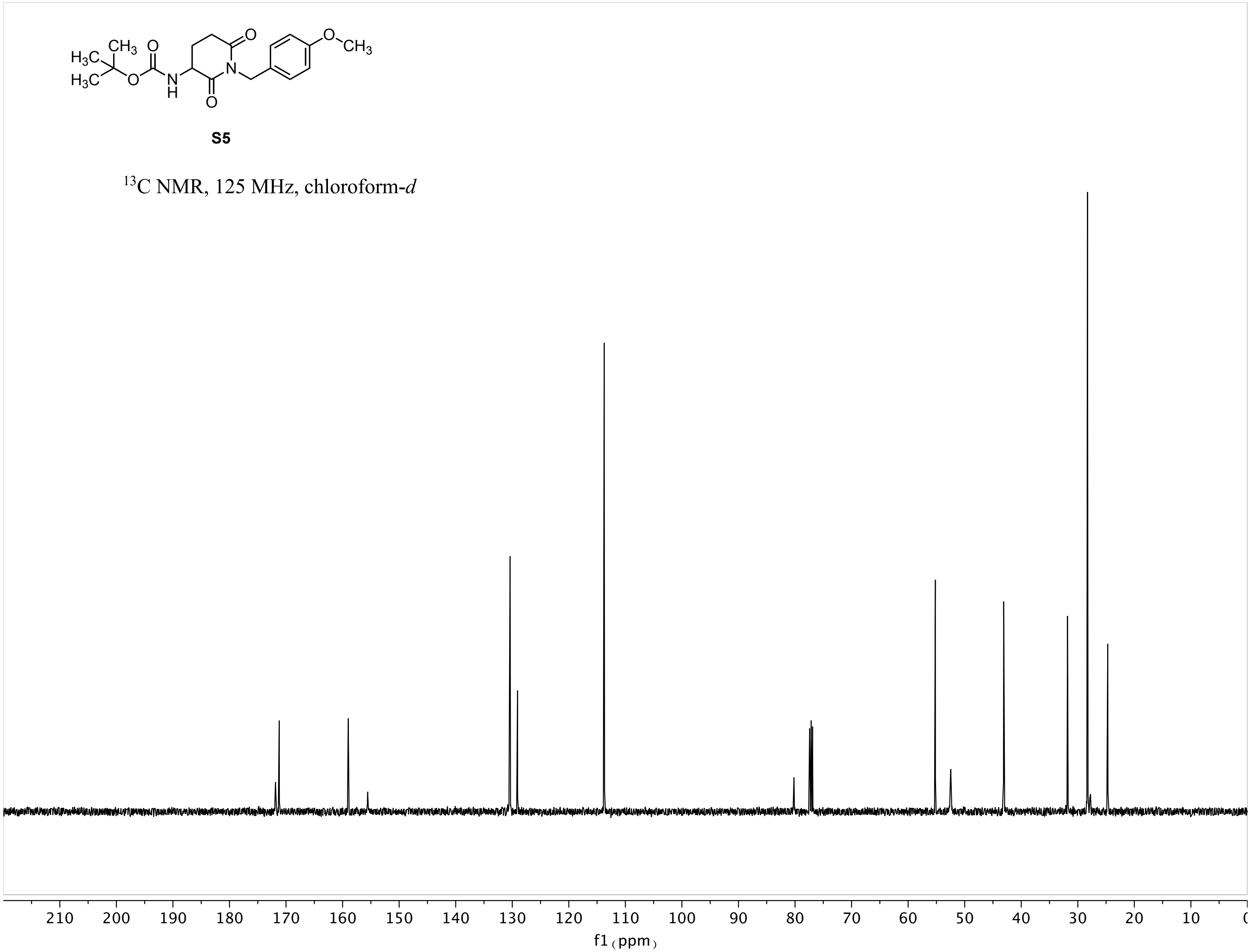
S5
$^{13}C$ NMR, 125 MHz, chloroform-*d*
210 200 190 180 170 160 150 140 130 120 110 100 90 80 70 60 50 40 30 20 10
f1 (ppm)

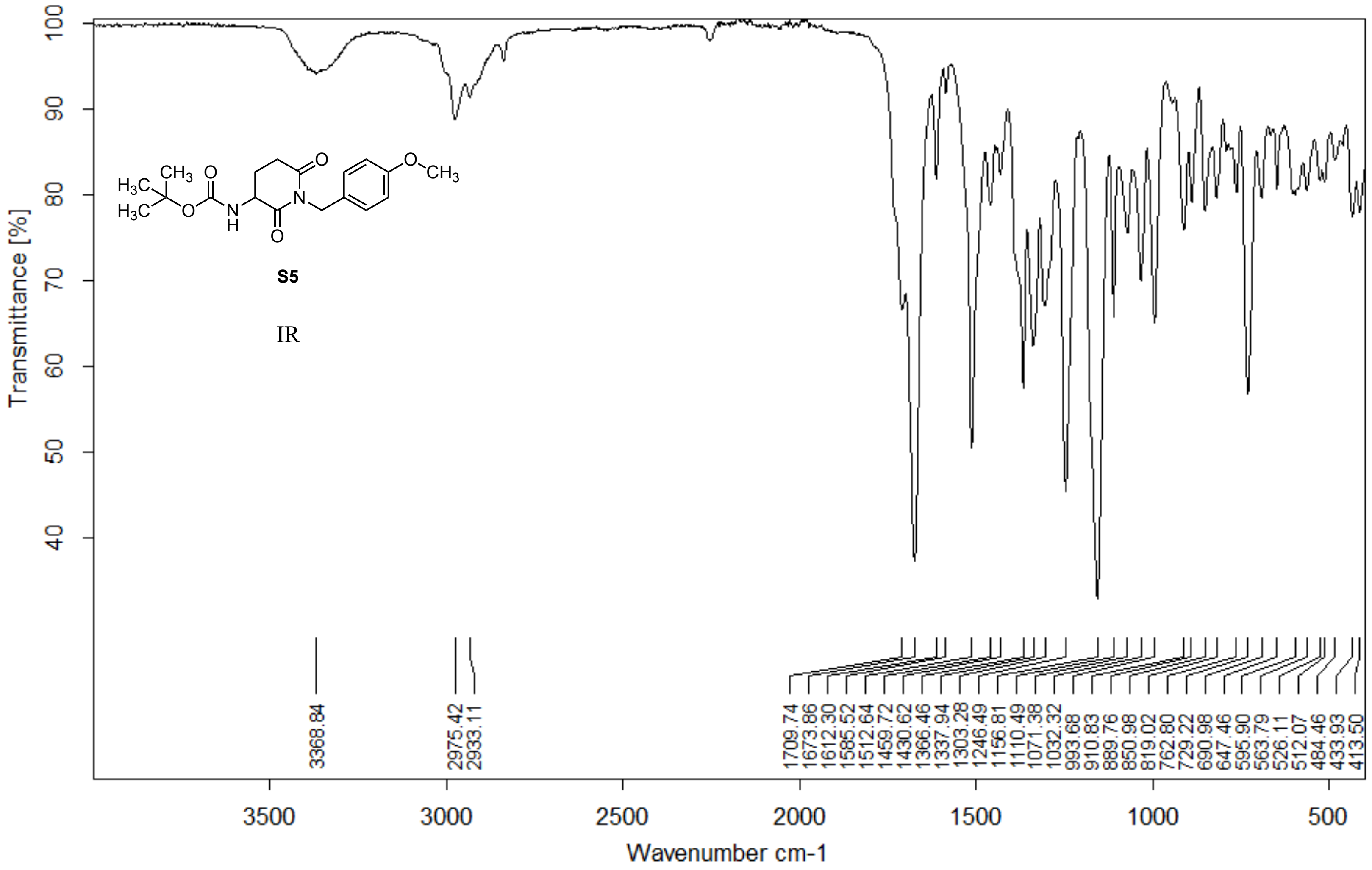


| | | | |
|---|---|---|---|
| Z:\Alpha FTIR Data\Woo\Yuka\YA503.0 | YA503 | Diamond ATR | 27/02/2020 |

**Page 1/2**

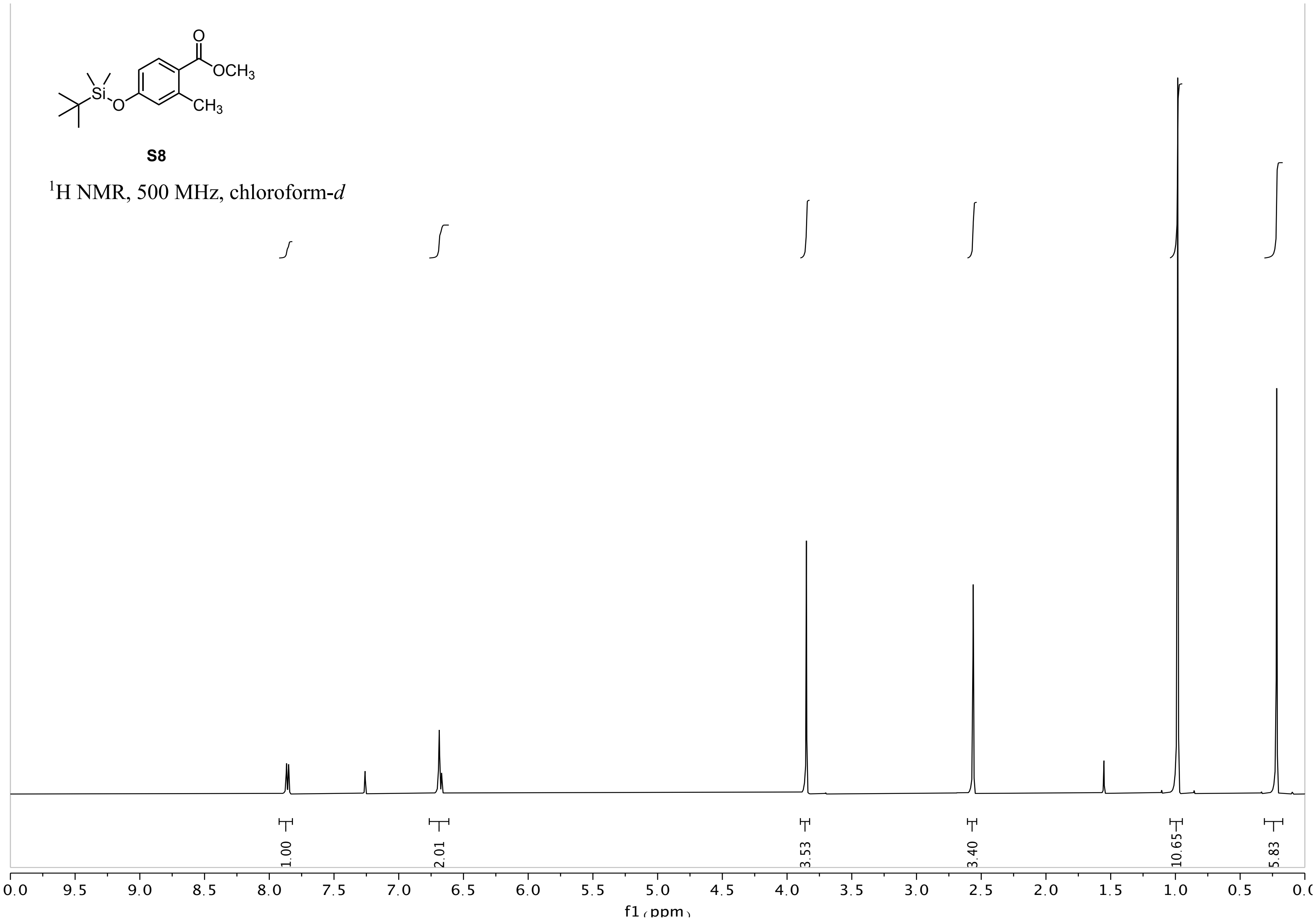
OCH3
CH3
Si
O
S8
1H NMR, 500 MHz, chloroform-d
1.00
2.01
3.53
3.40
10.65
5.83
f1 (ppm)

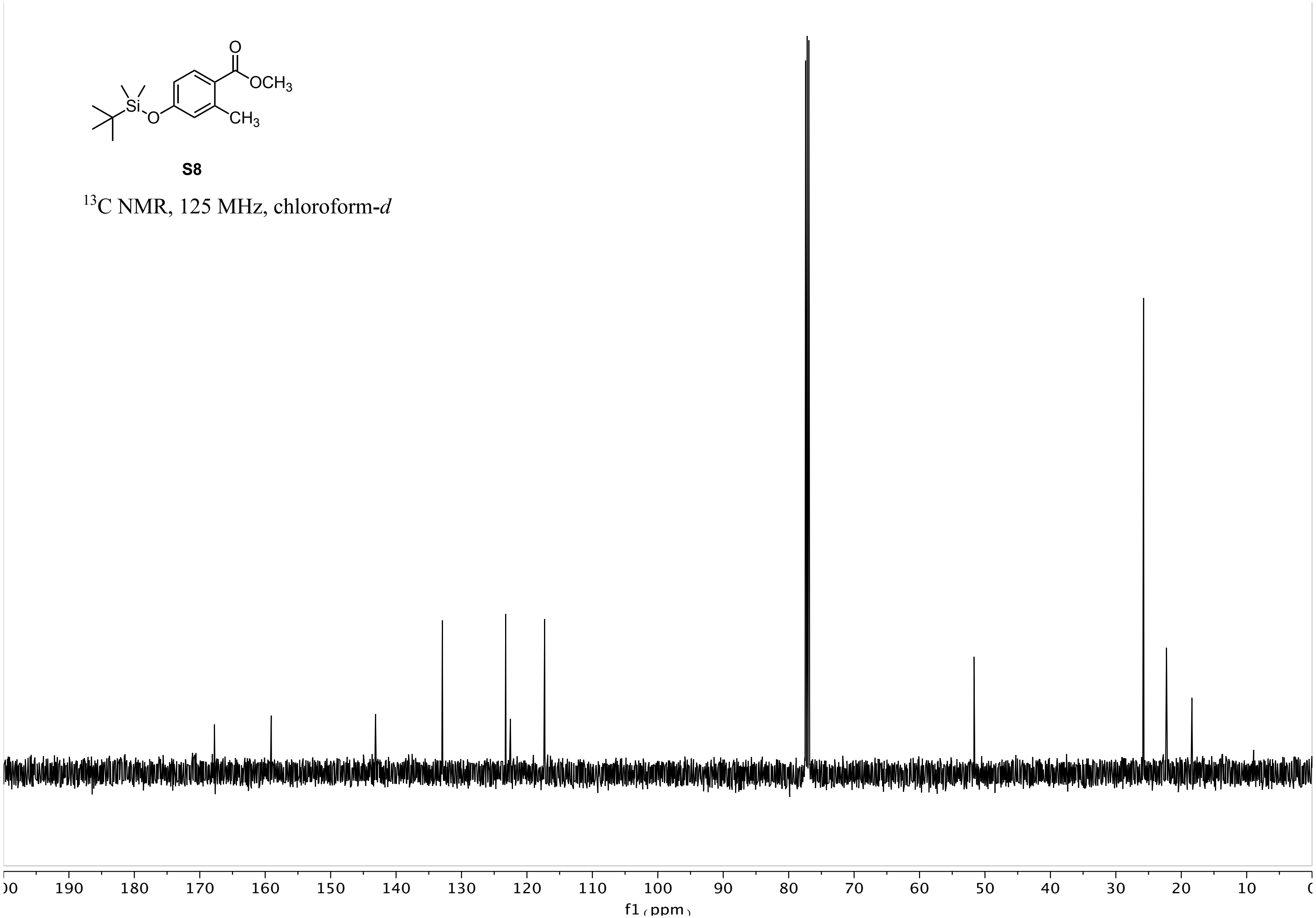
OCH3
CH3
Si
O
S8
13C NMR, 125 MHz, chloroform-d
190 180 170 160 150 140 130 120 110 100 90 80 70 60 50 40 30 20 10
f1 (ppm)

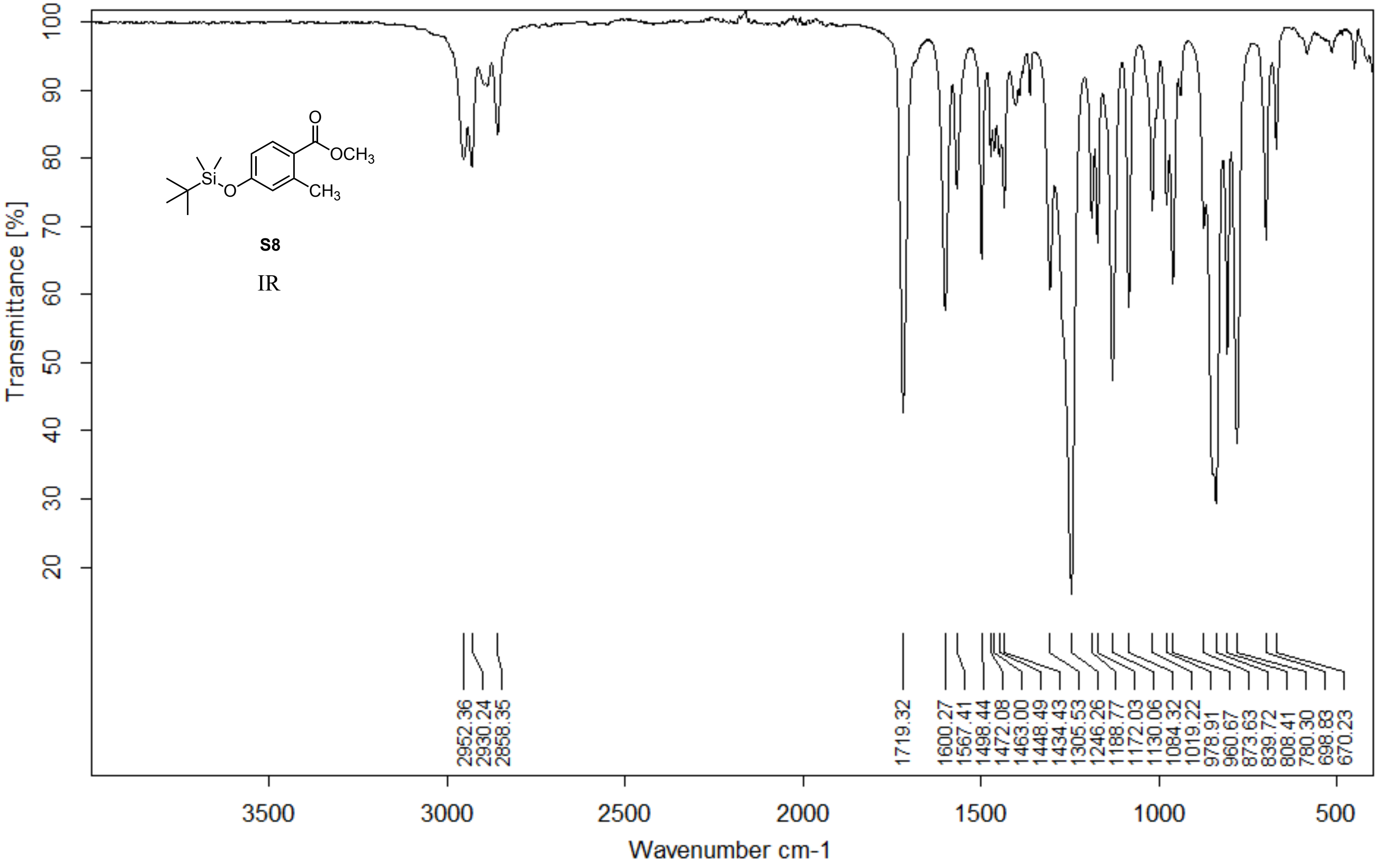


| | | | | |
|---|---|---|---|---|
| Z:\ALPHA FTIR DATA\Woo\Yuka\YA513.0 | YA513 | Diamond ATR | | 06/03/2020 |

**Page 1/2**

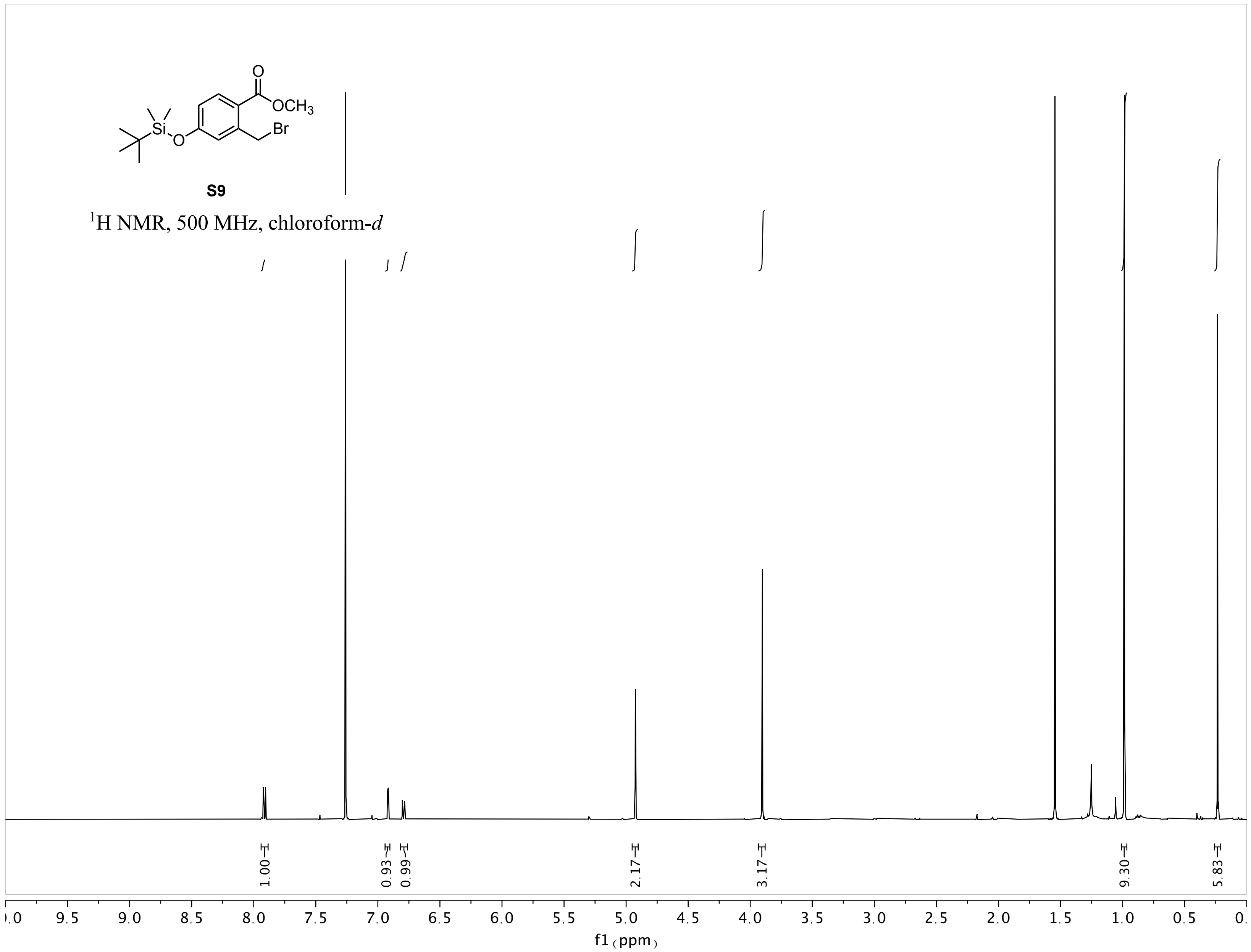
OCH3
Br
Si
O
S9
1H NMR, 500 MHz, chloroform-d
1.00
0.93
0.99
2.17
3.17
9.30
5.83
f1 (ppm)

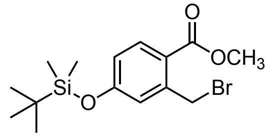


**S9**

$^{13}$C NMR, 125 MHz, chloroform-*d*

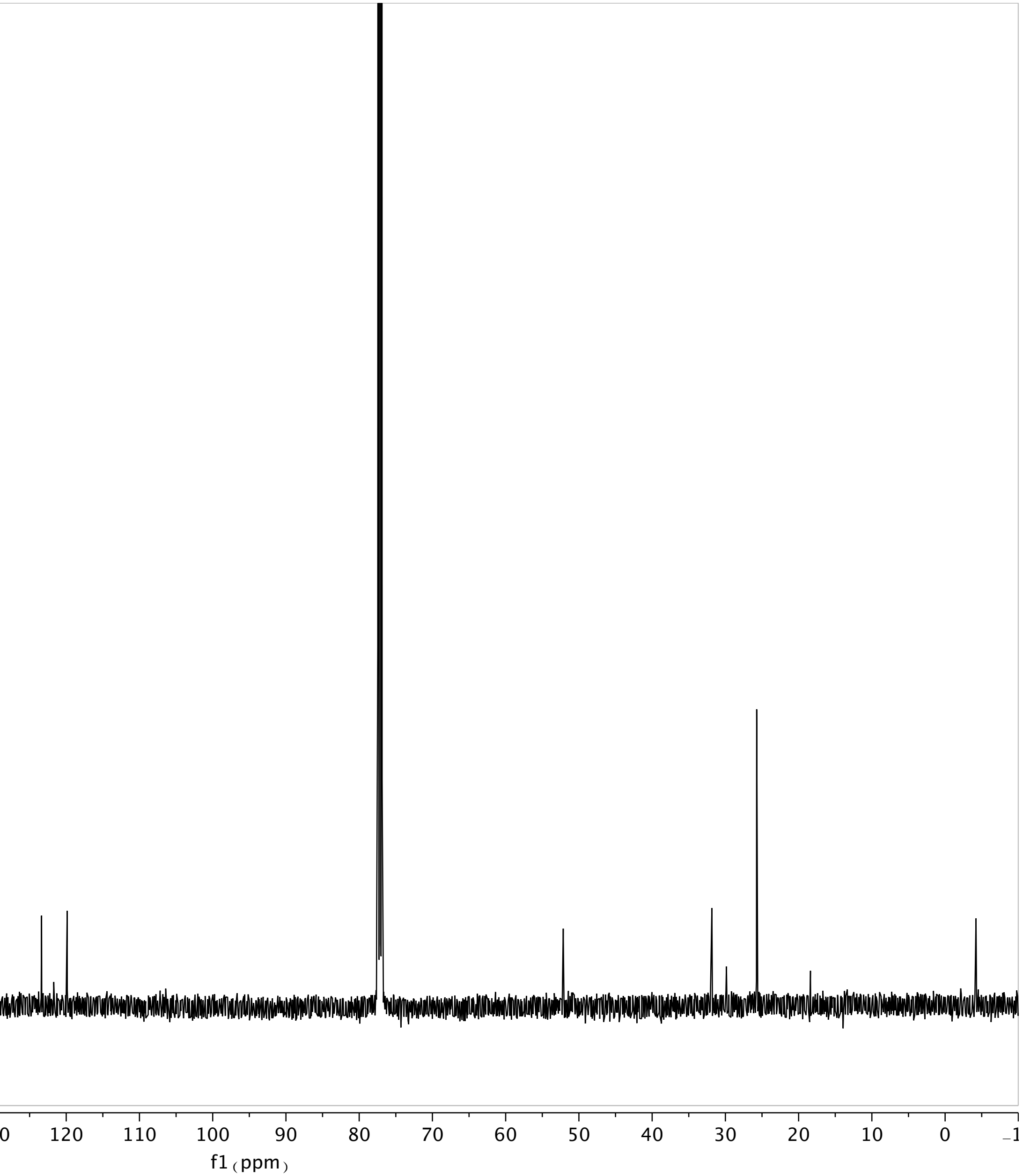

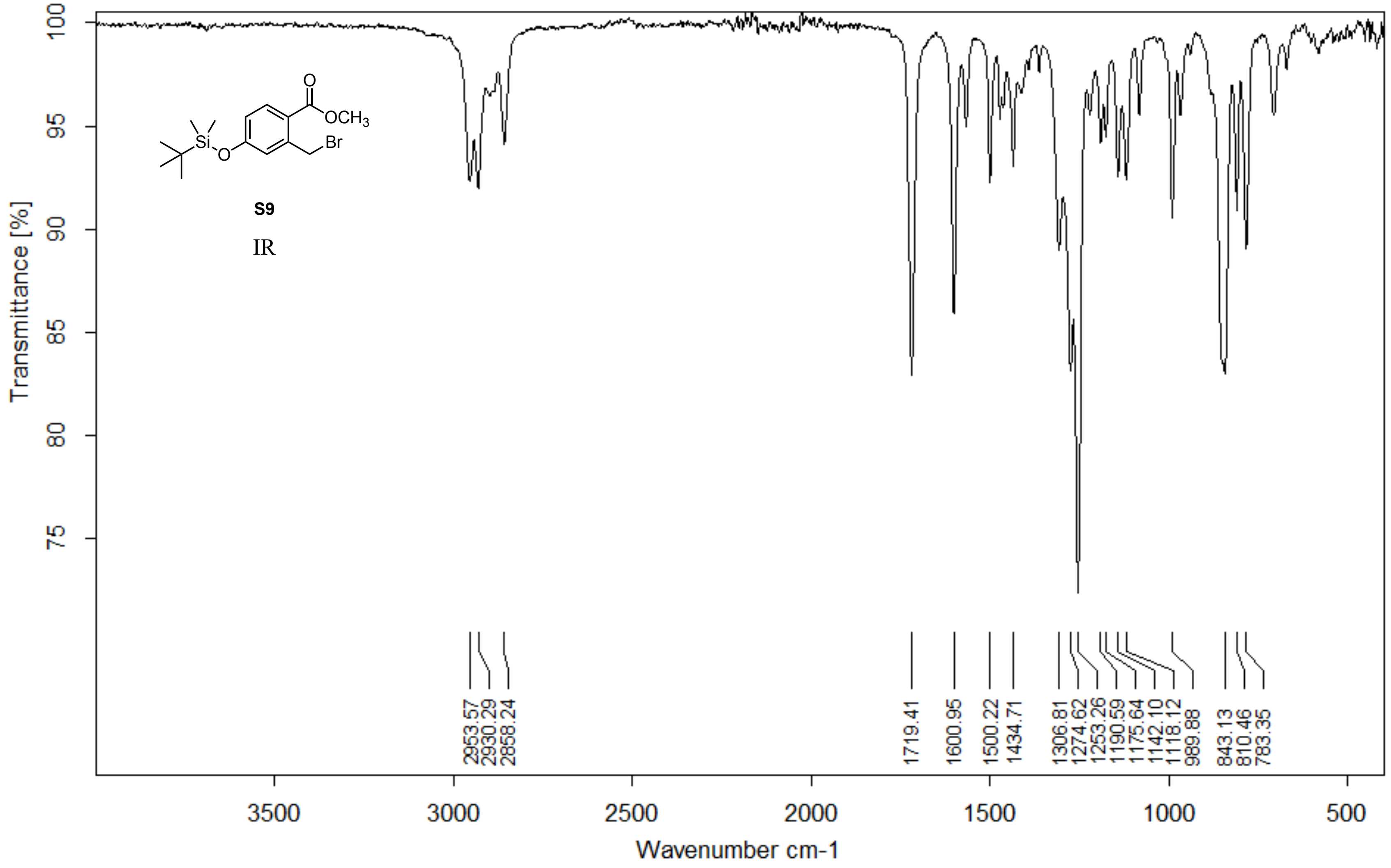


Z:\Alpha FTIR Data\Woo\Yuka\DKM-IV-A-134.0 DKM-IV-A-134 Diamond ATR

10/03/2020

**Page 1/2**

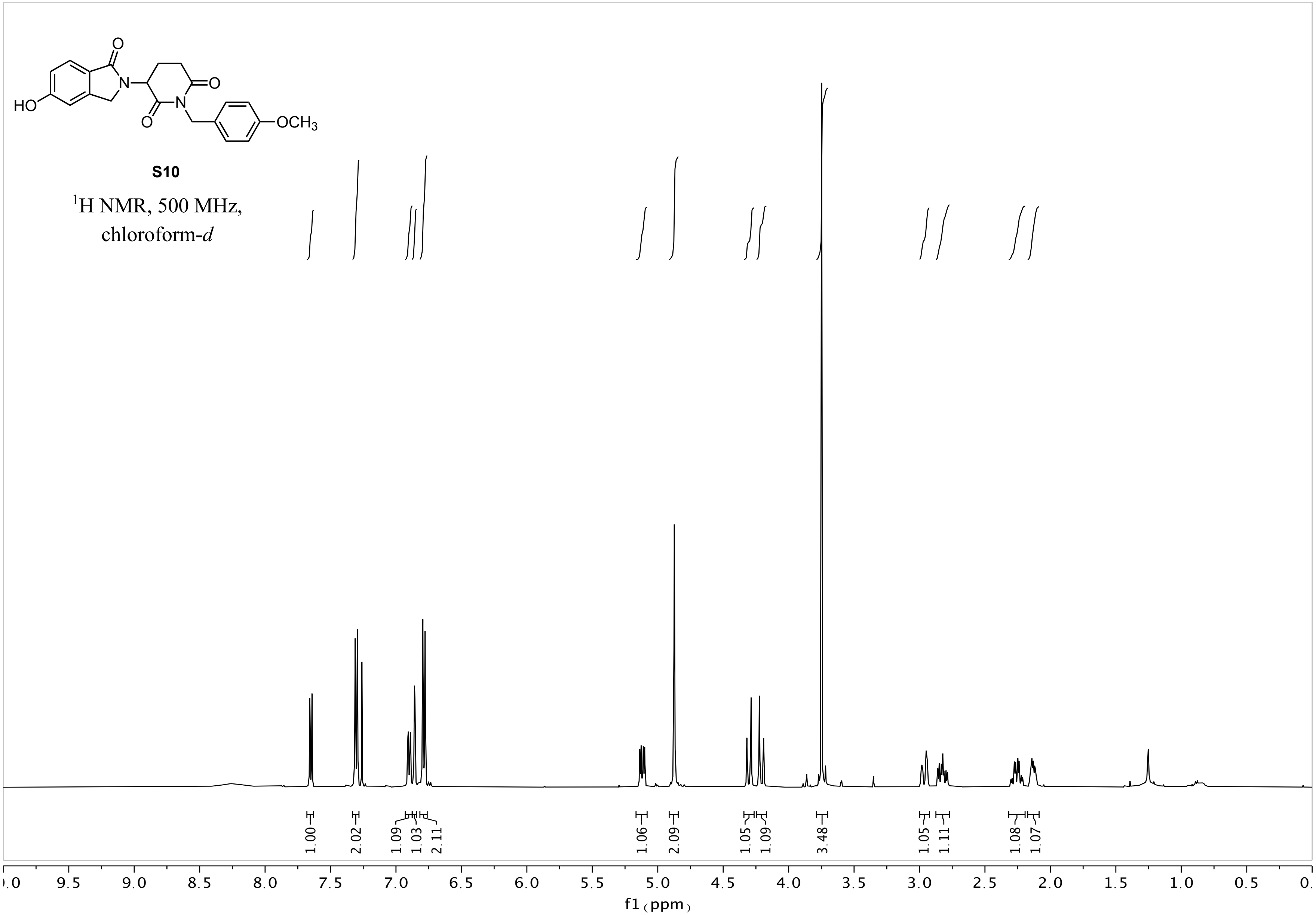
HO
N
O
O
N
O
OCH3
S10
1H NMR, 500 MHz,
chloroform-d
1.00
2.02
1.09
1.03
2.11
1.06
2.09
1.05
1.09
3.48
1.05
1.11
1.08
1.07
9.5
9.0
8.5
8.0
7.5
7.0
6.5
6.0
5.5
5.0
4.5
4.0
3.5
3.0
2.5
2.0
1.5
1.0
0.5
f1 (ppm)

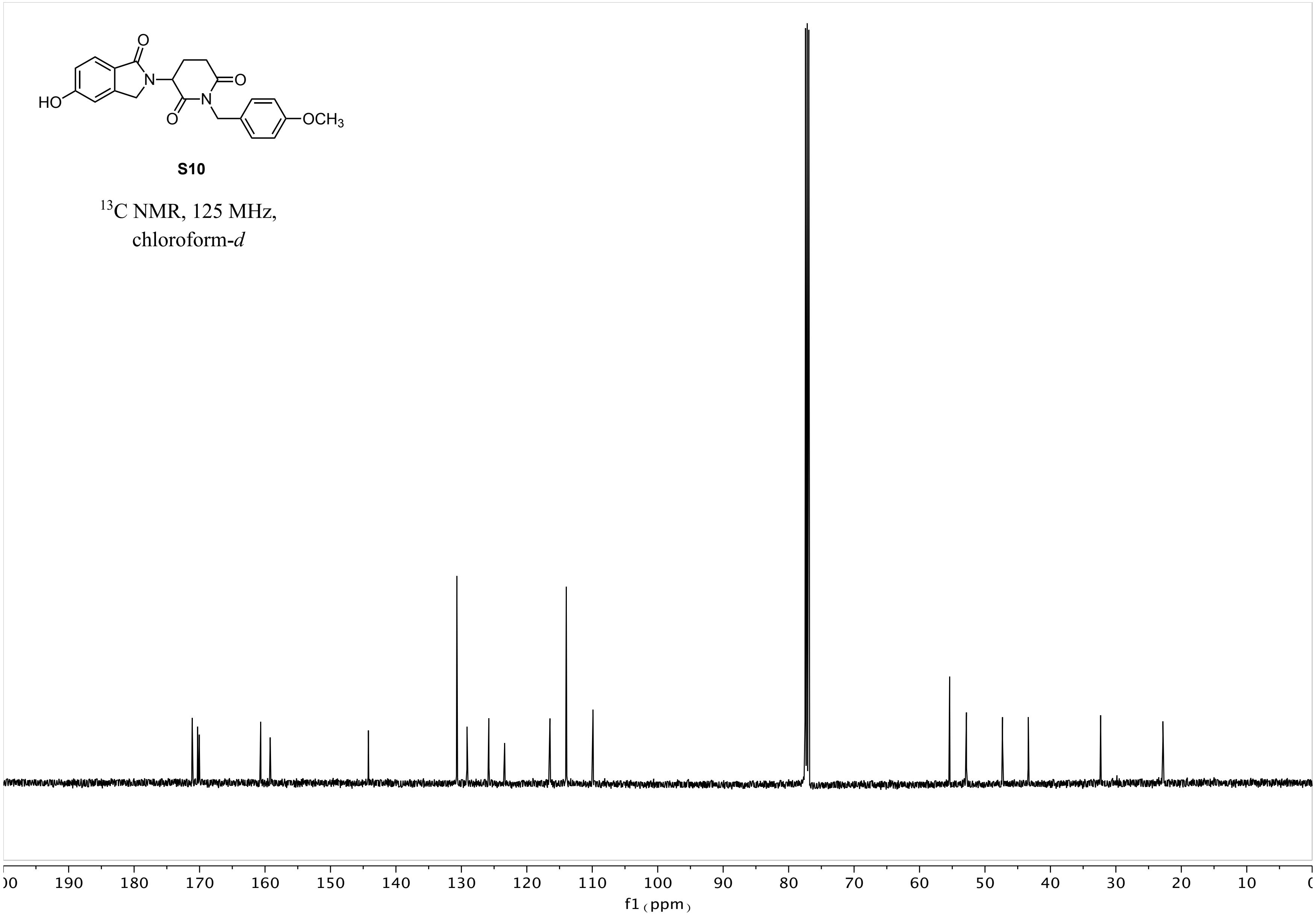
HO
N
O
O
N
O
OCH3
S10
13C NMR, 125 MHz, chloroform-d
190
180
170
160
150
140
130
120
110
100
90
80
70
60
50
40
30
20
10
f1 (ppm)

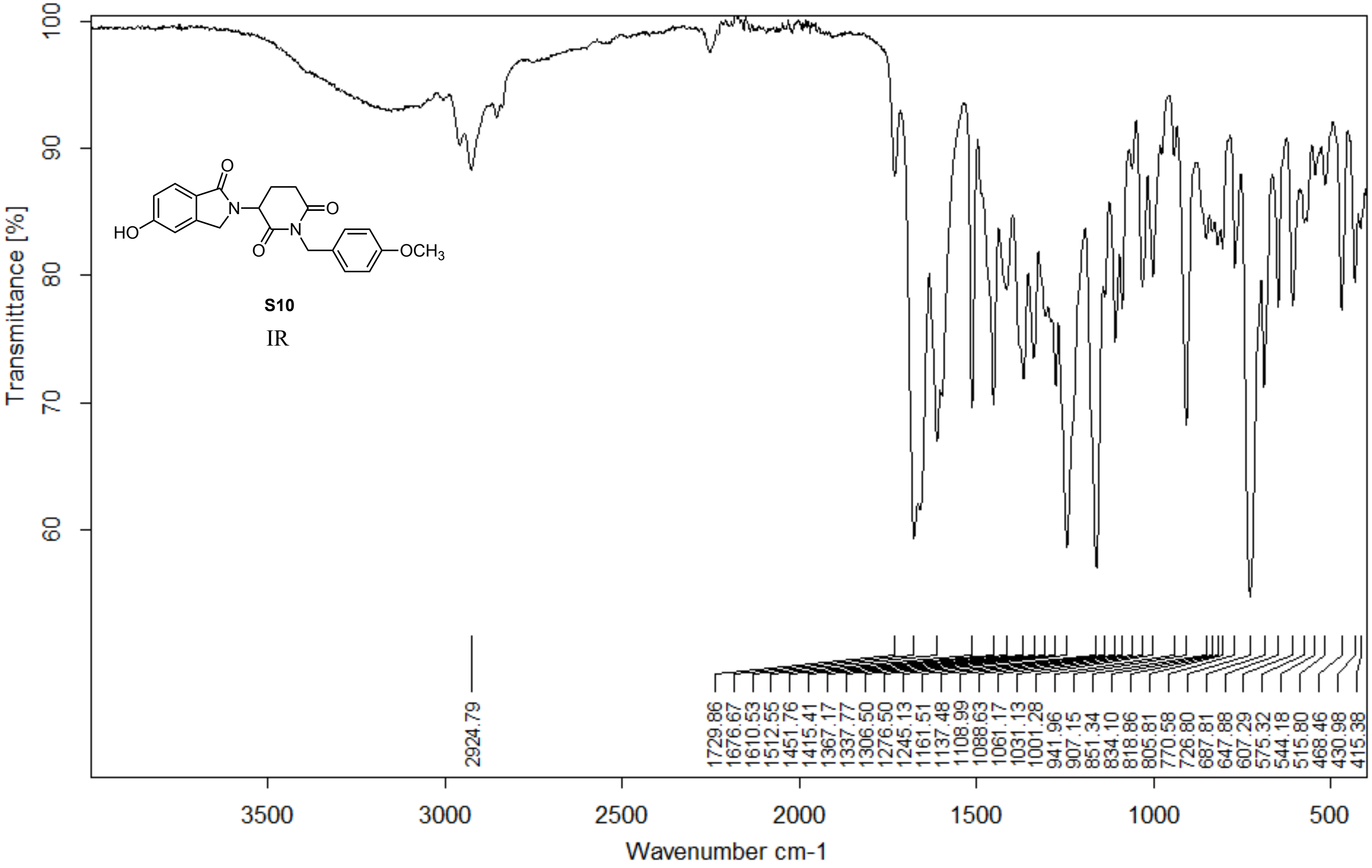


Z:\ALPHA FTIR DATA\Woo\Yuka\YA510.0 YA510 Diamond ATR

04/03/2020

**Page 1/2**

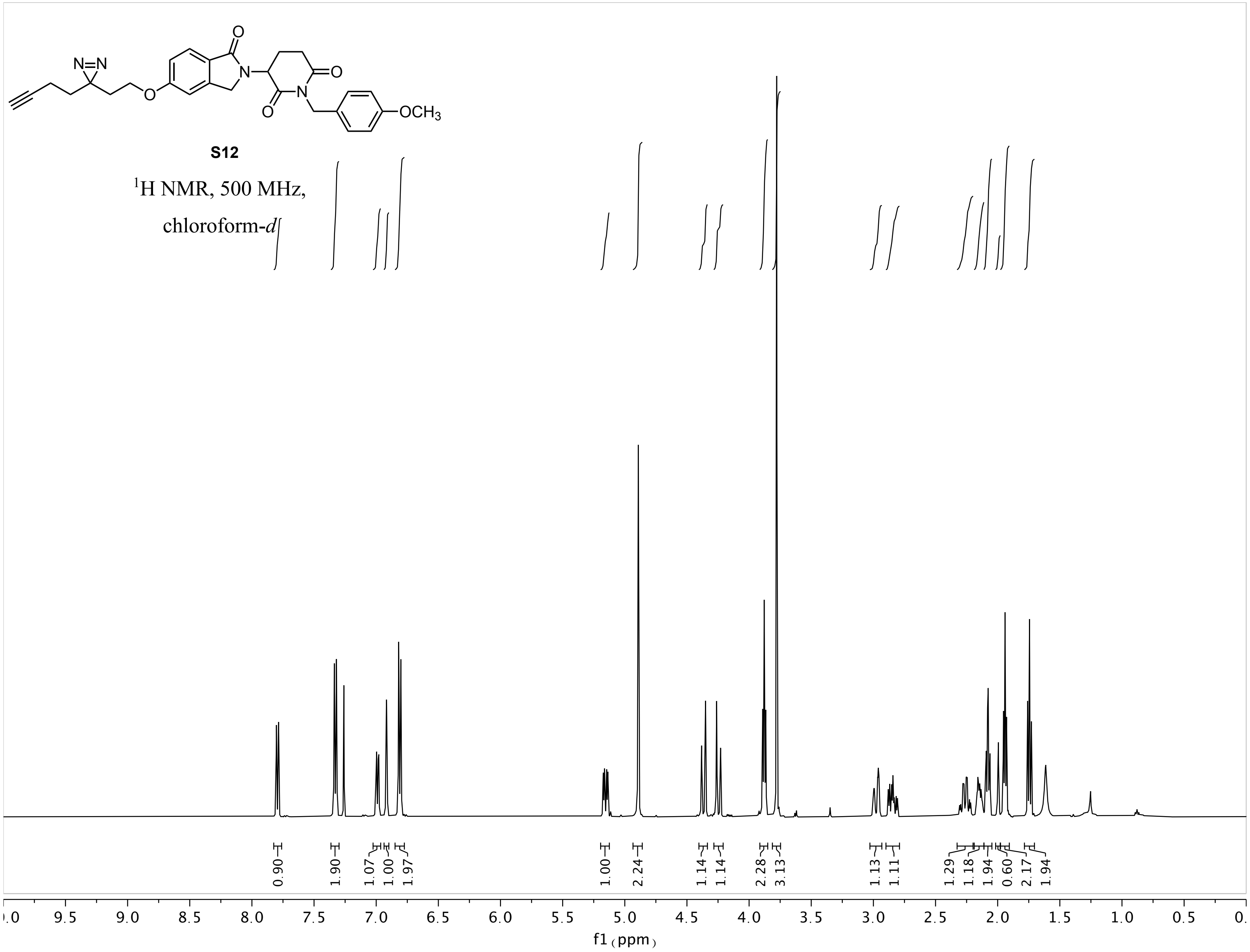
N=N
O
N
O
N
O
OCH3
S12
1H NMR, 500 MHz,
chloroform-d
0.90
1.90
1.07
1.00
1.97
1.00
2.24
1.14
1.14
2.28
3.13
1.13
1.11
1.29
1.18
1.94
0.60
2.17
1.94
9.5
9.0
8.5
8.0
7.5
7.0
6.5
6.0
5.5
5.0
4.5
4.0
3.5
3.0
2.5
2.0
1.5
1.0
0.5
f1 (ppm)

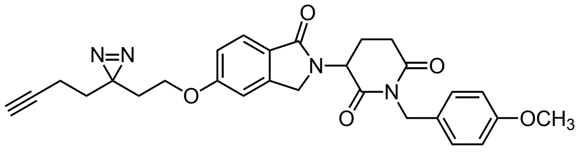

**S12**

$^{13}$C NMR, 125 MHz,

chloroform-*d*

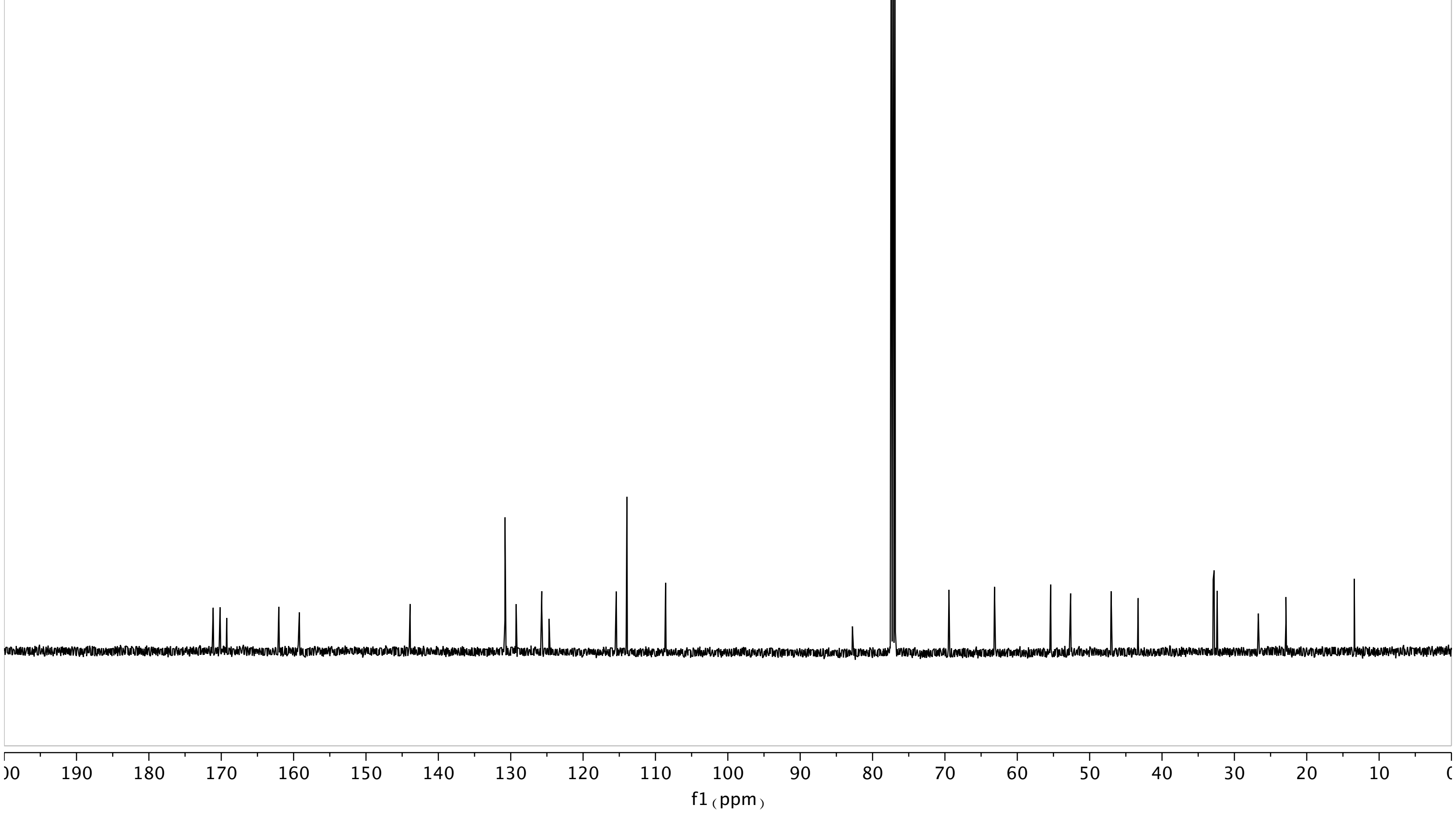

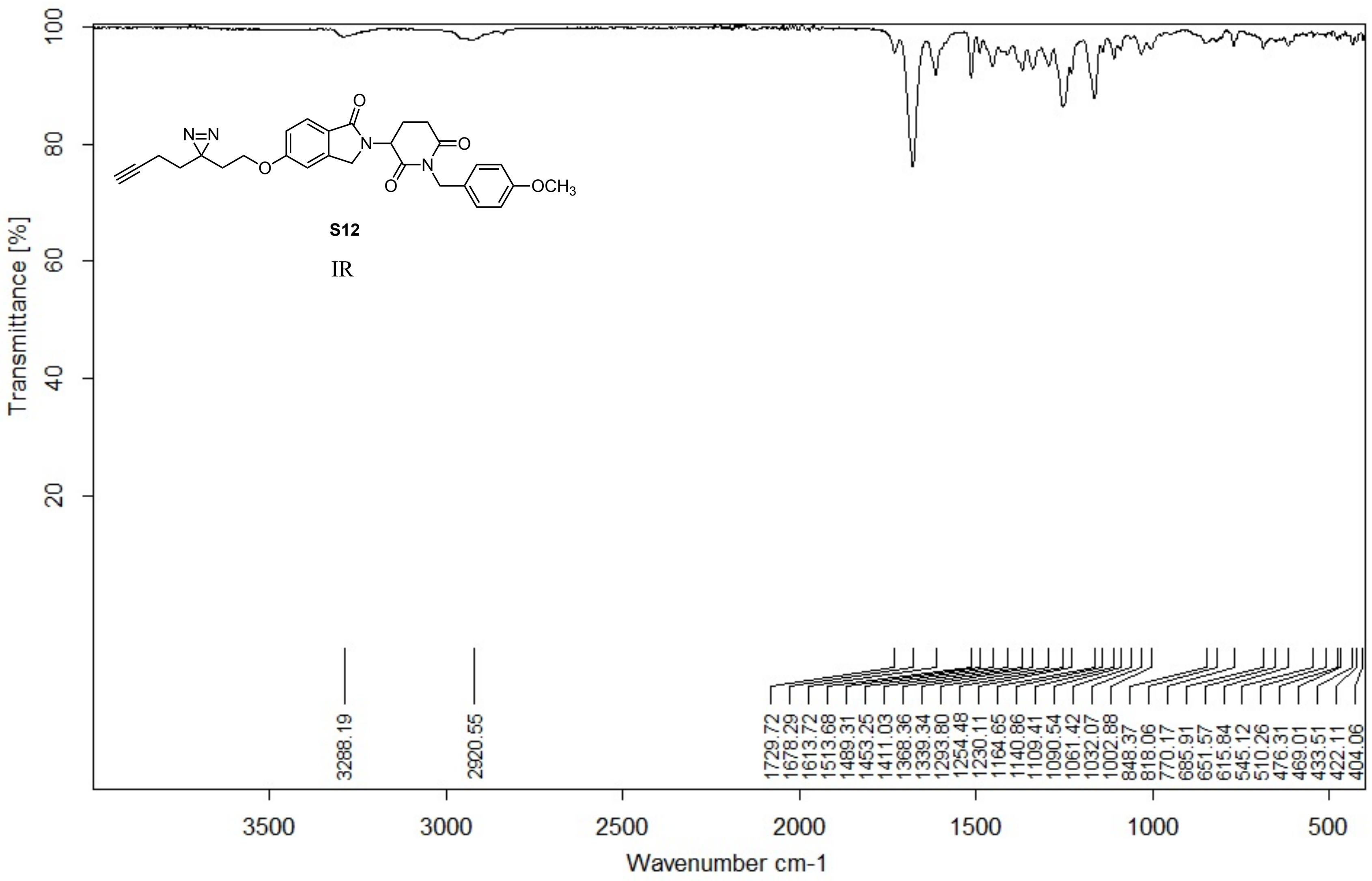


Z:\Alpha FTIR Data\Shair\dhuang\YA336.1 YA336 Diamond ATR

02/06/2018

Page 1/2

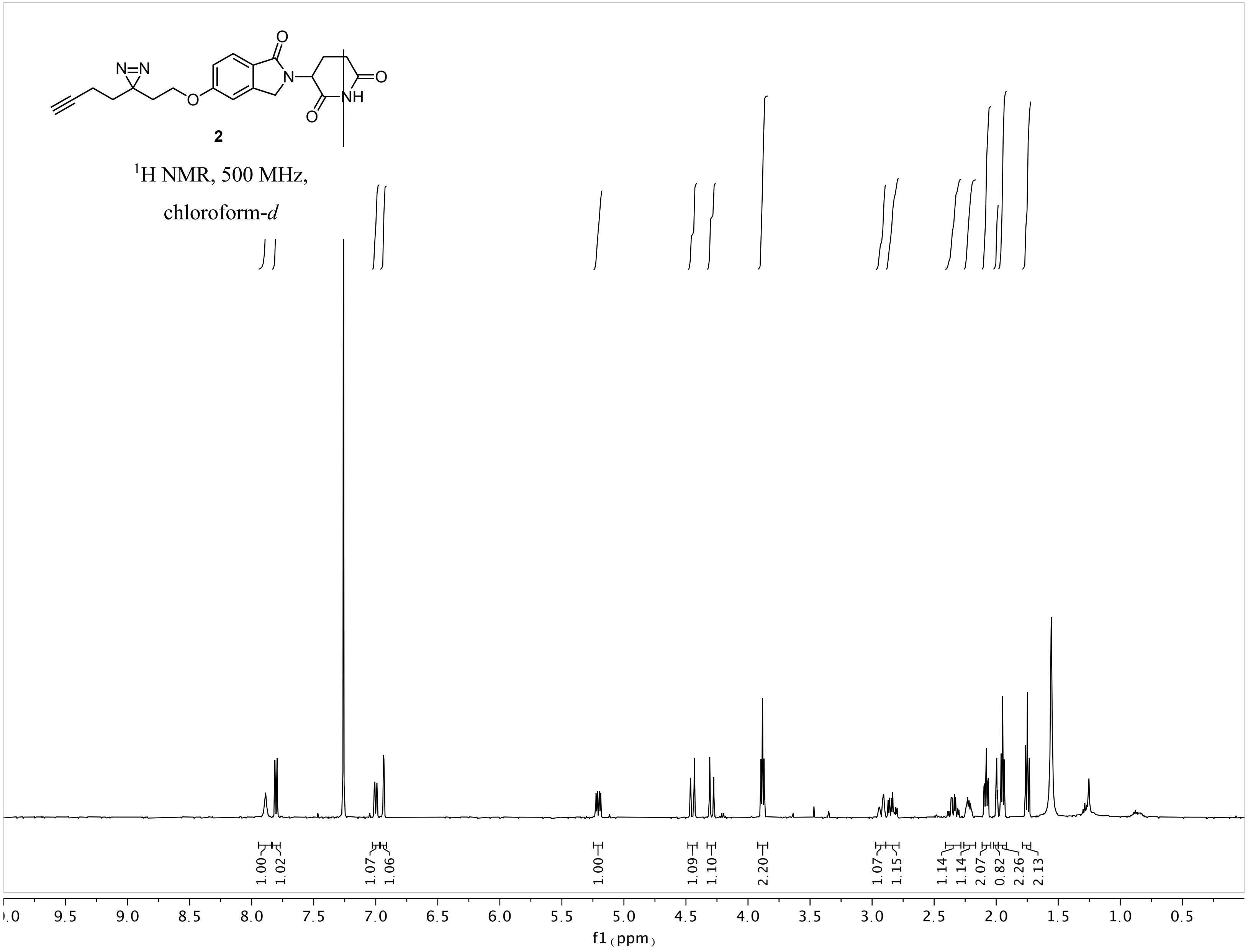
N=N
O
N
NH
2
$^1$H NMR, 500 MHz,
chloroform-*d*
1.00 1.02 1.07 1.06 1.00 1.09 1.10 2.20 1.07 1.15 1.14 1.14 2.07 0.82 2.26 2.13
9.5 9.0 8.5 8.0 7.5 7.0 6.5 6.0 5.5 5.0 4.5 4.0 3.5 3.0 2.5 2.0 1.5 1.0 0.5
f1 (ppm)

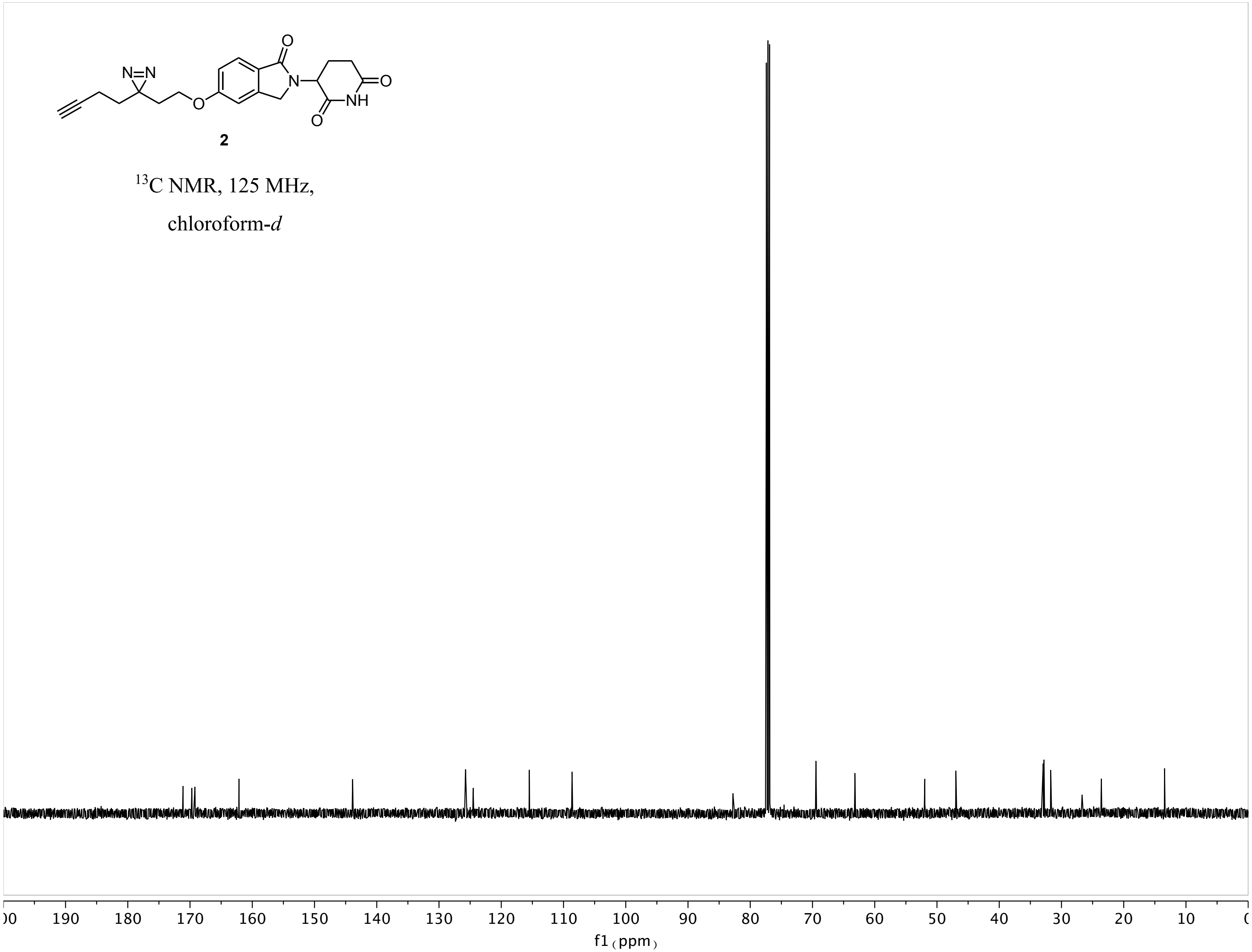
N=N
O
NH
2
13C NMR, 125 MHz,
chloroform-d
190 180 170 160 150 140 130 120 110 100 90 80 70 60 50 40 30 20 10
f1 (ppm)

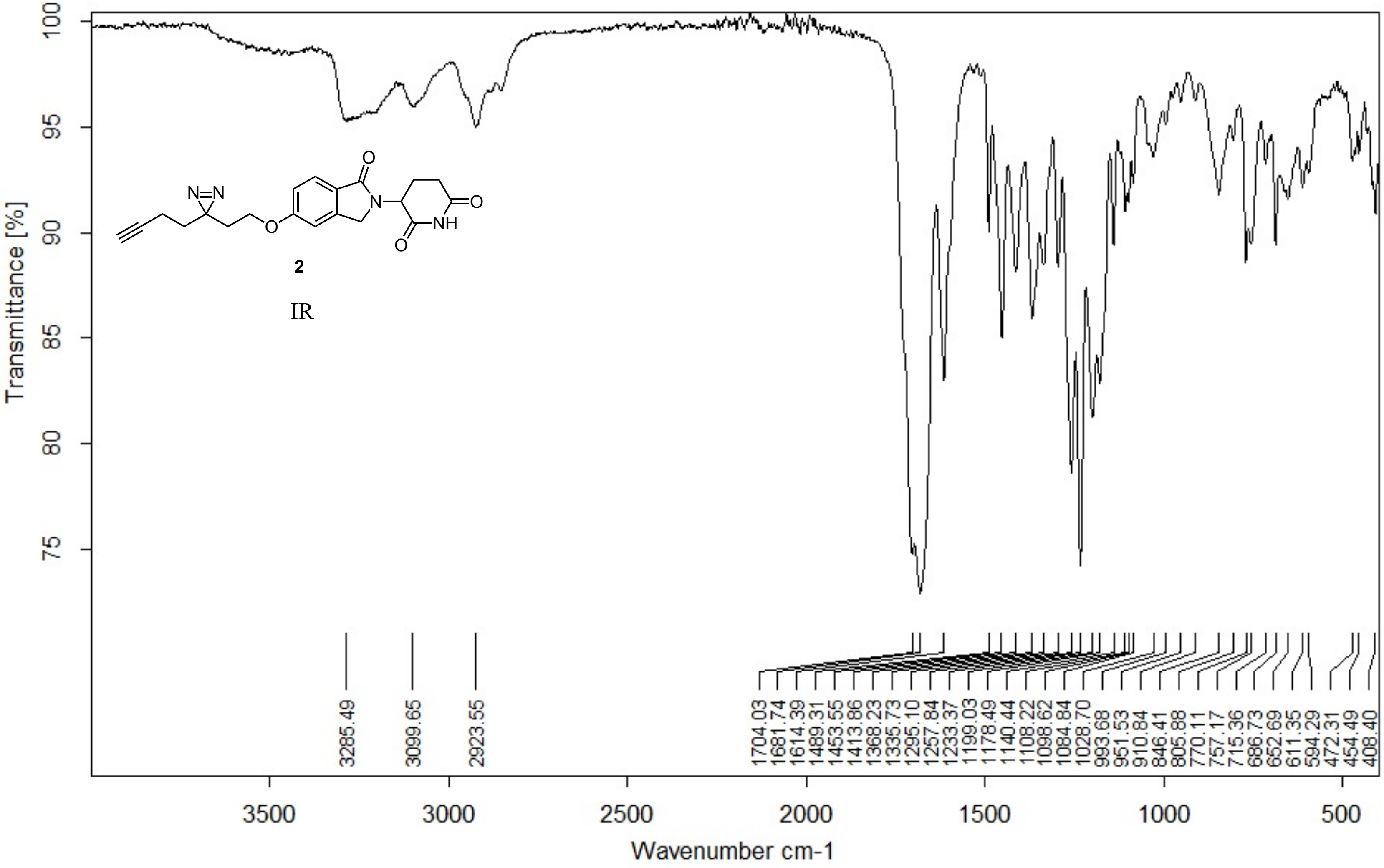


Z:\Alpha FTIR Data\Woo\Yuka\YA340.0 YA340 Diamond ATR

07/06/2018

**Page 1/2**

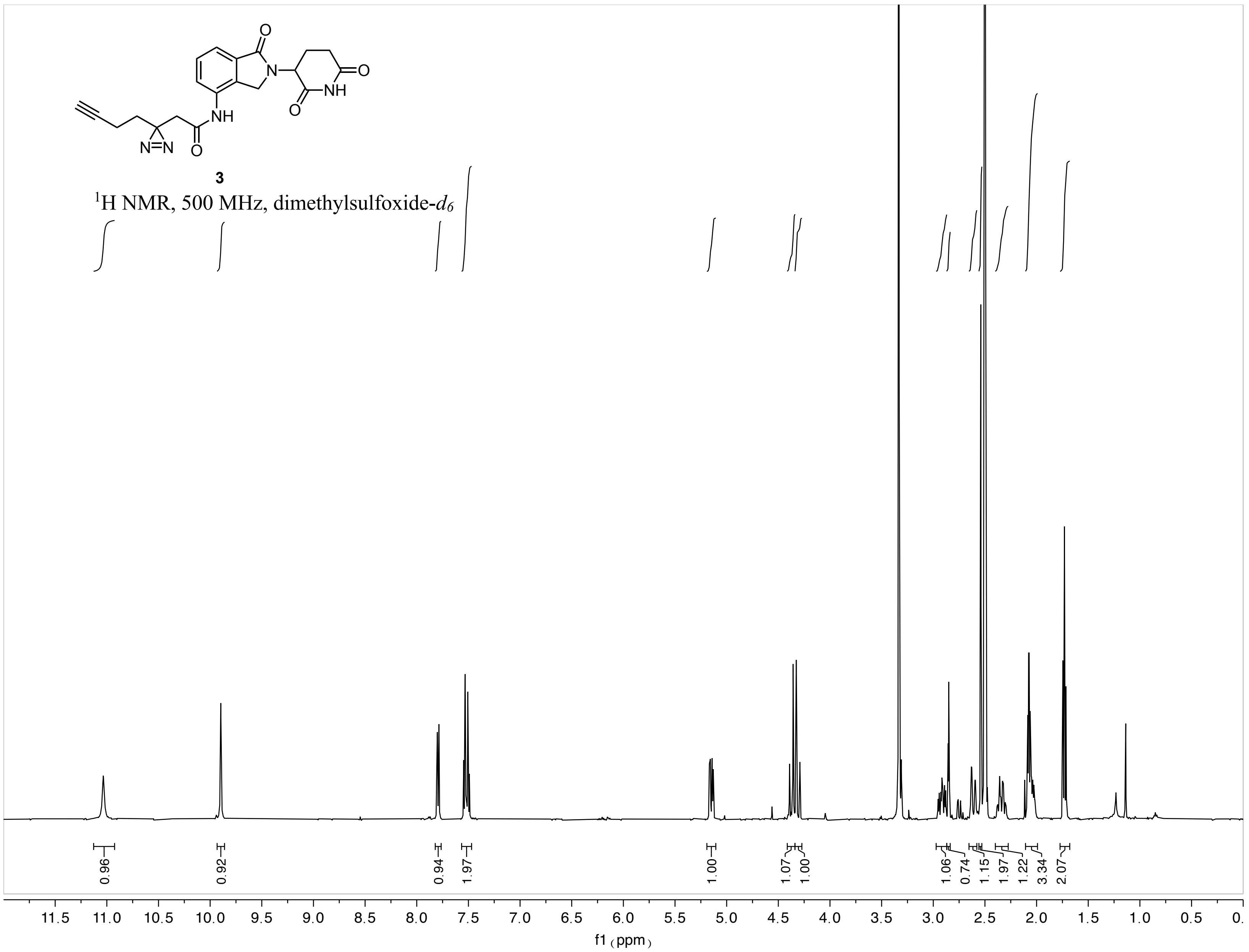
3
$^1$H NMR, 500 MHz, dimethylsulfoxide-$d_6$
0.96
0.92
0.94
1.97
1.00
1.07
1.00
1.06
0.74
1.15
1.97
1.22
3.34
2.07
11.5 11.0 10.5 10.0 9.5 9.0 8.5 8.0 7.5 7.0 6.5 6.0 5.5 5.0 4.5 4.0 3.5 3.0 2.5 2.0 1.5 1.0 0.5 0.
f1 (ppm)

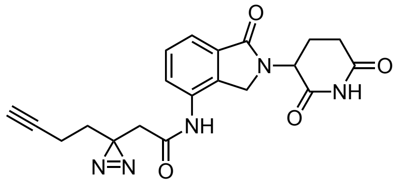

**3**

$^{13}$C NMR, 125 MHz, dimethylsulfoxide-$d_6$

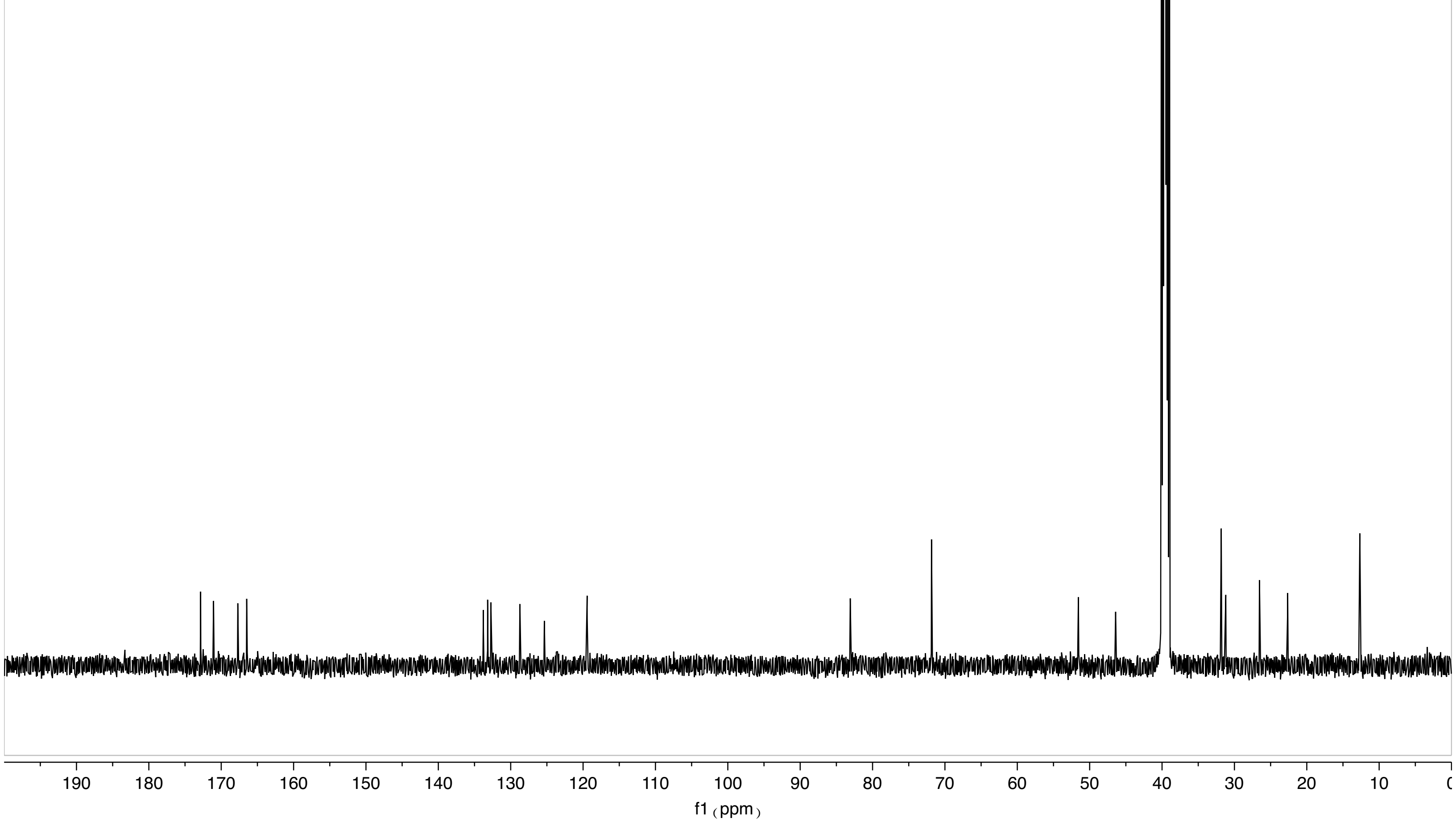

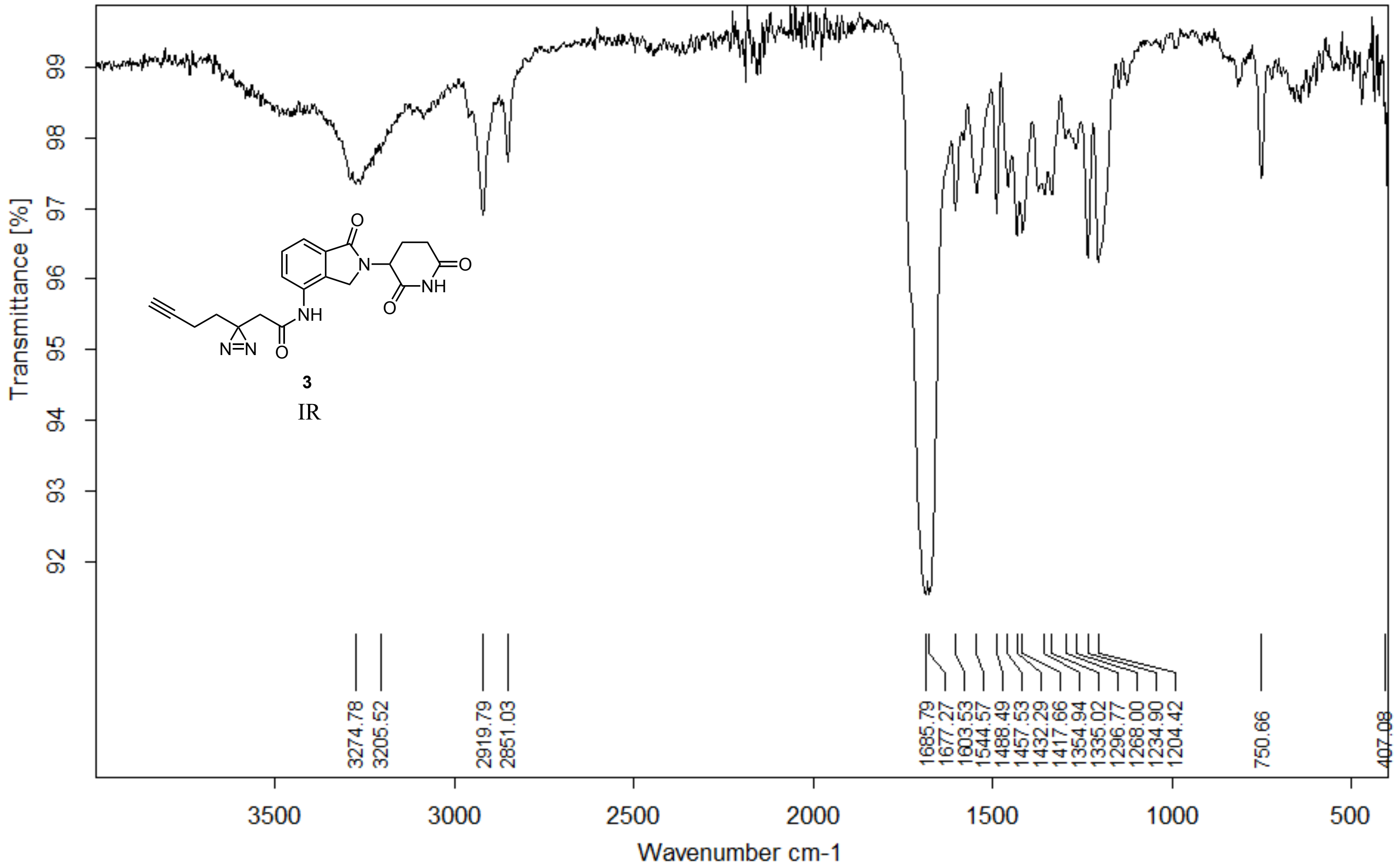


Z:\Alpha FTIR Data\Woo\Yuka\YA241 NpLEN.1 YA241 NpLEN Diamond ATR

09/06/2018

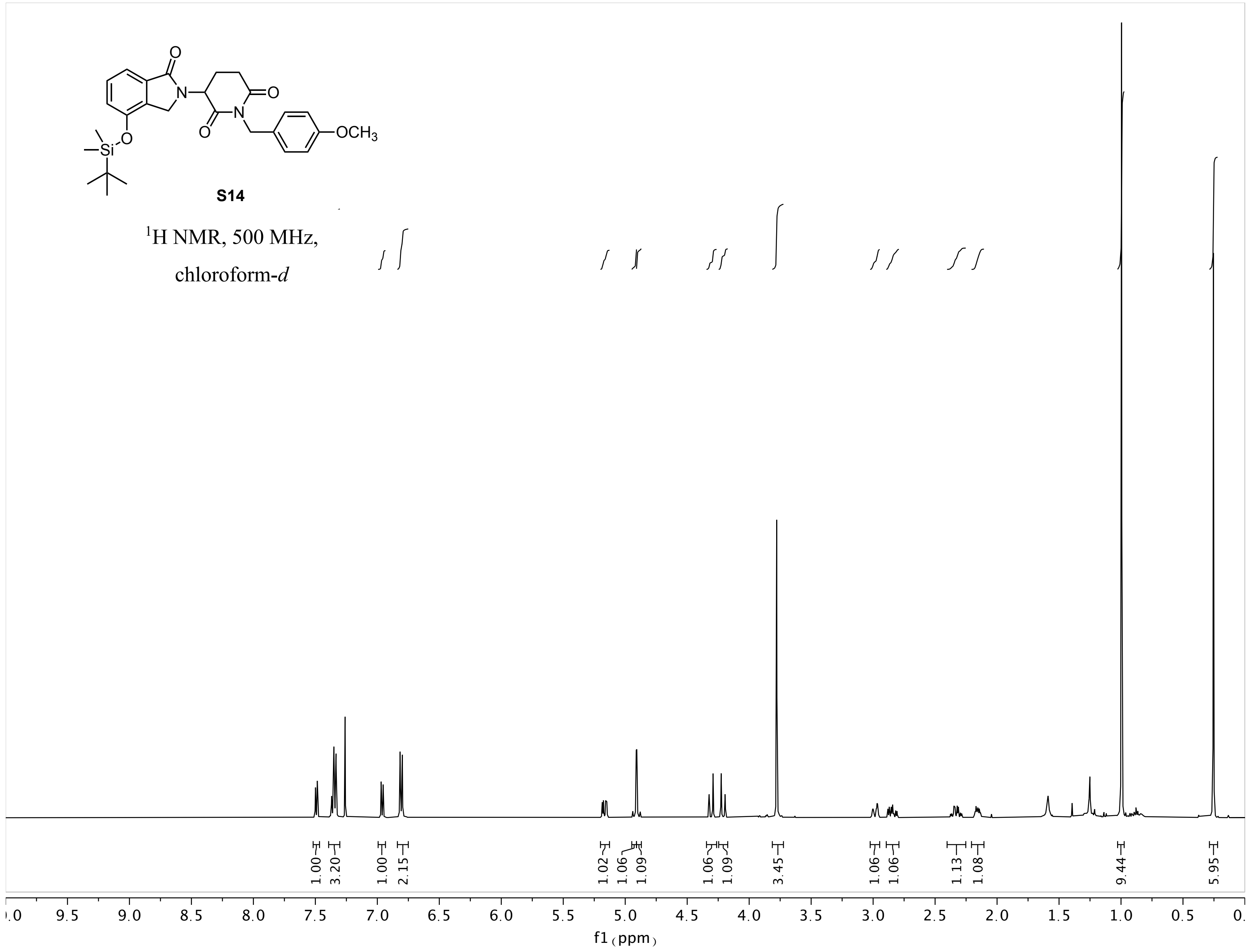
S14
$^{1}$H NMR, 500 MHz,
chloroform-$d$
1.00
3.20
1.00
2.15
1.02
1.06
1.09
1.06
1.09
3.45
1.06
1.06
1.13
1.08
9.44
5.95
9.5
9.0
8.5
8.0
7.5
7.0
6.5
6.0
5.5
5.0
4.5
4.0
3.5
3.0
2.5
2.0
1.5
1.0
0.5
f1 (ppm)

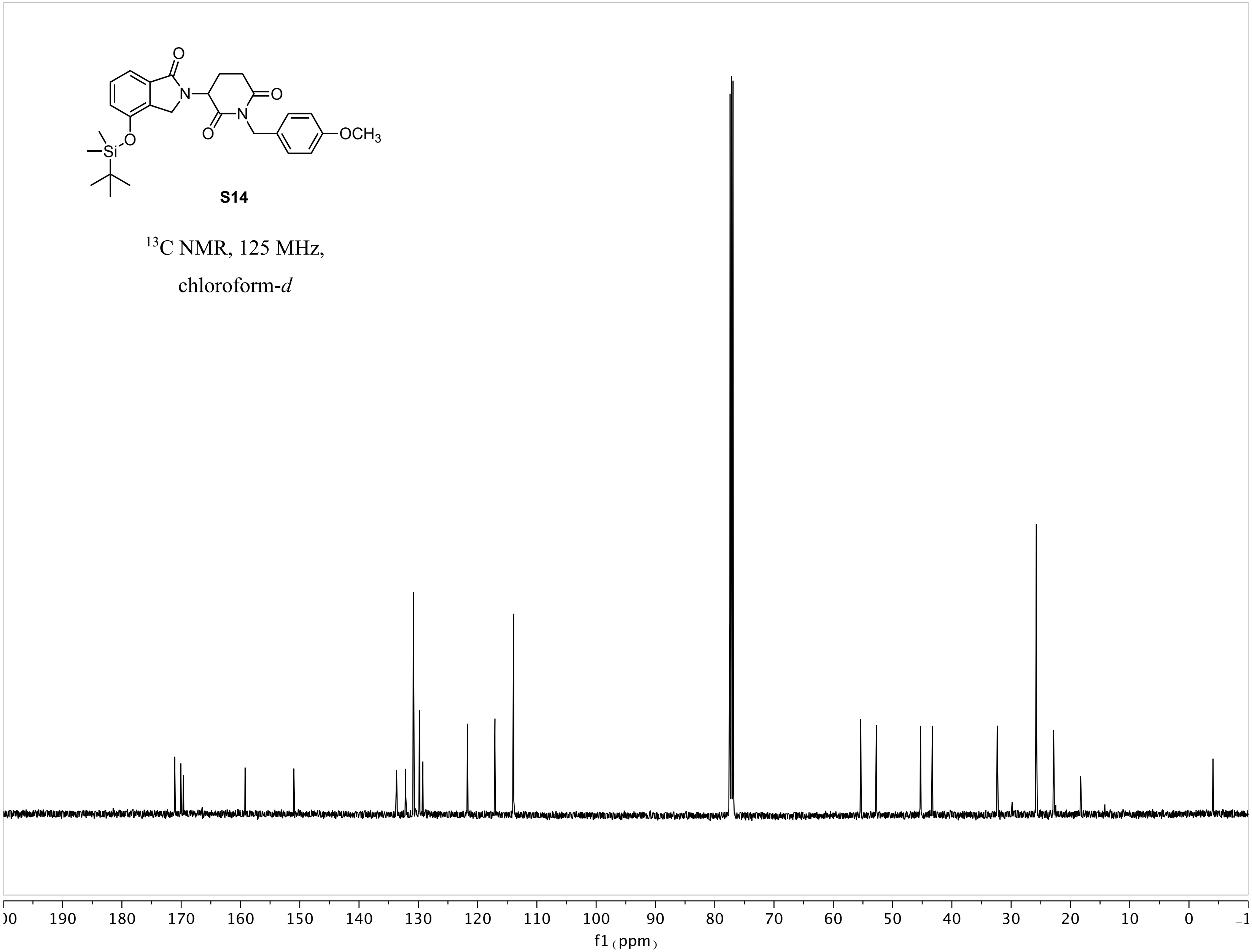
OCH3
S14
13C NMR, 125 MHz,
chloroform-d
190 180 170 160 150 140 130 120 110 100 90 80 70 60 50 40 30 20 10 0
f1 (ppm)

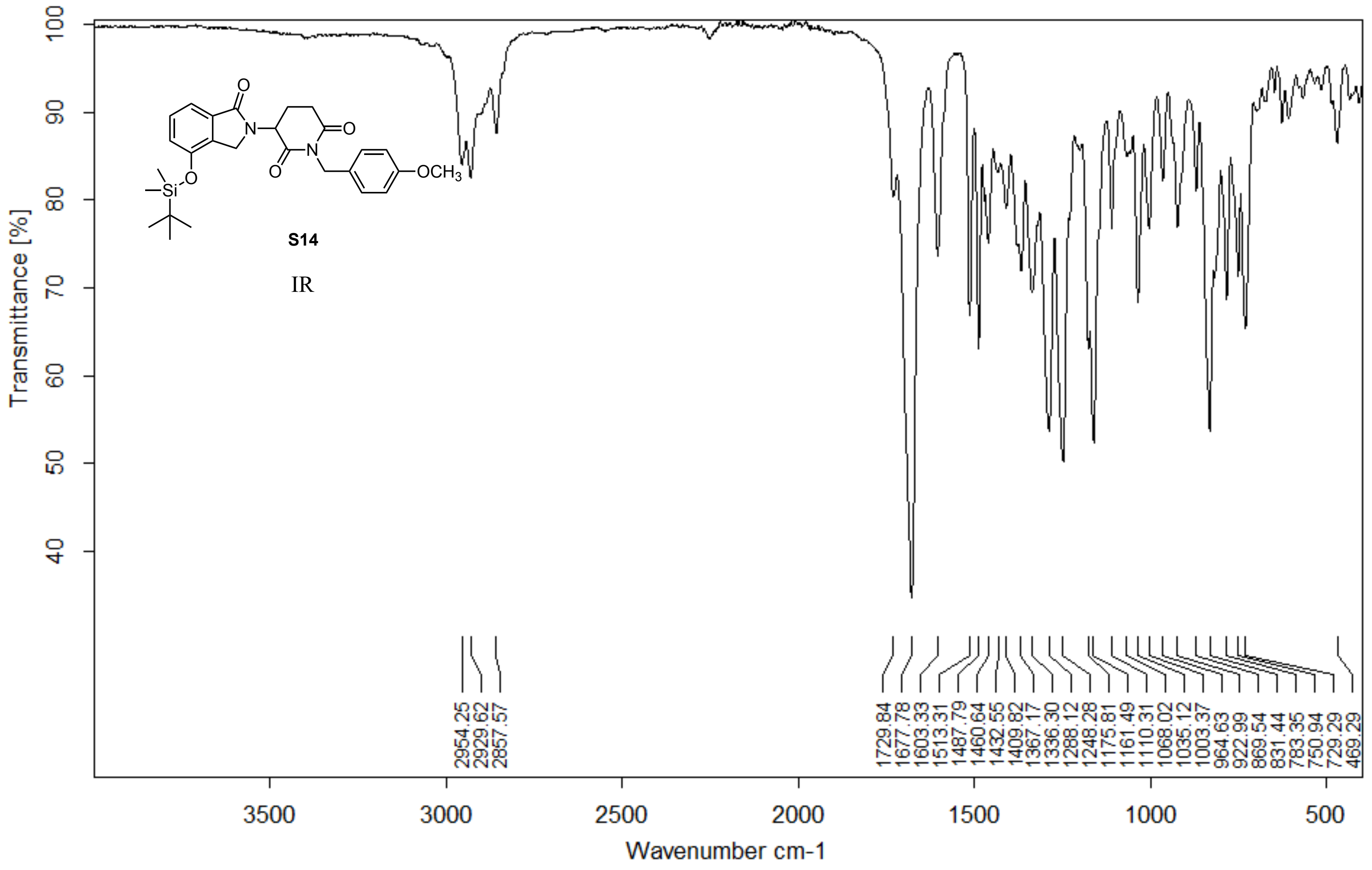


Z:\ALPHA FTIR DATA\Woo\Yuka\YA511.0 YA511 Diamond ATR

06/03/2020

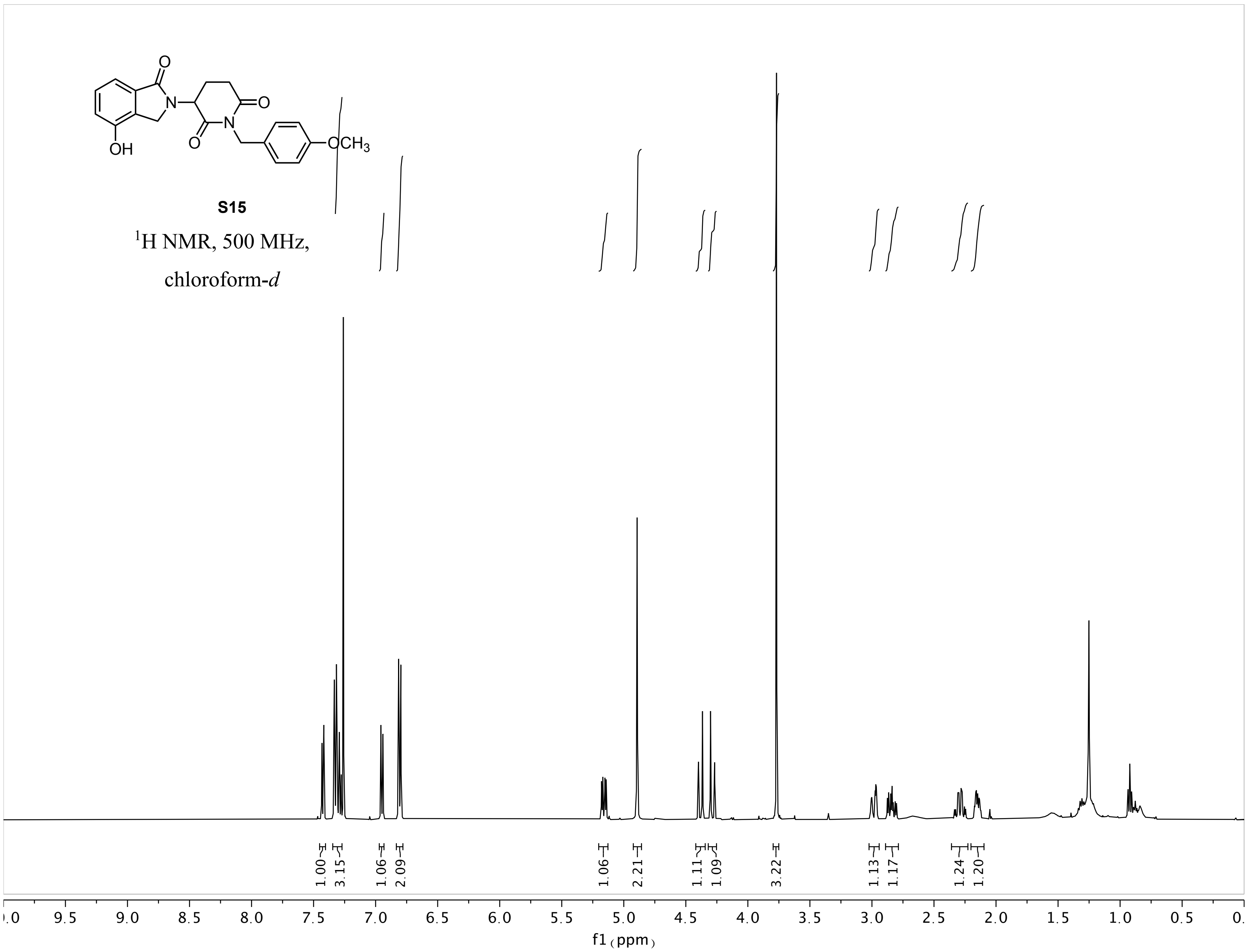

S15
1H NMR, 500 MHz,
chloroform-d
1.00
3.15
1.06
2.09
1.06
2.21
1.11
1.09
3.22
1.13
1.17
1.24
1.20
f1 (ppm)

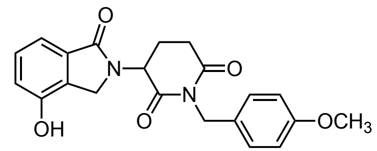


**S15**

$^{13}$C NMR, 125 MHz,

chloroform-*d*

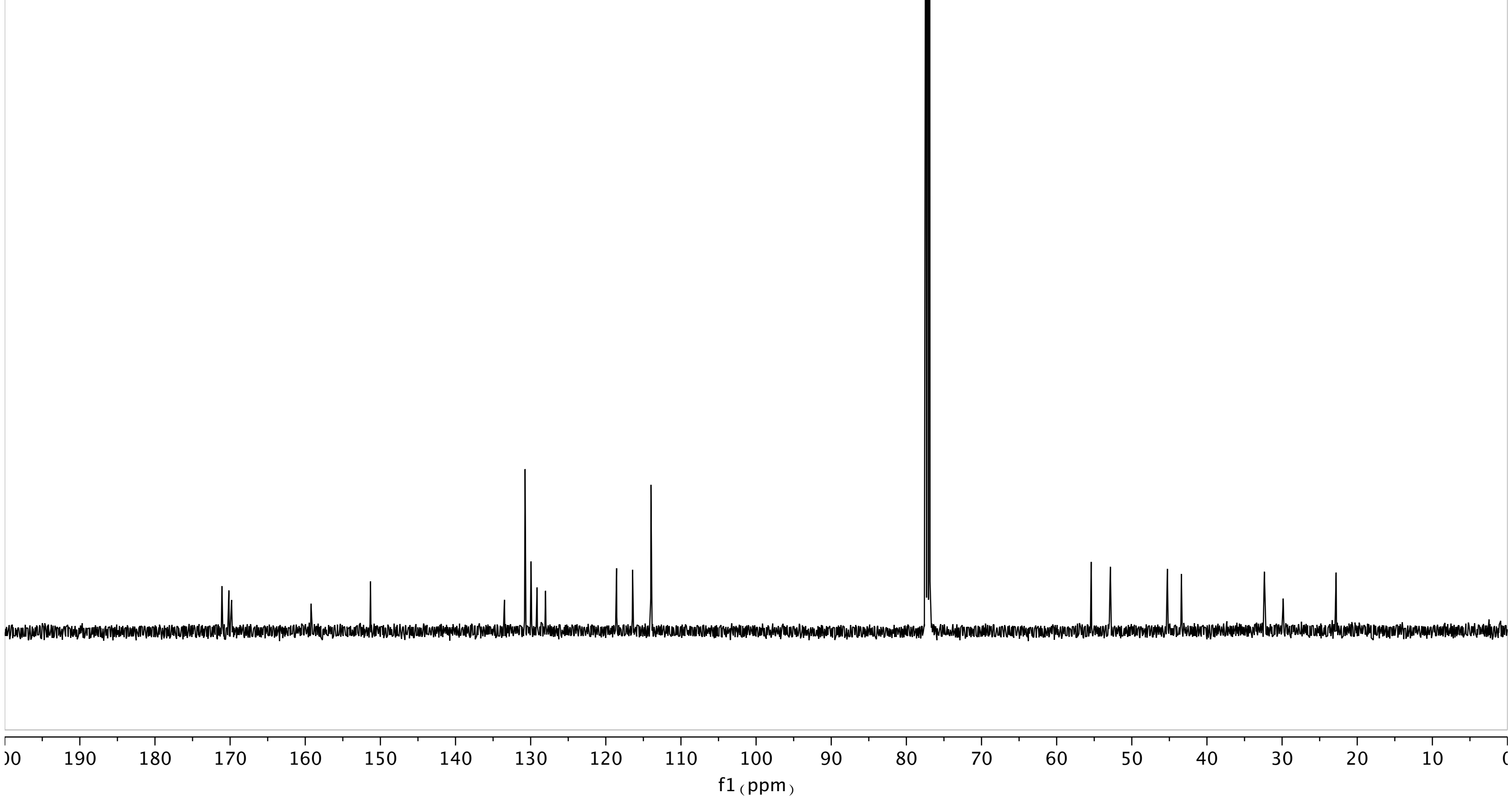

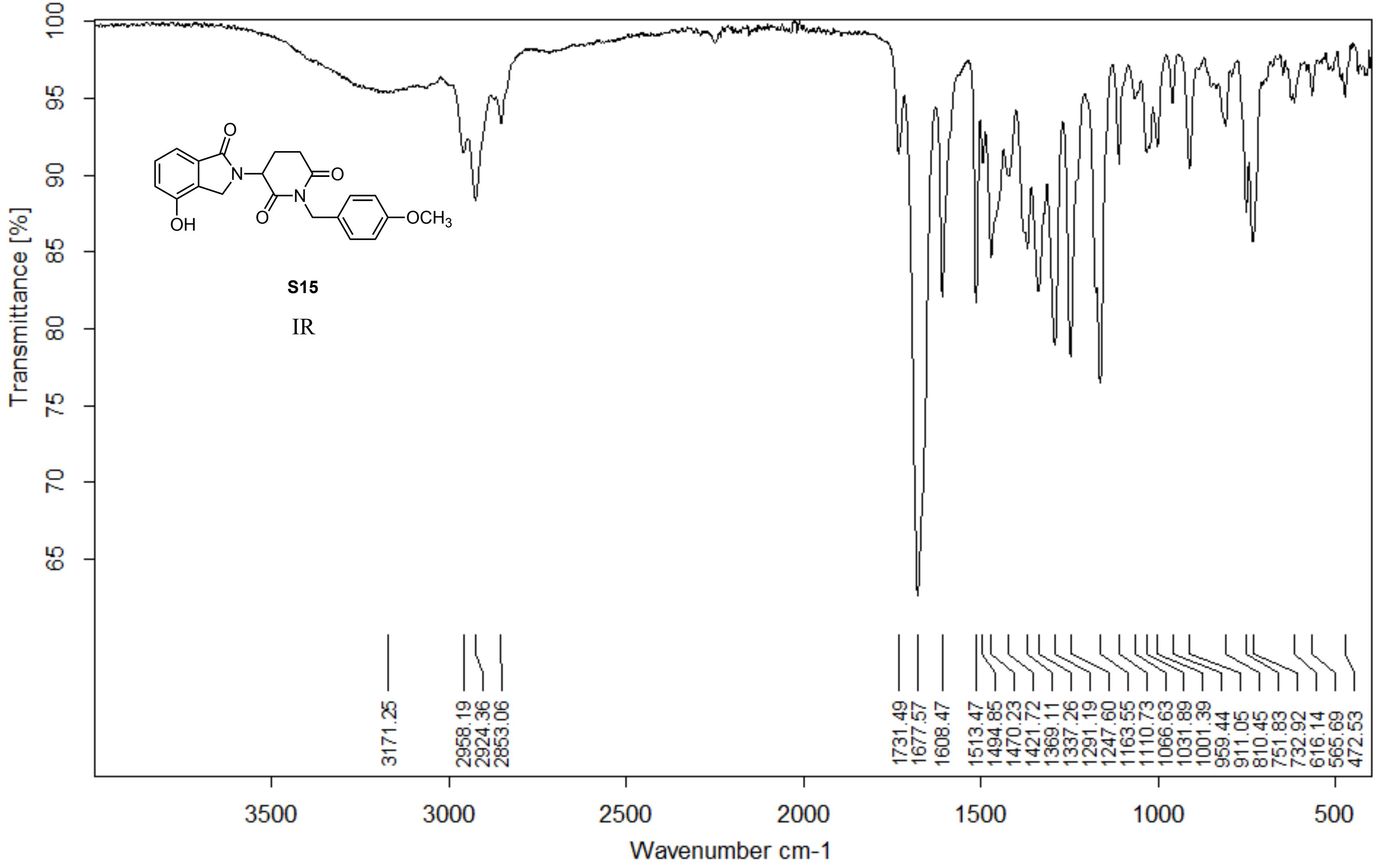


Z:\ALPHA FTIR DATA\Woo\Yuka\YA520.0 YA520 Diamond ATR

15/03/2020

**Page 1/2**

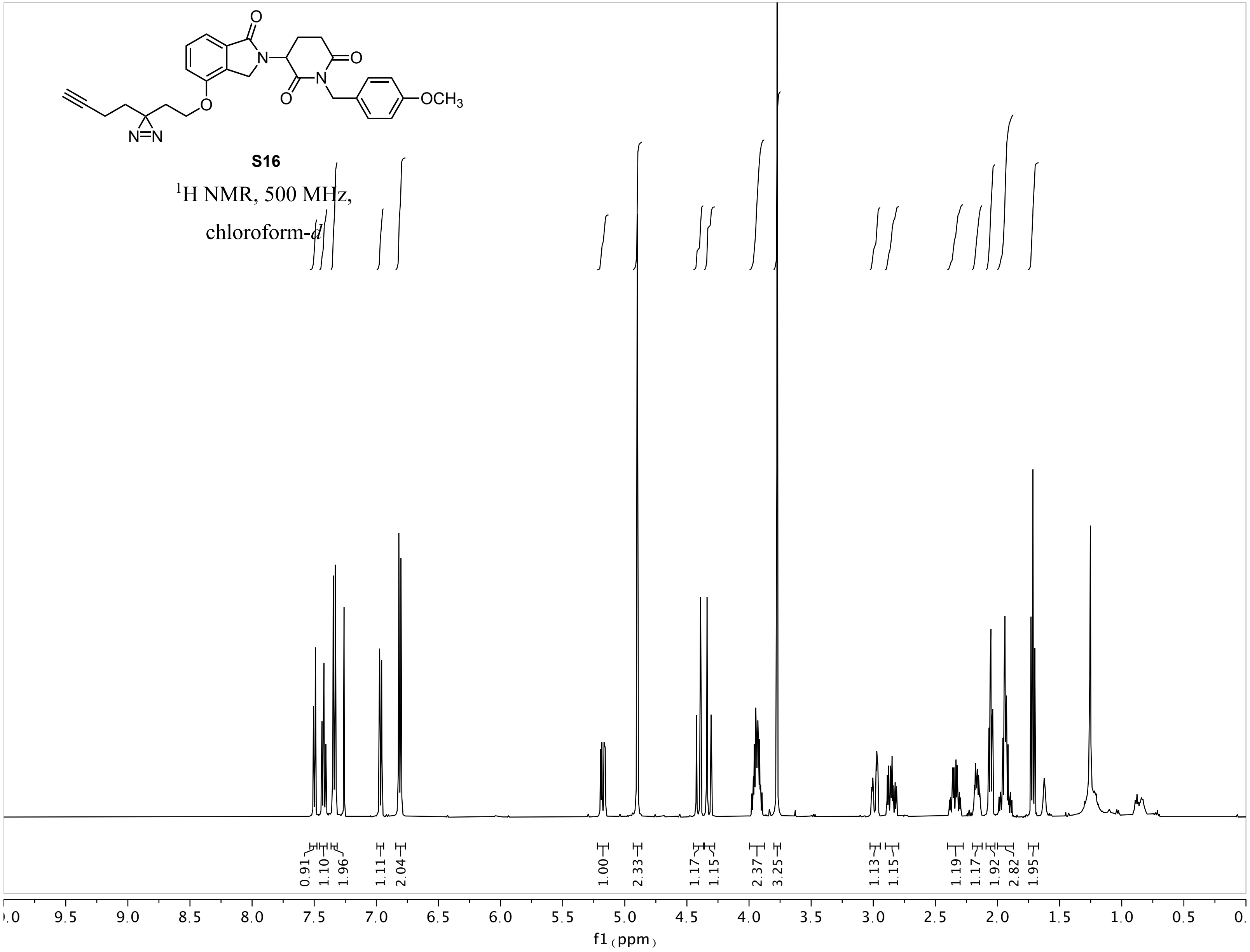

OCH3
N=N
S16
1H NMR, 500 MHz,
chloroform-d
0.91
1.10
1.96
1.11
2.04
1.00
2.33
1.17
1.15
2.37
3.25
1.13
1.15
1.19
1.17
1.92
2.82
1.95
9.5 9.0 8.5 8.0 7.5 7.0 6.5 6.0 5.5 5.0 4.5 4.0 3.5 3.0 2.5 2.0 1.5 1.0 0.5
f1 (ppm)

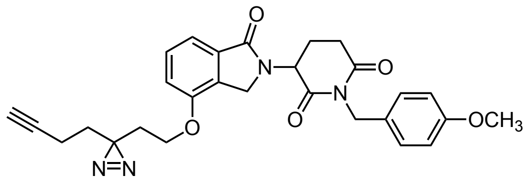


**S16**

$^{13}$C NMR, 125 MHz,

chloroform-*d*

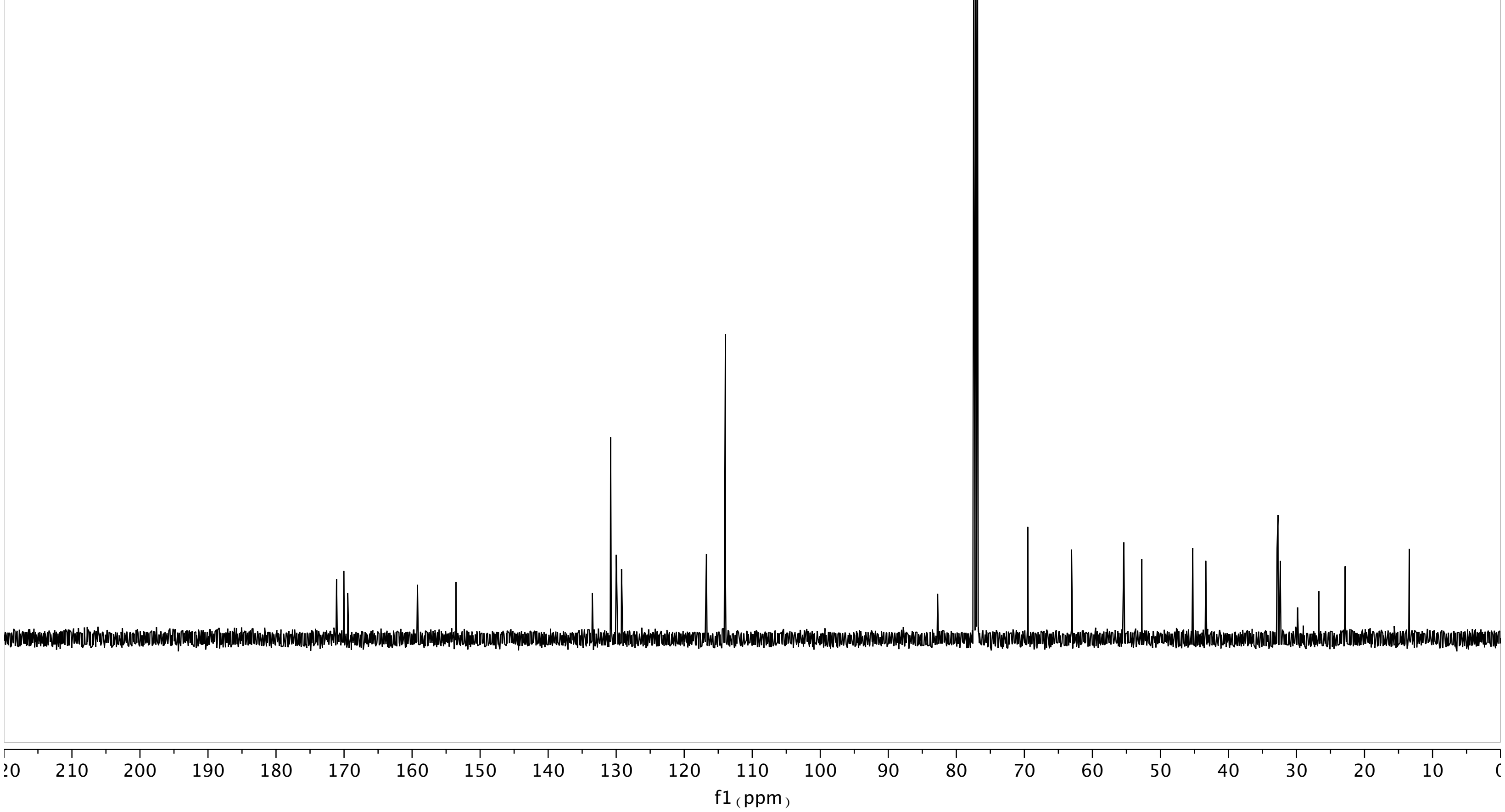

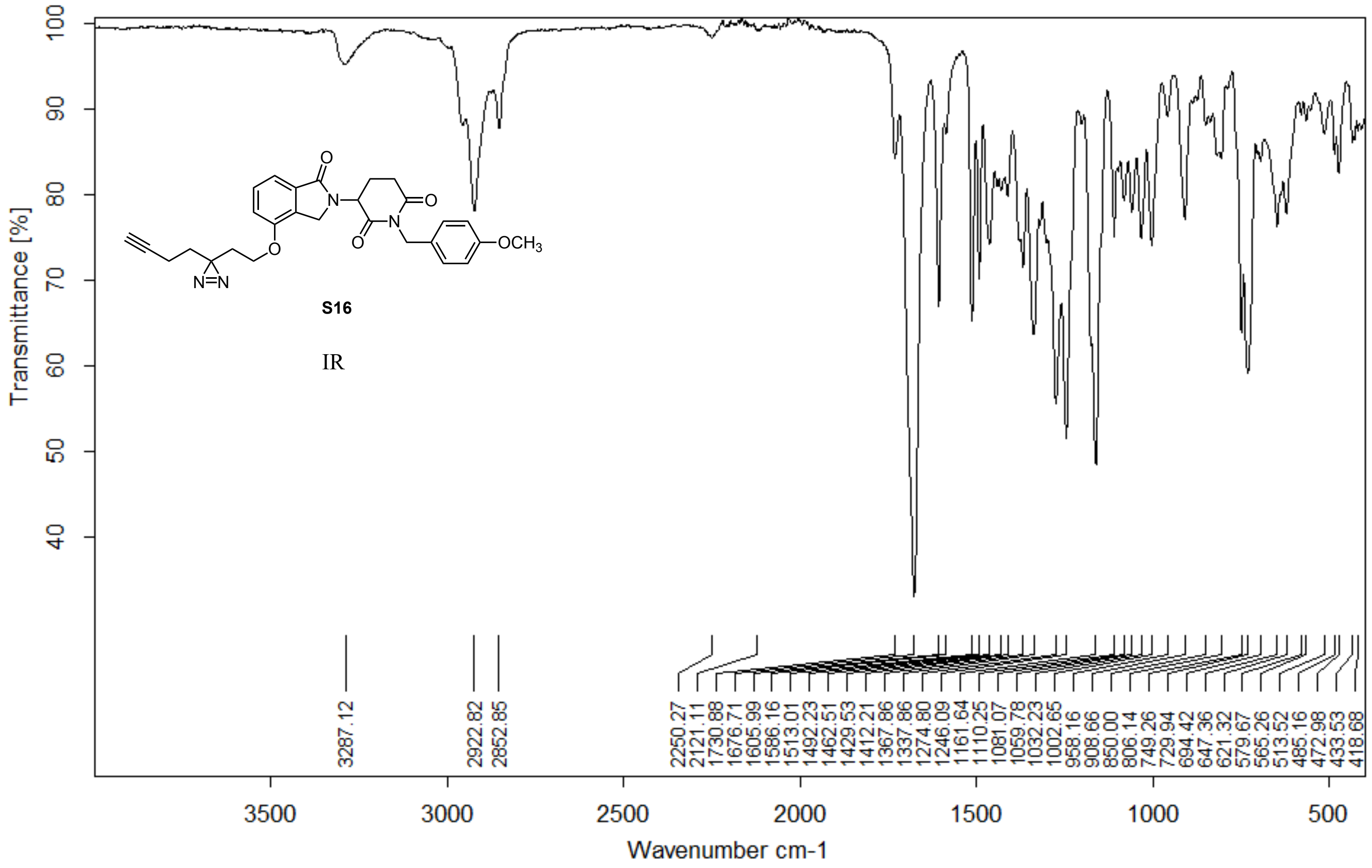


Z:\Alpha FTIR Data\Woo\Yuka\YA225-tm.0 YA225-tm Diamond ATR

13/01/2018

**Page 1/2**

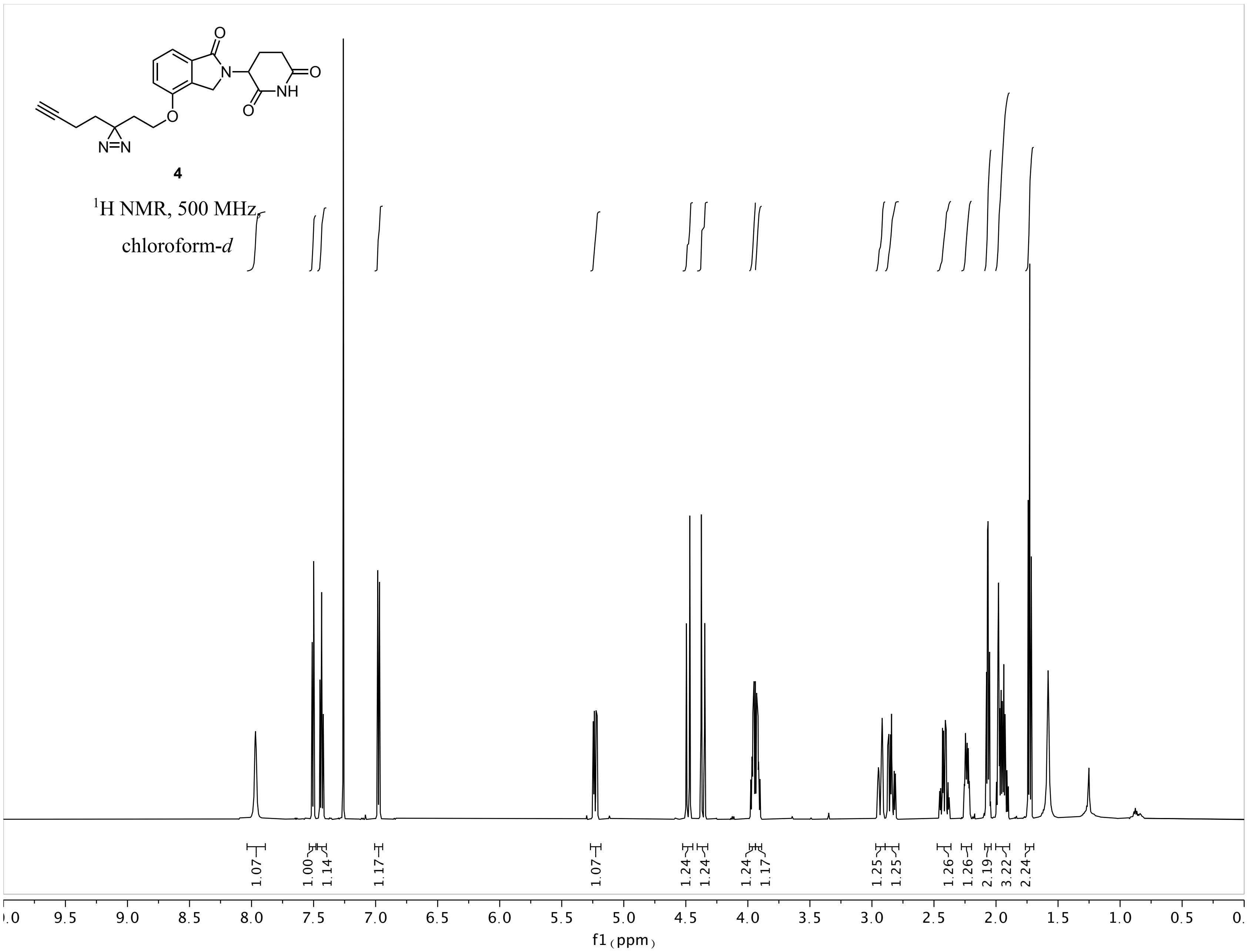
4
$^{1}$H NMR, 500 MHz,
chloroform-d
1.07
1.00
1.14
1.17
1.07
1.24
1.24
1.24
1.17
1.25
1.25
1.26
1.26
2.19
3.22
2.24
f1 (ppm)

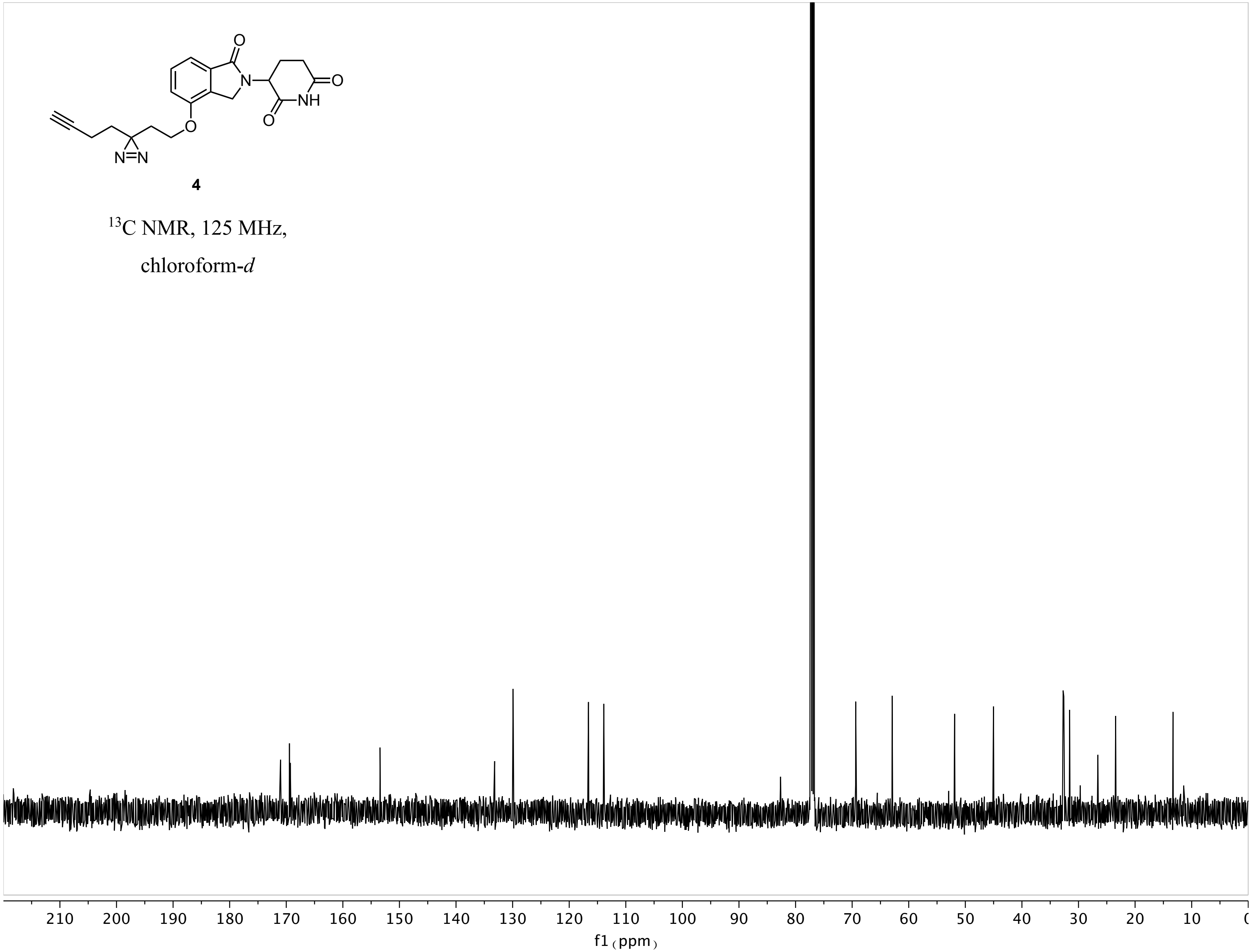
4
$^{13}$C NMR, 125 MHz,
chloroform-*d*
210 200 190 180 170 160 150 140 130 120 110 100 90 80 70 60 50 40 30 20 10
f1 (ppm)

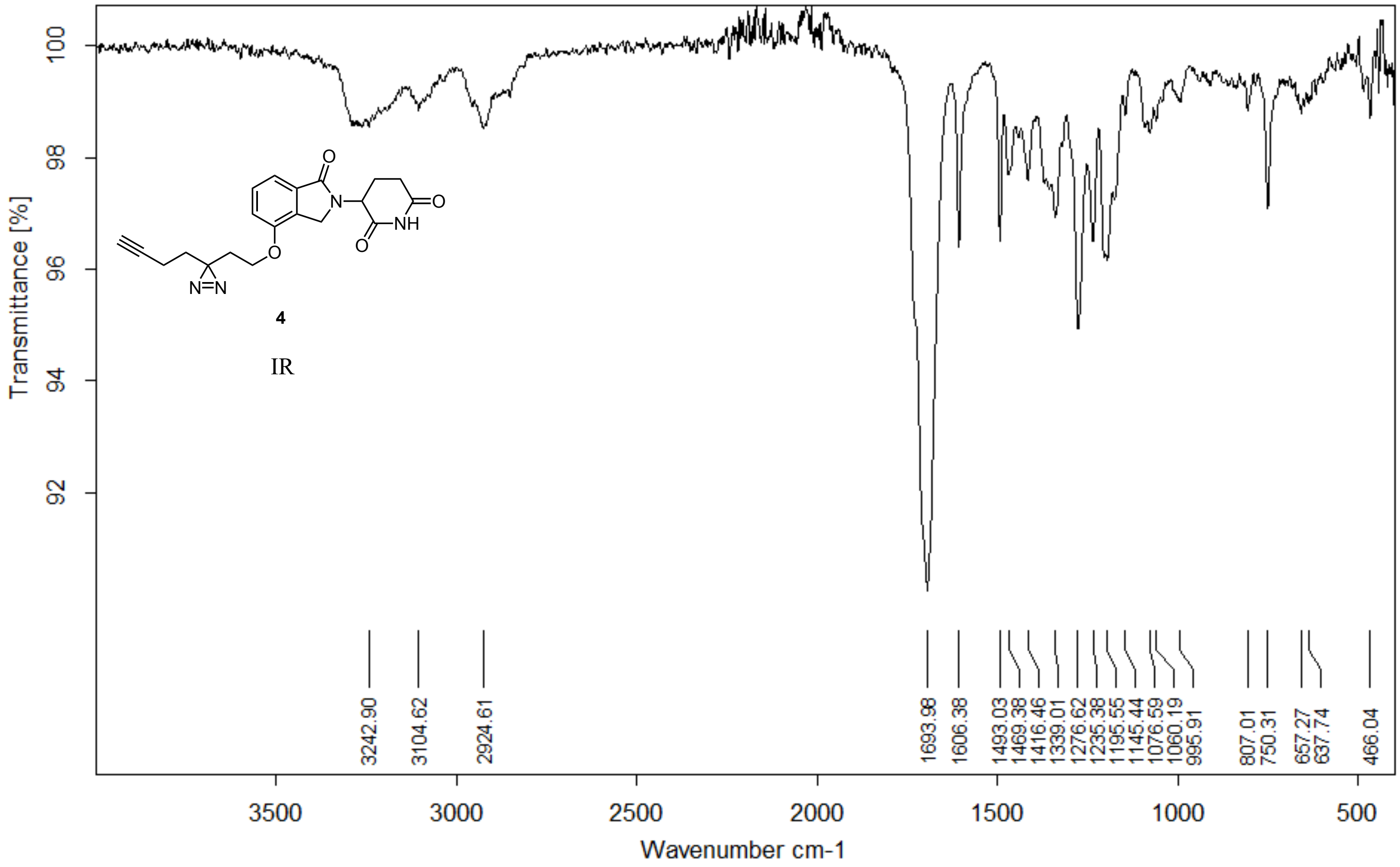


| | | | |
|---|---|---|---|
| Z:\Alpha FTIR Data\Woo\Yuka\YA227.0 | YA227 | Diamond ATR | 15/01/2018 |

**Page 1/2**

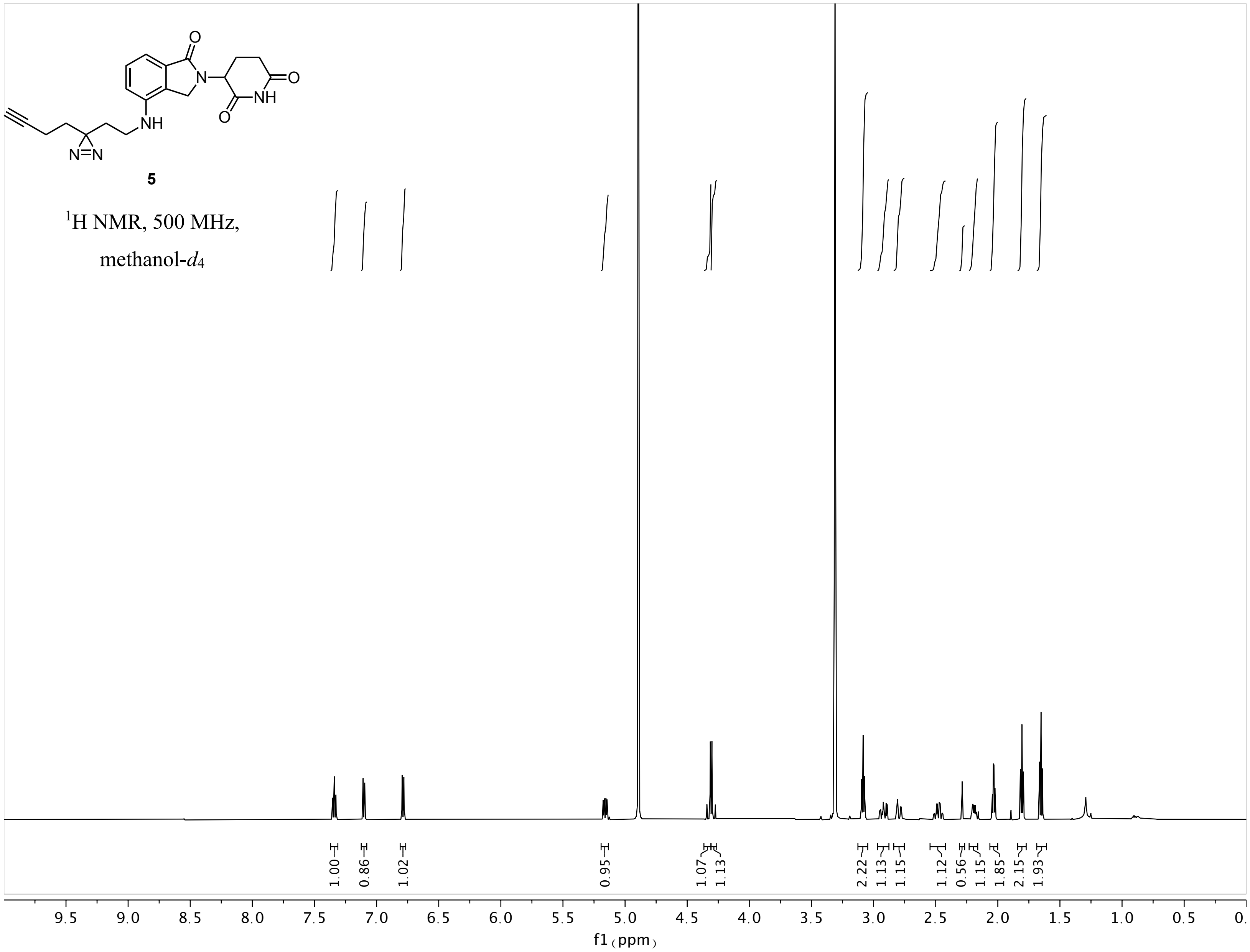
5
¹H NMR, 500 MHz,
methanol-d4
1.00
0.86
1.02
0.95
1.07
1.13
2.22
1.13
1.15
1.12
0.56
1.15
1.85
2.15
1.93
9.5
9.0
8.5
8.0
7.5
7.0
6.5
6.0
5.5
5.0
4.5
4.0
3.5
3.0
2.5
2.0
1.5
1.0
0.5
f1 (ppm)

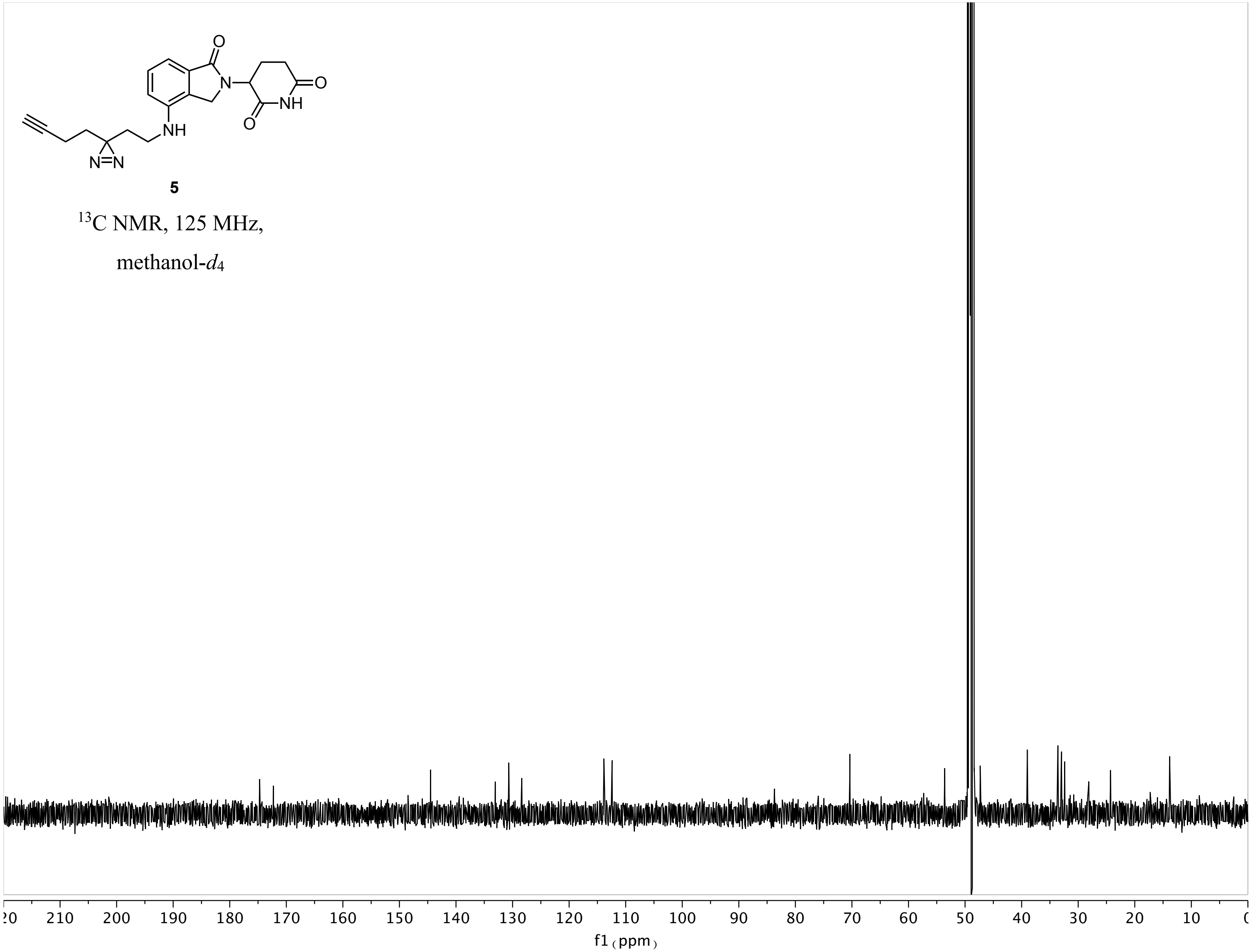

5
13C NMR, 125 MHz,
methanol-d4
f1 (ppm)

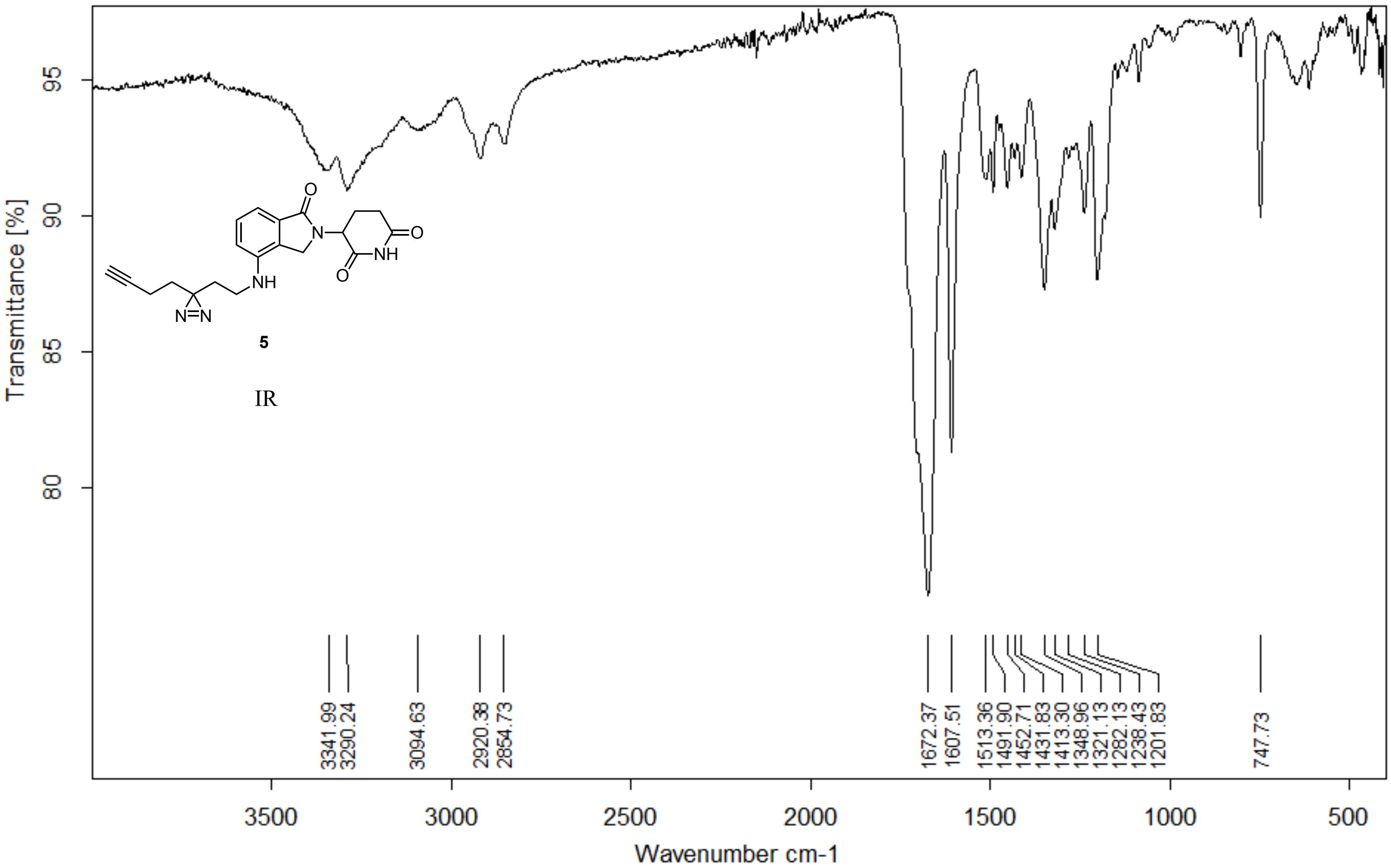


| | |
|---|---|
| Z:\Alpha FTIR Data\Woo\Yuka\YA241 NpLEN.2 YA241 NpLEN Diamond ATR | 09/06/2018 |

**Page 1/2**

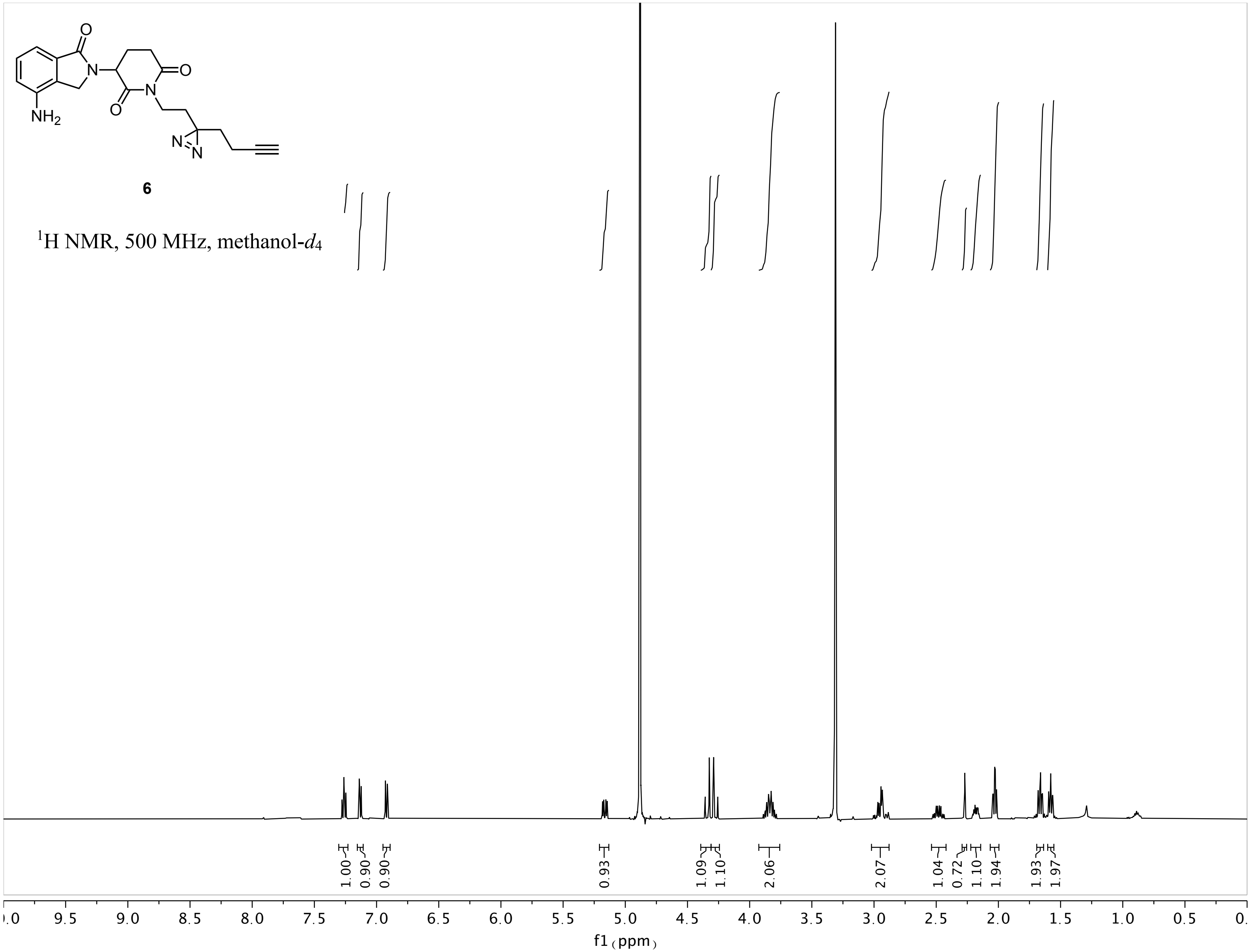
6
$^1$H NMR, 500 MHz, methanol-$d_4$
1.00
0.90
0.90
0.93
1.09
1.10
2.06
2.07
1.04
0.72
1.10
1.94
1.93
1.97
f1 (ppm)

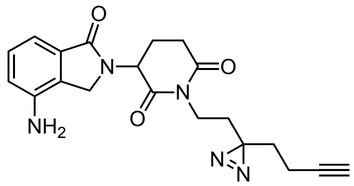


**6**

$^{13}$C NMR, 125 MHz, chloroform-*d*

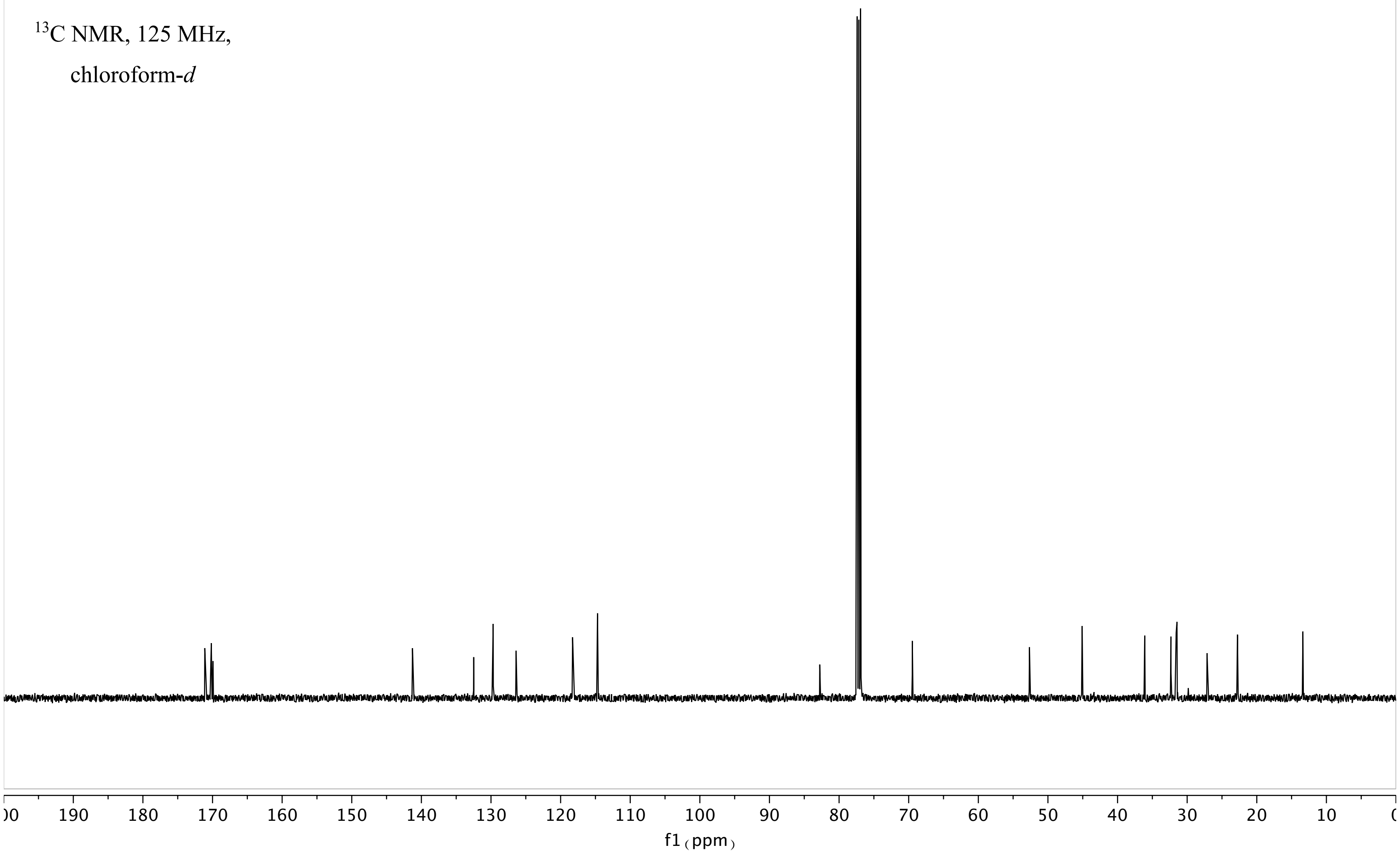

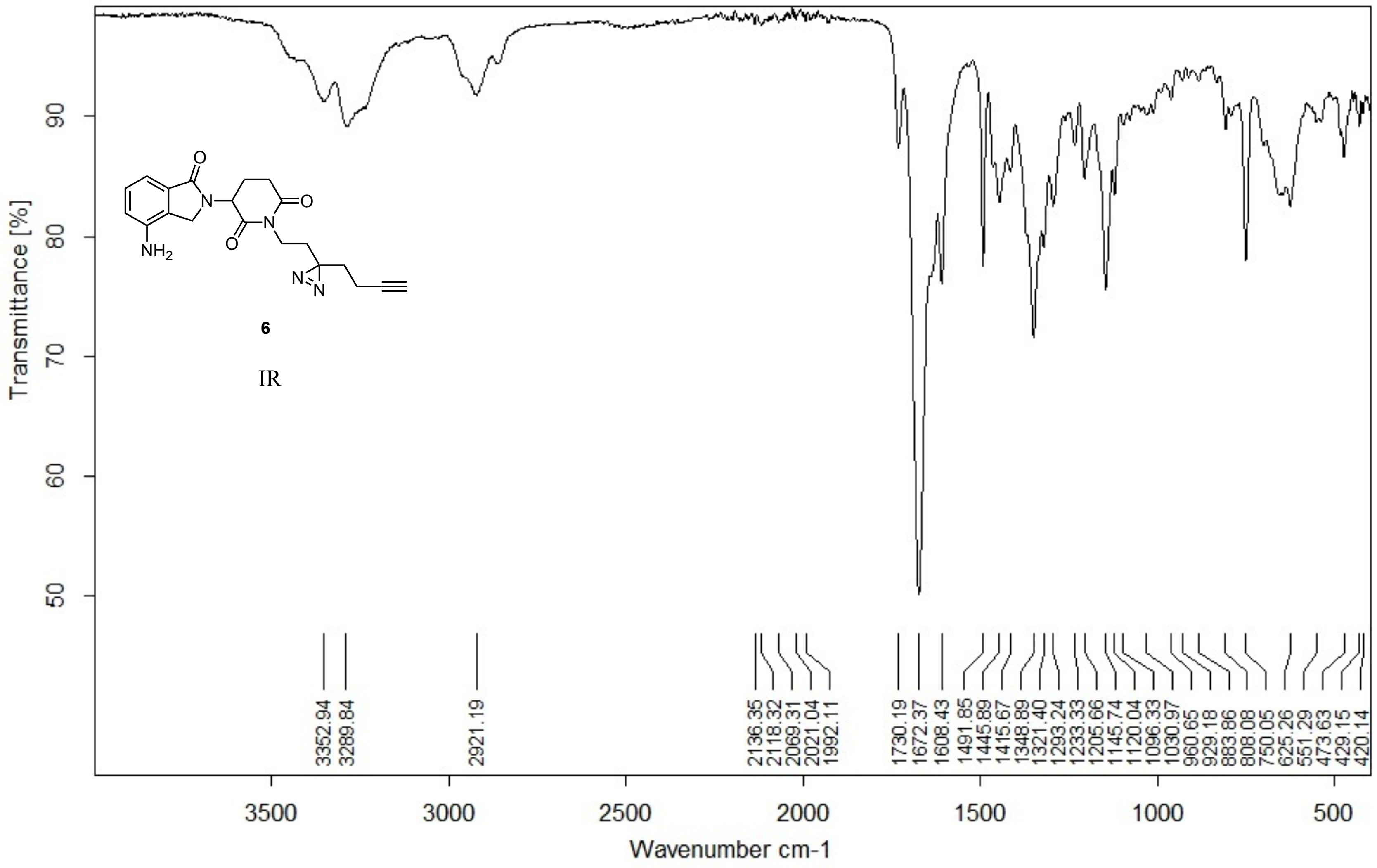


Z:\ALPHA FTIR DATA\Woo\Yuka\YA280.0 YA280 Diamond ATR

20/04/2018
